# A multi-omics atlas of human hindbrain development

**DOI:** 10.64898/2026.02.08.704646

**Authors:** Piyush Joshi, Mari Sepp, Ioannis Sarropoulos, Nils Trost, Konstantin Okonechnikov, Tetsuya Yamada, Céline Schneider, Julia Schmidt, Ashwyn A. Perera, Andrea Wittmann, Mirjam Blattner-Johnson, Barbara Jones, Cornelis M. van Tilburg, Olaf Witt, Steven Lisgo, Miklós Palkovits, David T.W. Jones, Supat Thongjuea, Henrik Kaessmann, Stefan M. Pfister, Lena M. Kutscher

**Affiliations:** Hopp Children’s Cancer Center (KiTZ), Heidelberg, Germany; Division of Pediatric Neurooncology, German Cancer Research Center (DKFZ) and German Cancer Consortium (DKTK), Heidelberg, Germany; Developmental Origins of Pediatric Cancer Junior Research Group, German Cancer Research Center (DKFZ), Heidelberg, Germany; National Center for Tumor Diseases (NCT), NCT Heidelberg, a partnership between DKFZ and Heidelberg University Hospital, Germany; Center for Molecular Biology of Heidelberg University (ZMBH), DKFZ-ZMBH Alliance, Heidelberg, Germany; Centre of Genomics, Evolution and Medicine (cGEM), Institute of Genomics, University of Tartu, Tartu, Estonia; Cambridge Stem Cell Institute and Department of Medicine, University of Cambridge, Cambridge, United Kingdom; Division of Pediatric Glioma Research (B360), German Cancer Research Center (DKFZ), Heidelberg, Germany; Department of Pediatric Oncology, Hematology & Immunology, Heidelberg University Hospital, Heidelberg, Germany; CCU Pediatric Oncology, German Cancer Research Center (DKFZ) and German Cancer Consortium (DKTK), Heidelberg, Germany; Newcastle University BioSciences Institute, Newcastle upon Tyne, United Kingdom; Semmelweis University, Budapest, Hungary

## Abstract

The human hindbrain controls essential motor and autonomic functions and is the site of neurodevelopmental diseases. Yet, its cellular diversity, developmental trajectories and underlying regulatory logic remain poorly understood. We present a comprehensive multi-omics atlas of human hindbrain development spanning embryonic to adult stages, encompassing 594,817 transcriptomic and 422,568 chromatin-accessibility single-nucleus profiles. This dataset resolved the cellular architecture of hindbrain cellular lineages, and delineated coordinated gene expression, cis-regulatory programs and regulatory grammar guiding their developmental trajectories. By integrating multi-omics data, we discovered context-specific roles of transcription factors across cell types and deciphered the role of HOX genes in driving divergent cellular identity in related lineages. We further leveraged the atlas to contextualize pediatric gliomas to decode how subtle yet coordinated shifts in gene expression context can define oncogenic transformation. Together, this atlas provides a foundational resource for hindbrain biology and establishes a gene-regulatory framework linking development and disease.

## Introduction

The hindbrain is a critical central nervous system region implicated in autonomic control^1,2^, motor coordination^3^, sensorimotor integration^4^, cognitive and linguistic processing^5^, and acts as a bridge between the forebrain and spinal cord^6-8^. During embryonic development, the hindbrain gets morphologically and molecularly segmented into rhombomeres (7/8 morphological segments, r1-8, r8 comprising four molecularly segmented crypto-rhombomeres r8-r11^9-11^), giving rise to isthmus at the midbrain-hindbrain boundary (r0), cerebellum (dorsal r1), pons (r2-r6) and medulla oblongata (r7-r11), with structure-specific cellular composition and functionality^3,12-15^. Despite these essential and functionally heterogeneous roles, it remains one of the least understood regions of the central nervous system, particularly due to a lack of mapping of the underlying cellular heterogeneity. Recent large-scale brain atlas initiatives have profiled millions of cells from forebrain, midbrain, and, to a limited extent, hindbrain, across developmental stages, generating valuable multimodal maps of mammalian systems^16-21^. Region-focused studies, including our groups’ cerebellum atlases^14,22,23^, have provided important insights into molecular mechanisms guiding development of this major hindbrain structure. Yet, these efforts have not achieved a comprehensive coverage or deep annotation across the complete human hindbrain, particularly in the pontine and medullary hindbrain regions, where sampling remains sparse and lineage relationships poorly resolved. This lack of knowledge is particularly significant given the region’s susceptibility to neurodevelopmental disorders^24,25^, and malignant transformation including brain tumors in children and adults^26^.

Here, we present an integrated single-nucleus multi-omics atlas of the developing human hindbrain, encompassing comprehensively annotated 594,817 transcriptomic and 422,568 chromatin accessibility single-nuclei profiles, spanning embryonic to adult stages. This atlas enables systematic reconstruction of neuronal and glial lineages, and provides a high-resolution view of the gene expression and underlying regulatory programs guiding their development. By combining single-modality analyses with integrative approaches, we resolved gene expression trajectories, delineated cis-regulatory landscapes, and inferred transcription factor activities underlying cellular identity. Leveraging these maps, we uncovered a regulatory grammar that organizes lineage fate decisions and deciphered divergent regulatory programs in the upper versus lower rhombic lip lineages. Finally, using the map of normal hindbrain development as a reference, we dissected pediatric glial tumor biology and revealed context drift in glial gene expression programs as a mechanism underlying oncogenic transformation. Together, these data and results establish a foundational resource for understanding human hindbrain development, its regulatory logic, and its vulnerability to disease. To facilitate further discoveries from this extensive resource, we provide user-friendly access to the processed data through the *HindbrainExplorer* (*https://apps.kaessmannlab.org/HindbrainExplorer/*) app and analysis scripts via github.

## RESULTS

### Developmental trajectories and cellular diversity revealed by a single-nucleus multi-omics atlas of the human hindbrain

To catalog cellular diversity in the developing human hindbrain, and resolve the underlying transcriptomic signatures, chromatin accessibility profiles and cis-regulatory elements (CREs), we processed hindbrain, pontine, and medullary tissues spanning embryonic to adult stages for single-nucleus RNA-sequencing (snRNA-seq) and single-nucleus ATAC-sequencing (snATAC-seq) using 10x Genomics-based 3’ single-cell transcriptomic and single-cell ATAC profiling technologies, respectively (Fig. 1A), and integrated data from previously published multi-omics cerebellar atlases^14,23^.

**Figure 1:**
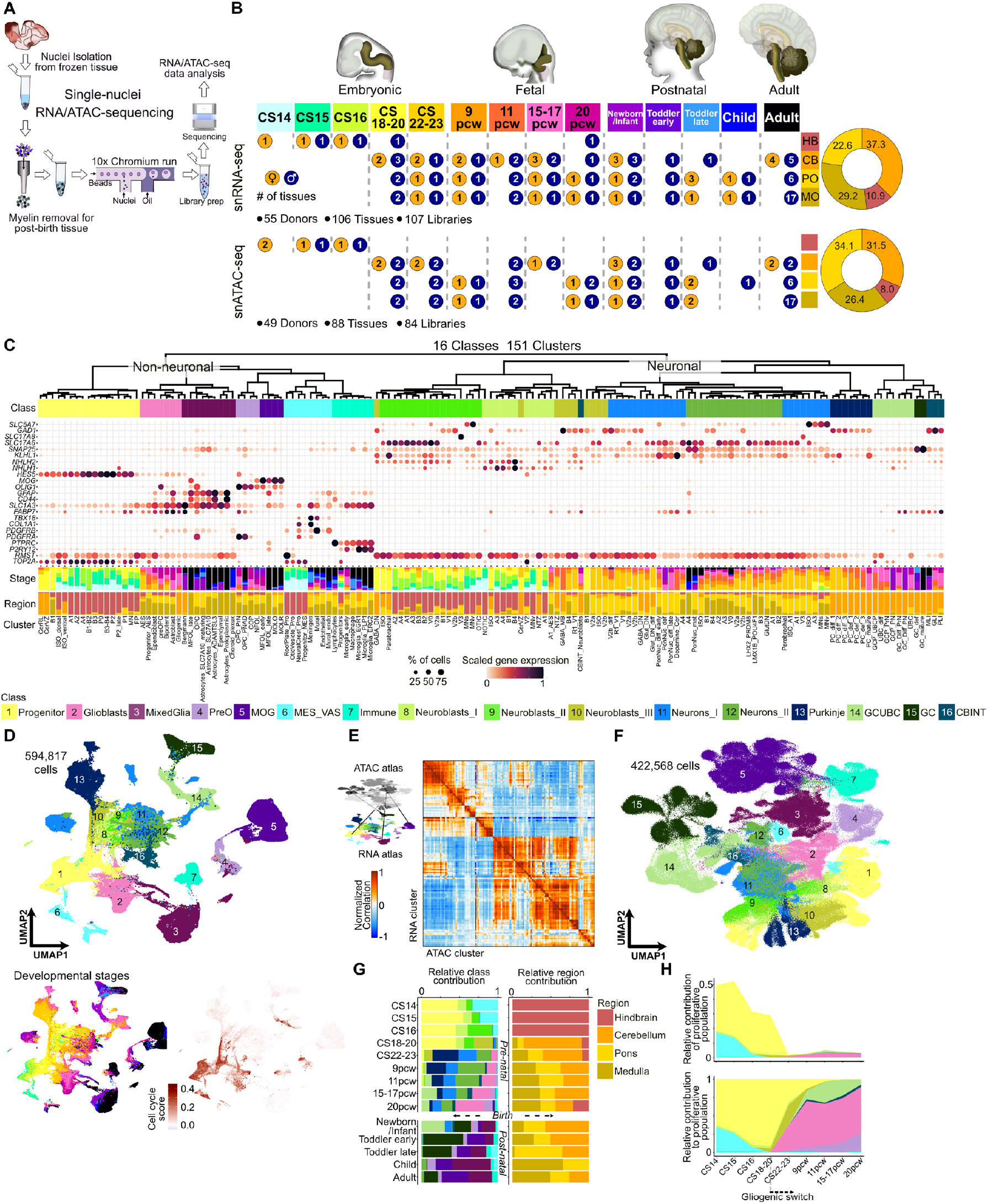
A multi-omics atlas of the developing human hindbrain. **A**) Schematic of single-nucleus RNA-seq and ATAC-seq experiment. **B**) Schematic of sample composition for the multi-omics atlas. Small circles depict number of male (blue) or female (orange) donors per stage per region (hindbrain (HB), cerebellum (CB), pons (PO) and medulla (MO)). HB represent undissected tissue encompassing CB, PO and MO regions. In adult, only selected CB, PO or MO regions were included instead of entire structure (Details in Table S1). Donot *legend cont. next page* ▸ plot depicts proportional contribution of regions to final annotated atlas per modality. **C**) Hierarchical clustering of annotated cell clusters and classes. Marker gene expression shown as a bubble plot. Bubble size, proportion of cells expressing. Bubble color, scaled expression. Bar plots depict proportional contribution of developmental stages and region of constituting cells per cluster. Uniform Manifold Approximation and Projection (UMAP) of all cells colored by cell-class identity (top), developmental stage (bottom, left) and cell cycle score (bottom right). **E**) Transfer of cell-cluster labels from the reference transcriptomic atlas to the chromatin accessibility atlas; correlation heatmap of pseudobulked gene scores (ATAC) and gene expression (RNA). **F**) UMAP of all cells colored by cell-class identity in the ATAC-seq atlas. **G**) Proportional distribution of cell classes (left) and regional contribution (right) across developmental time. Dashed line separates pre- and post-birth samples. Importantly, due to technical limitations of profiling large postnatal and adult hindbrain tissues, with depletion of oligodendrocyte populations to enrich other cell populations, the cellular composition in postnatal/adult stages is not reflective of actual cellular composition. **H**) Proportion of proliferating cells by stage, colored by cell class (top), and relative class contributions to the proliferating population (bottom). Gliogenic cell types appear between CS18-20.

For the transcriptomic atlas, we generated 107 libraries from 106 tissues sampled across 55 donors in total (Fig. 1B, Table S1). Quality control of individual libraries included removal of low-quality cells and empty droplets, correcting for background contamination, followed by doublet removal and filtering for outliers (see *Methods*; Table S2). Post-filtering, normalized gene expression data for 612,249 cells obtained from all libraries were embedded together using non-negative matrix factorization (NMF)-based dimensional reduction, which was used as input for graph-based clustering in a two-step process. Singletons, small cell clusters (<25 cells), and clusters that lacked definitive marker genes or exhibited low quality metric even after initial filtering, were removed from the atlas, resulting in a final set of 594,817 cells.

To generate complementary single-nucleus chromatin accessibility profiles, we generated 84 libraries from 88 tissues sampled from 49 donors derived from an overlapping subset of donors used for snRNA-seq (Fig. 1B, Table S1). We applied quality controls filtering low quality cells based on fraction of reads in peaks (FRiP) and transcription start site (TSS) enrichment metrics per library, followed by doublet removal per library (see *Methods*, Table S3). Post-filtering, 424,533 cells derived from snATAC-seq data across all samples were embedded together using latent semantic indexing (LSI)-based dimension reduction, clustered using a graph-based approach in a two-step process to obtain a joint representation of the atlas. Removal of small cell clusters and clusters with non-informative profiles resulted in 422,850 cells across human hindbrain development. For both transcriptomic and chromatin accessibility profiles, novel data represented around two-thirds of the integrated atlas (Fig. 1B).

Next, we annotated cells in the transcriptomic dataset based on expression of marker genes obtained from extensive literature-based evidence, developmental stage, and inferred differentiation state. We defined 151 clusters and 375 subclusters (Fig. 1C; Fig. S1A,B; Table S4). To provide an overview of cell types comprising the hindbrain, we grouped the 151 clusters into 16 classes that broadly separated into neuronal and non-neuronal lineages (Fig. 1C,D). The non-neuronal classes included radial glial progenitors (Progenitors), multipotent glial progenitors and committed glial precursors (Glioblasts), astrocytic and ependymal derivatives (MixedGlia), pre- and post-natal oligodendrocyte progenitors (PreO), mature oligodendrocytes (MOG), neural crest-derived mesenchymal cells and vascular/endothelial cells (MES_ VAS), and immune cells comprised mostly of microglia (Immune). Neuronal classes include early and late differentiating neuroblasts (Neuroblasts_I, Neuroblasts_ II; Neuroblasts_I with relatively higher cell-cycle score and Neuroblast_II representing more mature neuroblasts; Fig S1C,D), neuroblasts dominated by cerebellar contribution (Neuroblasts_III), proliferating and differentiating granule/unipolar brush cells (GCUBC), early and late differentiated neurons (Neurons_I, Neurons_II; Neurons_ II with relatively higher contribution from adult tissues; Fig S1E), differentiated granule cells (GC), cerebellar interneurons (CBINT), and Purkinje cells (Purkinje). Importantly, neuroblasts and neurons obtained from the cerebellum versus pons/medulla were molecularly distinct; for example, cerebellar neurons including Purkinje, CBINT and GC formed separate classes compared with Neurons_I and Neurons_II representing pontine/ medullary neurons (Fig. 1C). Additionally, pontine and medullary neuronal populations clustered together at class level, even though they represented a heterogenous mix of glutamatergic and GABAergic cell lineages, with some noradrenergic, serotonergic and cholinergic populations included, a pattern similar to ‘splatter neurons’ described by Siletti *et al*^17^. In contrast to neuronal populations, glial populations from all hindbrain regions, including the cerebellum, were grouped together at the class level (Fig. 1C), in line with previous observations^17^, suggesting higher molecular similarity among axially distinct but similar glial populations with little influence of abutting neuronal populations on glial molecular programs.

Next, to characterize chromatin accessibility profiles across annotated clusters, we used the transcriptomic atlas as a reference to annotate the integrated ATAC-seq atlas. Cluster labels were transferred based on canonical correlation analysis^27^ between gene expression profiles from the transcriptomic atlas and predicted gene expression (gene score values) derived from chromatin accessibility in the snATAC-seq atlas. This integration yielded chromatin accessibility states for 132 of the 151 clusters identified in the transcriptomic atlas, each represented by at least 50 cells in the snATAC-seq dataset, totaling 422,568 cells (Fig. S2). The predicted gene scores for individual clusters showed the strongest correlation with their corresponding reference clusters from the transcriptomic atlas (Fig 1E; Fig. S3A-P). Accordingly, these clusters were assigned to the same cell classes as in the transcriptomic atlas (Fig 1F; Fig. S2).

We next evaluated stage-specific class contributions, which showed that progenitors were derived from embryonic stages, neuroblasts and glioblasts peaked in late embryonic to early fetal stages, and differentiated neurons and glia predominated in late fetal and postnatal stages (Fig. 1G; Fig S1E). Notably, by Carnegie stages 22-23, radial glial progenitors were nearly depleted, with residual proliferating populations predominantly consisting of gliogenic progenitors, first detected around Carnegie stages 18-20 (or around post-conception week 7), marking the gliogenic switch in the hindbrain, aligning with a previous observation^18^ (Fig. 1H). Notably, this gliogenic switch is much earlier than the reported gliogenic switches in the cortex or the midbrain, which occur around post-conception week 10.5 and 9.5, respectively^18^. Post birth, oligodendrocytes progenitors (OPC_PNAD) and granule cell progenitors (GCP_PN) represented the dominant proliferating population in the newborn (Fig. S1F). Interestingly, *PDGFRA*+ post-natal and adult oligodendrocyte progenitors (OPC_PNAD), while presumptively representing a progenitor population, based on their gene expression similarity to prenatal oligodendrocyte progenitors (OPC_PrN), exhibit very low cell cycle activity (Fig. S1G), suggesting a quiescent state. How these molecularly similar pre- and post-birth progenitor cell types exhibit drastically different proliferative capacity requires further investigation.

In summary, our integrated atlas provides a comprehensive multi-omics overview of cellular diversity in the developing human hindbrain, identifying major cell classes and key cellular events.

### Generation of neuronal diversity in the pons and medulla

Neuronal diversity in the vertebrate hindbrain is known to arise from integrated activity of orthogonal signaling factor gradients along the dorso-ventral (DV) and anterior-posterior (AP) axes^28-32^. These signaling factor gradients set up a transcription factor (TF) code, where *PAX* and *LHX* genes define DV identity^13,33,34^ and *HOX* genes define AP identity^30,35^. As a result, the hindbrain is patterned along the DV axis into dorsal/alar (A1-A4), intermediate (B1-B4), and ventral/basal (0-3, MN) domains (Fig. 2A) and along the AP axis into rhombomeres (r1-r11). In broad terms, DV patterning biases neurotransmitter identity, while AP patterning refines neuronal fates and circuit integration^7,28,33^.

**Figure 2:**
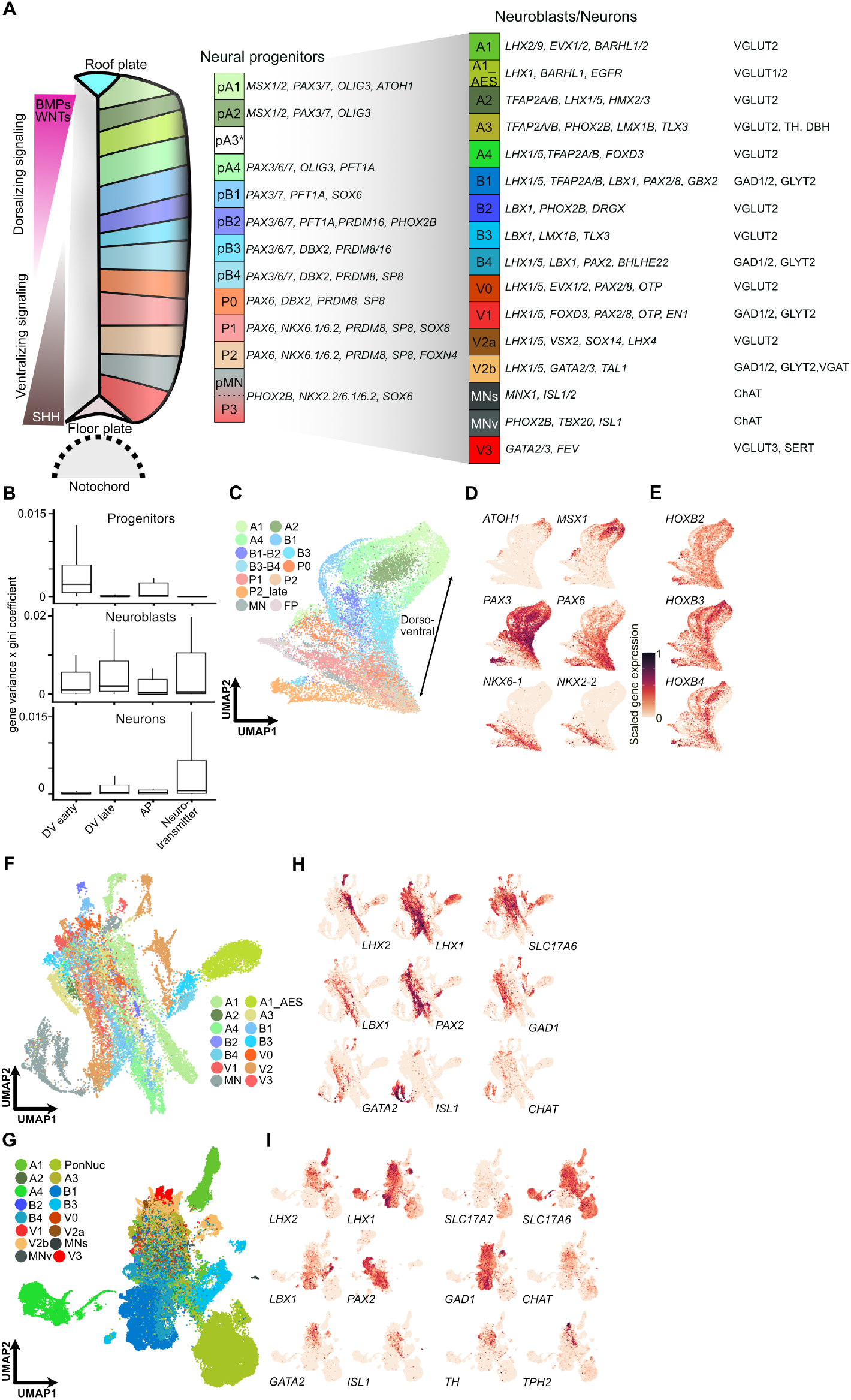
Neuronal diversity in the pons and medulla. **A**) Schematic of neuronal lineages in pons and medulla shown as clusters grouped by lineage from progenitors to mature neurons. Rectangle represent domains, defined by marker genes. Neurons are identified as glutamatergic (*SLC17A7/6/8*: VGLUT1/2/2), GABAergic (GAD1/2), glycinergic (*SLC6A5*: GLYT2), noradrenergic (TH and DBH) cholinergic (ChAT) or serotonergic (*SLC6A4*: SERT and TPH2) with appropriate neurotransmitter markers. **B**) Boxplot distribution of product of gene’s variance and gini coefficient for early DV, late DV, AP and neurotransmitter gene sets (Table S5) in pontine/medullary progenitor, neuroblasts and neuron populations. **C**) UMAP embedding of annotated DV progenitor domains. Arrow highlights the observed DV axis. **D**,**E**)UMAP-expression plot of selected early DV (**D**) and AP (**E**) markers in progenitor population. **F**,**G**) UMAP embedding of annotated pontine/medullary DV neuroblasts (**F**) and neuron (**G**) clusters. Multiple clusters representing the same DV domains in distinct classes of neuroblasts (Neuroblasts_I, Neuroblasts_II, Neuroblasts_III) and neurons (Neurons_I, Neurons_II) were grouped together. **H**,**I**) UMAP-expression plot of selected late DV and neurotransmitter markers in neuroblasts (**H**) and neuron (**I**) population.

To examine the neuronal lineages in pontine/medulla in detail, we removed cerebellar cells from the analysis and focused separately on progenitor, neuroblast, and neuron populations. We observed that DV markers showed higher expression and specificity (product of expression variance and gini-index) than AP markers across clusters of these cell populations (Fig. 2B). Expression of early DV markers was more cluster-specific in progenitors compared to that of late DV markers, which resolved clusters in neuroblasts and neurons (Fig. 2B-I, Table S5). In neuroblasts, and even more so in neurons, expression of neurotransmitters further defined cluster identity (Fig. 2B,H,I). As in the integrated UMAP embedding (Fig. 1D), UMAP embedding of the pontine/medullary neurons only (Neurons_I and II) resulted in a scattered, salt-and-pepper-like distribution of neuronal populations, consistent with ‘splatter’ patterning observed by Siletti *et al*.^17^; probably driven by lower expression of cluster-defining specific DV markers and inability of mature neuron markers, including neurotransmitter genes, to properly resolve cell populations in a 2-dimensional embedding (Fig. 2B,G). Altogether, DV markers constituted the strongest transcriptional signals delineating neuronal trajectories from progenitors to neurons, and became increasingly intermingled with neurotransmitter signatures as neurons matured.

Using markers for DV progenitor domains, including *PAX, DBX, NKX*, we identified progenitor clusters representing the dorsal (pA1-2,4), intermediate (pB1-4), and ventral progenitor domains (P0-2, pMN); however, no distinct cluster corresponding to the pA3 domain was resolved, and the P3 (marked by *ARX, NKX2-2*) population potentially clustered together with pMN (Fig. 2A,C-D; Fig. S4A-G). To define the DV neuroblast/neuron populations, we used the established domain markers, such as *LHX, LBX1, PAX*, and identified dorsal (A1-A4, including migratory A1_AES giving rise to pontine nuclei), intermediate (B1-B4) and ventral (V0-V3, MNv, MNs) neuronal clusters (Fig. 2A,F,G; Fig S4A-G). Finally, we used the expression of genes associated with the neurotransmitter phenotype to classify neurons into glutamatergic (*SLC17A7/6/8*; VGLUT1-3), GABAergic (*GAD1/2*), glycinergic (*SLC6A5*; GLYT2), cholinergic (CHAT), noradrenergic (TH and DBH), and serotonergic (*SLC6A4*: SERT and TPH2) neuronal populations (Fig. 2A,H,I; Fig. S4A-G). These classifications are consistent with prior knowledge^33^ and the overall dorsoventral gradient, progressing from dorsal glutamatergic populations through mixed and inhibitory domains to ventral cholinergic and serotonergic neuron types.

Next, we performed differential gene expression analysis to identify novel markers across the progenitor and neuronal clusters. We did not identify any previously uninvestigated TF markers, but uncovered a novel set of long noncoding marker RNAs (Fig. S5A,B). These included anti-sense transcripts to known TF marker genes such as *LHX1-DT* and *PHOX2B-AS1*, potentially driven by transcription from a common promoter, in addition to other potentially important microRNA host genes such as *MIR124*−*1HG* (*mir124-1*) and *MIRLET7BHG* (*mirlet7*). This expands the repertoire of molecular markers available for accessing the hindbrain cell populations.

Integration of our cerebellar atlas with data from other hindbrain regions reaffirmed our previous annotations for cerebellar cells^14^ (Table S4), although the dataset was reannotated using the same framework applied to the pons and medulla. This integration nonetheless provided novel insights into potential migratory behaviors of neurons originating from the cerebellar primordium. Specifically, we identified populations similar to unipolar brush cells (UBCs) and granule cells in the pontine and medullary samples (Fig. S6A,B). Importantly, these populations lacked significant expression of *HOX* genes, as expected in cerebellar populations that do not express any *HOX* genes, which are abundantly expressed in all pontine/medullary neural populations^35^, suggesting potential posterior migration of *HOX*-negative, upper rhombic lip-derived neuroblasts during early hindbrain development. Similarly, we identified cerebellar ventricular zone-derived interneuron-like (Fig. S6C,D) and relatively less numerous Purkinje cell-like (Fig. S6E,F) populations in pons or medulla tissues. These observations match pontine/medullary contributions to upper rhombic lip and cerebellar inhibitory superclusters described by Siletti *et al*^17^. While extracerebellar migration of upper rhombic lip-derived populations is documented^36,37^, the localization and fate of these cells, as well as of possible cerebellar ventricular zone-derived neurons, within the pons and medulla warrant further investigation.

### Coordinated transcriptomic programs shaping hindbrain cellular diversity

Having cataloged the cellular diversity in the developing human hindbrain, we next aimed to resolve the coordinated gene expression programs underlying this diversity. While differentially expressed genes are essential for cellular annotation, they fail to fully capture underlying coordinated and biologically relevant transcriptomic programs. This gap could be filled by dimensional reduction approaches, such as NMF or latent Dirichlet allocation (LDA)^38,39^. Therefore, we leveraged an LDA-based approach using a two-step process. First, we applied multi-ranked LDA factorization to gene expression matrices per class to obtain representative gene sets per latent factor across ranks. Then, we clustered the obtained gene sets across all the classes based on their overlap, to obtain a representative gene set, or meta-gene program, per gene set cluster (Fig. 3A, Table S6).

**Figure 3:**
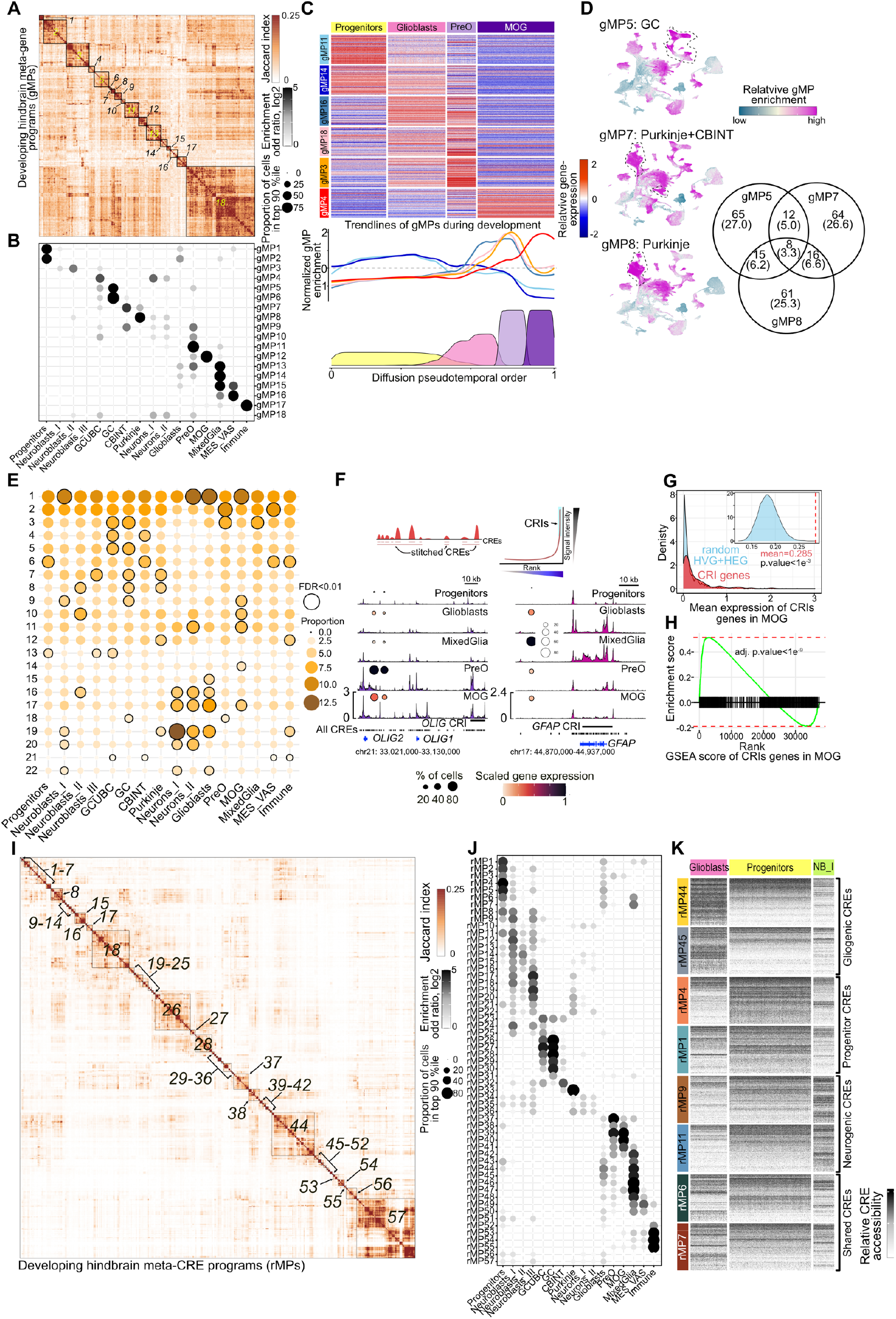
Transcriptional and cis-regulatory programs driving human hindbrain development. **A**) Heatmap of meta-gene programs (gMPs) derived from coordinated gene expression. Boxes highlight clustered gene set contributing to a meta-program. **B**) Enrichment analysis for class-specific gMPs shown as a bubble plot. Bubble size, proportion of cells in the top 90% ranked by gene set activity score. Bubble color, log2 enrichment odds ratio. **C**) Heatmap of gMPs underlying oligodendrocyte development (top), spanning progenitors (gMP11, 14), glioblasts (gMP16, 17), pre-oligodendrocytes (PreO; gMP3) and mature oligodendrocytes (MOG; gMP4). Trendline of scaled gMP enrichment along pseudotime (middle). Cell density plot along the pseudotime trajectory with cells colored by class (bottom). **D**) Scaled enrichment of gMPs associated with neuronal differentiation in mature granule neurons (GC; gMP5), Purkinje cells and cerebellar interneurons (Purkinje +CBINT; gMP7), and Purkinje cells (gMP7,8). Venn diagram depicting overlap among constituting genes. Number in parentheses represent percentage. **E**) Enrichment of top 10,000 (maximum) marker cis-regulatory elements (CREs) per class per chromosome. Bubble size, proportion of marker CREs per chromosome. Circled bubble with adjust p-value < 0.05, Chi-squared test. **F**) Schematic of cis-regulatory islands (CRIs, top). Chromatin accessibility profiles at the *OLIG1/2* (bottom left) and *GFAP* (bottom right) loci with associated cis-regulatory islands highlighted, showing lineage-specific enrichment. Bubbles, expression of *OLIG1/2* and *GFAP*. Bubble size, proportion of cells. Color, scaled expression. **G**) Density plots of expression for randomly selected set of highly expressed and highly variable genes in mature oligodendrocytes (MOG, blue) and genes located in topologically associated domains around MOG associated top 100 CRIs (left). Inset shows mean values of 10,000 random permutation and red dashed line the mean of CRIs associated gene set. **H**) GSEA of genes associated with top 100 CRIs in MOG in the marker genes (pair-wise Wilcoxon test) for the MOG cell population. **I**) Heatmap of robust meta-regulatory programs (rMPs) identified from co-accessible chromatin accessibility profiles. Boxes highlight CRE-sets comprising a meta-program. **J**) Enrichment of rMPs across cell classes shown as a bubble plot. Bubble size, proportion of cells in the top 90% ranked by CRE-activity score. Color, log2 enrichment odds-ratio. **K**) Enrichment of CREs associated with class-specific rMPs involved in gliogenic versus neurogenic fates. Columns, pseudobulked groups of 50 cells. Rows, 200 randomly selected CREs per meta-program. CRE accessibility scaled across clusters.

We next evaluated individual meta-gene programs by examining their contributing genes and biological significance, based on overrepresented gene ontology terms (biological process and annotated reactome pathways; Fig. 3B; Fig. S7A-D). We also quantified their activity using gene set module scores across classes and clusters (Fig. 3B; Fig. S7A). This analysis identified both confirmatory and novel insights. Among the confirmatory results was cell cycle associated meta-gene program (gMP1), which included *TOP2A* and *MKI67* in the representative set, and exhibited high enrichments in Progenitors and slight enrichments in other classes containing proliferating cells, such as Glioblasts, Neuroblasts_I and MES_VAS. Mature oligodendrocytes associated meta-gene program (gMP12) was associated with cholesterol biosynthesis, a known feature of differentiated oligodendrocytes^40^, and included differentiated oligodendrocytes markers *PLP1* and *MOBP*. We also identified a housekeeping meta-gene program (gMP3) associated with ‘homeostatic processes’, comprising of known housekeeping genes such as *GAPDH, ACTG1* and *ACTB*, and derived from gene sets contributed by multiple classes (13 out of 16 classes; Fig. S7B).

Novel insights included meta-gene program (gMP10) shared between proliferating and differentiating granule cells (GCUBC) and oligodendrocyte progenitors (PreO), highlighting shared expression programs between these two distinct lineages (Fig. 3B). In addition to shared housekeeping meta-gene program (gMP3), we identified another shared meta-gene program (gMP18) derived from multiple class contributions (12 out of 16 classes, excluding mature glial classes MixedGlia and MOG, and non-neural classes Immune and MES_VAS). Gene ontology analysis for this gene set did not identify any housekeeping terms, but was instead enriched for generic neural biology, including ‘neural system’. This finding suggests a presence of shared non-housekeeping, but basal biology, between the neural cells. While delineating cell-type-specific or shared coordinated expression programs, meta-gene programs further traced transient gene expression changes along trajectories from progenitors to committed precursors to mature cell types (Fig. 3C). These data showed a dynamically coordinated transition in molecular identity leading to progressive cellular identity. Significantly, meta-gene programs also revealed modules underlying context-specific programs involved in equivalent biological processes. For example, distinct meta-gene programs were associated with differentiated granule cells (gMP5) or Purkinje cells and cerebellar interneurons (gMP7/8) (Fig. 3D). All these meta-gene programs were associated with biological process terms ‘neuron differentiation’, but included many unique genes associated with each meta-gene program.

Notably, for the majority of meta-gene programs, ontology terms did not shed light on cell-type-specific biology, outside of the biologically relevant meta-gene programs described above. For example, terms including ‘Neuronal System’ (associated with 9 meta-gene programs), generation of neurons/neurogenesis (13 times), and axonogenesis (9 times), failed to highlight the distinguishing biology characteristic of individual cell classes. This discrepancy could potentially arise because cell-type-specific ontology terms are usually associated with differentially expressed marker genes, a lack of hindbrain lineage-specific gene sets in the annotation dictionary, or a combination of both. In this respect, we validated that the obtained meta-gene programs are indeed cell-type-specific through overlap with marker genes (Fig. S7E) and enrichment of these meta-gene programs in equivalent cell classes in an independent brain atlas^18^ (Fig. S7F). Thus, the obtained meta-gene programs provide resources to address the lack of cell-specific gene sets, and provide instructive analysis from gene expression datasets of equivalent tissues.

### Cis-regulatory programs and regulatory grammar underlying hindbrain cellular diversity

To further resolve regulatory programs underlying human hindbrain development, we next dissected the underlying CRE landscape. First, we identified differentially accessible marker CREs per class and per cluster. Next, we examined their distribution across chromosomes to determine whether class-specific regulatory activity is spatially restricted (Fig. 3E). In general, for all classes, proportion of marker CREs showed a decreasing trend of enrichment from chromosome 1 to 22, with chromosomes 1/2 among the most enriched and chromosomes 21/22 being the most depleted, correlating with the sizes of these chromosomes. However, comparison of enrichment of marker CREs on a given chromosome across classes yield certain cell-type-specific patterns. For example, chromosomes 16/17/19 were significantly enriched in differentiated neurons (Neurons_I/II) compared to differentiated glia (MixedGlia and MOG), and chromosomes 2/3/18 were significantly enriched in oligodendrocyte progenitors. Notably, the pattern of differential chromosomal activity aligns with the frequency of chromosomal aberration identified in glial versus neural tumors; for example, while high-grade gliomas typically exhibit gain of chromosome 2/3 and loss of chromosome 16,17 and19^41^, neuroblastoma and medulloblastoma frequently exhibit gain of chromosome 17^42,43^, and medulloblastoma further exhibits frequent gain of chromosomes 16 and 19^42^.

The cell type-enriched chromatin activity was restricted to large segments of chromosome, rather than the involvement of the entire chromosome, suggesting clustering of spatially localized CREs (Fig. S8). Thus, we next identified these chromosome segments with spatially clustered co-accessible CREs, constituting cis-regulatory islands (CRIs), using a modified ROSE-based super-enhancer identification approach^44^; we merged signals from closely spaced CREs, which revealed multiple cell-type-specific cis-regulatory islands (Fig. 3F; Fig. S9A). These cis-regulatory islands often abutted known lineage-specific genes, such as *OLIG1/2* or *GFAP*, and island accessibility correlated with neighboring marker gene expression (Fig. 3F). Additionally, genes associated with cell-type-specific cis-regulatory islands showed significantly enriched expression in the respective cell type, compared to other cell populations (Fig S9B).

Further, within a particular cell type, genes associated with cis-regulatory islands showed higher expression compared to other highly expressed and highly variable genes in that cell type, and they were also enriched for marker genes per class (Fig. 3G,H; Fig. S9C,D). Importantly, the cis-regulatory island-associated cell-type-specificity was clearer in glial cell classes such as MOG, compared to neuronal classes, particularly Neuroblasts_I/II/III and Neurons_I/II, which represent heterogenous cell populations, thus leading to lower intensity of cell-type-specific signature (Fig. S9A-D). Next, we clustered the co-accessible CREs, irrespective of their spatial co-localization, into meta-regulatory programs. Similar to the meta-gene approach, we first obtained a set of CREs per class through multi-rank LDA factorization and then clustered these sets to obtain representative meta-cis-regulatory programs (Fig. 3I). Similar to metagene programs, we identified both cell-type-specific meta-regulatory program (rMP1) enriched in Progenitors, program (rMP7) shared between Progenitors, Neuroblasts, Glioblasts and mature glial populations, and a cell-agnostic program (rMP57) (Fig. 3J; Fig S10A-C). Further, these meta-regulatory programs decoupled multiple class and cluster specific co-accessible CRE-set and identified transient programs defining cellular trajectories (Fig. 3K). Importantly, we identified a higher number of programs with greater cluster specificity using the cis-regulatory landscape, compared with using the transcriptomic landscape (Fig. 3A,I; Fig. S7A,C; Fig. S10A,B). These distinguishing characteristics potentially arose from larger sets of accessible CREs compared to expressed genes, but also represented more specific information of cellular identity at CREs than gene expression^22^, suggesting future annotation efforts could benefit from CRE-based definitions to further resolve cellular states.

CREs are sites that encode the regulatory grammar, translated into TFs that bind together with other transcriptional co-regulators, driving cell-type-specific gene expression. To decrypt this regulatory grammar encoded in the meta-regulatory programs, we trained a deep-learning classifier, which we call DeepHB, based on CREsted^45^. We trained and evaluated three different CREsted-based DeepHB models, a convolution neural network (CNN), CNN and RNN model (CNN+LSTM) and a hybrid model initialized with JASPAR motifs family (CNN+LSTM*) (Fig. S11A). All three models had comparable performance, with CNN+LSTM* performing marginally better than the other two; hence, we used this model for further analysis (Fig. 4A; Fig S11B). We first validated our model on an external dataset^19^ for its ability to identify cell-type-specific CREs. Using our model, we were able to classify marker CREs identified for Purkinje cells, the immune cluster, and oligo cells, to their respective cell-type-specific meta-regulatory programs (Purkinje=rMP33, Immune=rMP53-55, Oligodendrocyte=rMP37/39) with high fidelity (Fig. S11C). Next, we used TF-MoDISco^46^ to identify the syntaxes encoded in the CREs comprising the individual meta-regulatory programs, termed “seqlets”, which represent TF motifs. An important feature of these seqlets was the degeneracy of their regulatory syntax, indicating flexible motif configurations that enable context-dependent TF binding. We then merged the obtained seqlets into a set of 156 representative seqlets (minimum size 3 base-pairs, only with positive contribution), and obtained a seqlet count matrix representing the enrichment of each seqlet per meta-regulatory program (Fig. 4B; Fig. S11D). Clustering this seqlet/motif matrix showed three broad patterns of motifs distribution: broadly shared (positive importance in ≥9 meta-regulatory programs), moderately shared (>2 and <9 programs) and mostly unique (<3 programs), which comprised the majority of identified motifs (96 out of 156) followed by ‘moderately shared’ (46 out of 156) (Fig. S11E). Examples of ‘mostly unique’ motifs included the STAT dimer and HOX motifs, TEAD and ETV motifs were ‘moderately shared’, and NEUROD2/OLIG2 and its closely shared NEUROG2 motif were among the ‘broadly shared’ motifs across meta-cis-regulatory programs (Fig. 4B).

**Figure 4:**
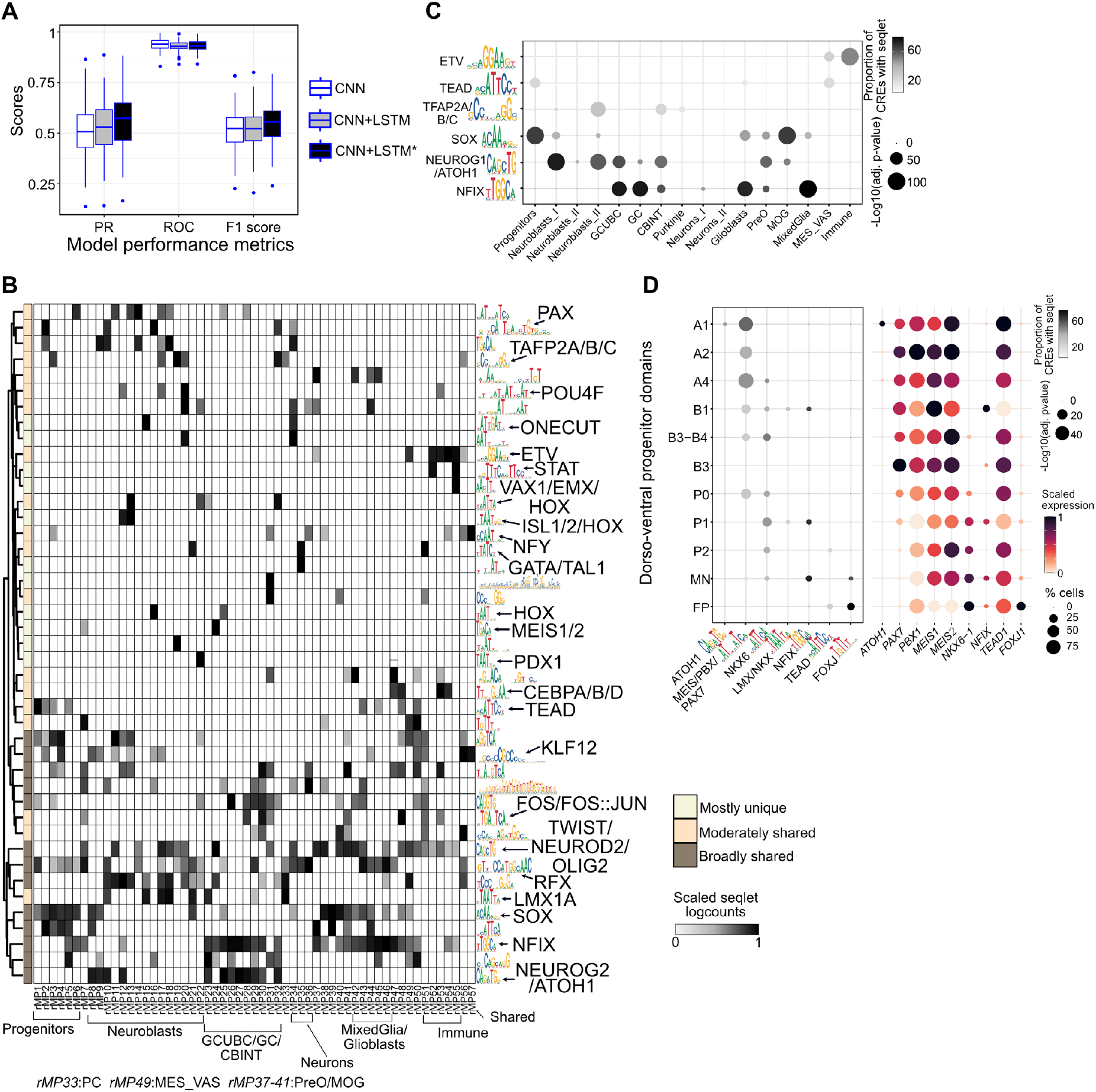
Regulatory grammar driving hindbrain developmental lineages. **A**) Boxplots of model performance metrics showing precision–recall (PR), receiver operating characteristic (ROC) and F1 score, for a convolutional neural network (CNN), a hybrid CNN plus long short-term memory model (CNN+LSTM), and a hybrid model with JASPAR motif initialization (CNN+LSTM*) across meta-regulatory programs. **B**) Heatmap of scaled seqlet counts for selected syntaxes (right) within meta-regulatory program. Class-specific enrichment of meta-regulatory programs shown below. **C**) Selected motif enrichment per class. Bubble size, -1*log_10_(adjusted p-value). Bubble color, percentage of top 500 class-enriched CREs with a motif based on homer enrichment analysis. **D**) Motif enrichment (left) and expression of associated TF (right) per progenitor cluster arranged from dorsal (top) to ventral (bottom) order. Left: Bubble size, -1*log_10_(adjusted p-value). Color, percentage of top 500 marker CREs with a motif based on homer enrichment analysis. Right: Bubble size, proportion of cells expressing. Color, scaled gene expression.

A combination of the meta-regulatory—seqlet matrix and the differential activity of meta-regulatory programs translates into cell-type-specific regulatory grammar. For example, NFIX-motif enrichment in meta-regulatory programs associated with GCUBC/GC/CBINT and PreO/MOG populations, was associated with increased accessibility of NFIX motifs in these cell types (Fig. 4C). Similar patterns were observed for SOX (progenitor classes), ETV (immune), NEUROG1/ATOH (neuroblasts/neurons), and TEAD (progenitor and MES_VAS) motifs (Fig. 4C).

Importantly, while the motif enrichment represents the accessible grammar, an additional level of complexity arises from expression of corresponding TFs, which translates this grammar into transcriptomic identity. The overlap between these two critical regulatory layers represents the regulatory code driving hindbrain cellular diversity (Fig. 4D).

Altogether, this analysis revealed that the co-accessible CRE landscape and its encoded regulatory syntax provide a higher resolution map of cellular identity than the transcriptome, shaping developmental trajectories during human hindbrain development.

### Integrative multi-omics analysis identifies cell-type-specific TF activity

TF motif accessibility and TF expression together represent the permissive and inductive components of the regulatory code (Fig. 4D), which is subsequently translated into the expression of downstream target genes, thereby defining TF-driven gene regulatory networks (TF-GRNs), analogous to the cellular interpretation of regulatory information. Accordingly, we next applied the SCENIC+^47^ framework to infer TF-GRN programs for each class (Fig. 5A). We first identified the number of TFs active per class and observed a positive association between cellular heterogeneity, as defined by the number of distinct subclusters, and the number of TFs associated with TF-GRNs (Fig. 5B). We then investigated the distribution of active TFs across classes and found more TFs that were active in a few classes than TFs shared across classes (Fig. 5C), closely resembling the dominance of ‘mostly unique’ motifs over ‘broadly distributed’ motifs, as described previously (Fig. 4B; Fig. S11E).

**Figure 5:**
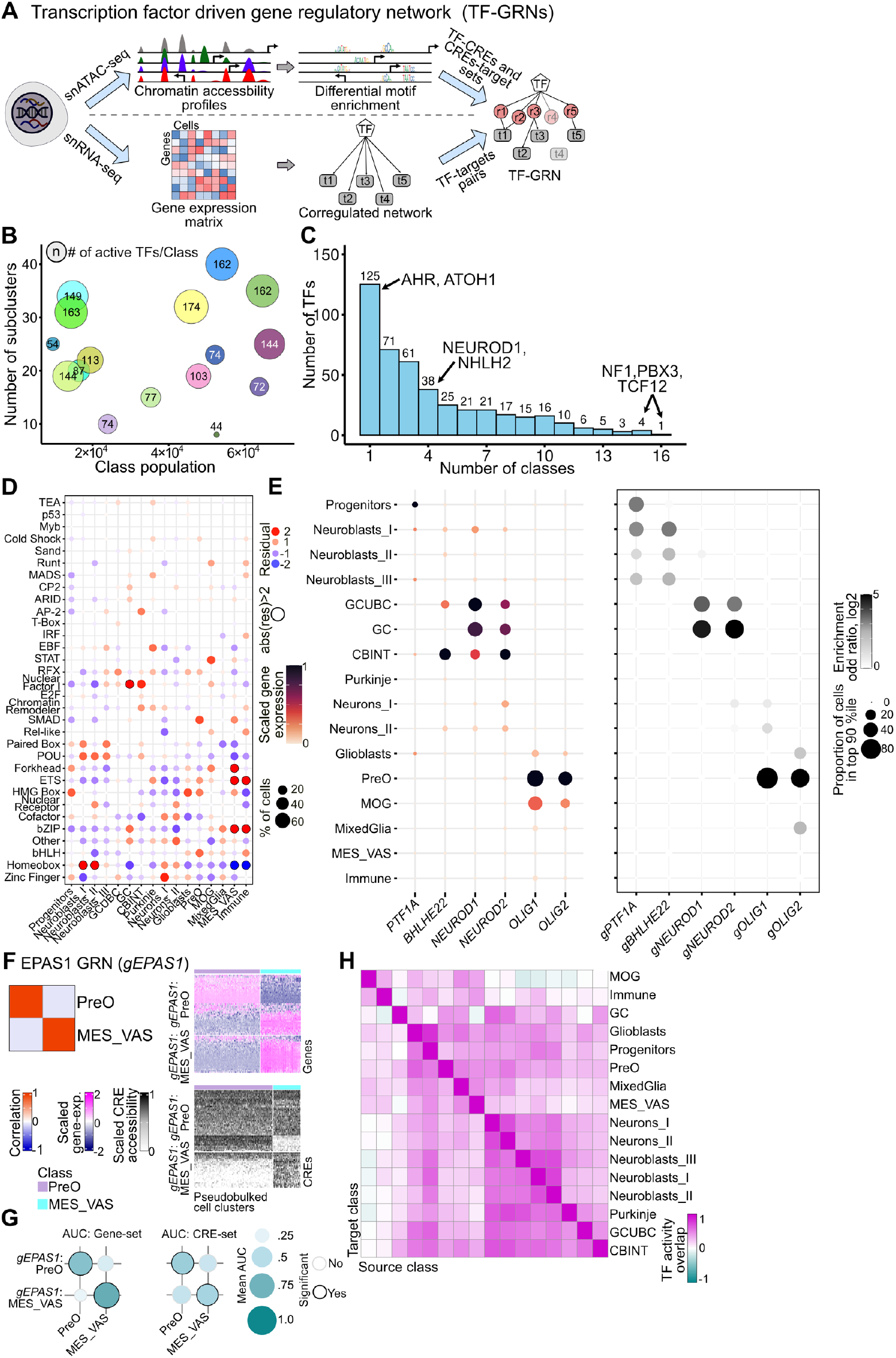
Context specific transcription factor activity. **A**) Schematic description of SCENIC+ approach for identifying transcription factor gene regulatory networks (TF-GRNs). **B**) Number of active TFs per class. Bubble size, Number of TF-GRNs. X-axis, cell-class population. Y-axis, number of sub-clusters. **C**) Number of classes in which each TF is detected. **D**) Negative-binomial test residual values (color and bubble size) fitted on Class and TF-family association. Circled bubbles, absolute residual value above 2 depicting significance. **E**) Scaled expression of TF (left) and enrichment of associated TF-GRNs (right) in cell populations per class in the human hindbrain. Left: bubble size, proportion of cells expressing. Bubble color, scaled gene expression. Right: Bubble size, proportion of cells in the top 90% ranked by gene set activity score. Bubble color, log2 enrichment odds ratio. *gPTF1A*:Progenitors, *gBHLHE22*:Neuroblasts_I, *gNEUROD1*:GCUBC, *gNEUROD2*:GCUBC, *gOLIG1*:MixedGlia and *gOLIG2*:PreO were used as representative TF-GRNs for each of the associated TF. **F**) Correlation heatmap of EPAS1-regulated TF-GRNs (*gEPAS1*) in pre-oligodendrocytes (PreO) versus mesenchymal/vascular cells (MES_VAS) (left). Heatmap of gene expression enrichment for *gEPAS1* in PreO and MES_VAS (top right) and associated CRE enrichment (bottom right). **G**) Pairwise *gEPAS1* (PreO and MES_VAS) gene sets activity (left) and CRE-sets activity scores (right) comparison in PreO and MES_VAS cell populations. Bubble size and color represent mean AUC scores from pairwise Wilcoxon tests. Significant enrichments are circled. **H**) Heatmap depicting similarity between cell classes in terms of conserved or context specific activity of shared TFs. X and y axis represent source and target classes. Order of classes is same on both the axes.

We next grouped TFs together based on their TF-family and evaluated the distribution of their activity across classes (Table S7). Most of the TFs belong to the zinc finger family, followed by Homeobox and bHLH (Fig. S12A). We next investigated the distribution of TF families across classes (Fig. 5D; Fig. S12B,C). Here, we identified the Homeobox family (*e*.*g*. HOX, DLX, LHX factors) as significantly enriched in neuroblasts and absent in immune and mesenchymal cell types. Conversely, ETS (*e*.*g*. ETV5) and the bZIP family were significantly enriched in immune and mesenchymal cell clusters. We also observed differential distribution for other TF families, which did not pass the significance threshold potentially due to sparsity of data but matched with their well-known cell type specific function, such as HMG box TFs (*e*.*g*. SOX factors) in progenitor cell types^48^, EBF (*e*.*g*. EBF1/2/3) in Purkinje cells^49,50^, and AP-2 (e.g. TFAP2A/B) in cerebellar interneurons^51^.

In most cases, the distribution of activity of TF closely matched the restricted accessibility of corresponding motif sites, such for PAX, HOX and ETS factors in progenitors, neuroblasts and immune cells, respectively. However, notable contradictory examples, such as for OLIG2 and NEUROD2, revealed critical insight into how development exploits redundant syntax to achieve cell-type-specific TF activity. These TFs shared a binding motif widely accessible across meta-regulatory programs associated with neuroblasts and oligodendroglia cell types (Fig. 4B), but, due to the restricted expression of the TFs themselves, NEUROD2 and OLIG2 TF-GRN are restricted to neuroblasts and oligodendroglia cell types, respectively (Fig. 5E).

Next, we investigated whether TFs that are active across multiple classes indeed share similar functions. The SCENIC+ approach prunes ‘weak’ TF-target links to obtain a representative TF-GRN set. This leads to disjointed sets, where the same TF has distinct targets, even though it positively influences the expression of the union of these targets in both contexts. Keeping this in view, we compared the activity of distinct TF-GRN sets of the same TF by obtaining correlation of gene sets’scores. Thus, two positively correlated gene sets will have the same trend of expression, and thus function, in distinct cellular context, even if the gene sets have a low overlap of constituents. Using this approach, we then investigated cell-type-specific activity of TFs across multiple classes. For example, EPAS1 was associated with a TF-GRN in both oligodendrocyte progenitors (PreO) and mesenchymal/vascular (MES_VAS) cell populations (Fig. 5F). However, correlation of TF-GRN activity suggested a distinct function, which was also supported by differential expression of targets and associated CREs. To identify the statistically significant differences between expression of gene sets and accessibility of associated CREs, we used a pair-wise Wilcoxon test that identified EPAS1 TF-GRNs obtained in PreO and MES_VAS as having significantly distinct activity in the respective classes (Fig. 5G). In a similar manner, we also obtained other TFs that exhibited further complex behavior, such as ARTN2, which exhibited neural versus glial activity, and NFIC activity in GC was similar to in Neurons_I but less similar to NFIC activity in other classes where it was found to be associated with a GRN (Fig. S12D;E). While the contextual TF activity depends on class-specific CRE, it did not overlap with the array of co-factors (TF binding to shared CREs; Fig. S12F) or coactivators (TF associated with shared genes; Fig. S12G).

Having established that the correlation of TF-GRN gene set activity across cell classes could be used as a measure of shared or context-specific activity, we next evaluated how cell classes were compared to each other in terms of similarity or dissimilarity of activity of TFs. We identified similarity between a pair of cell classes based on positive correlation of activity of shared TF and penalizing for negative correlation (Fig. 5H). This similarity analysis once again clustered closely related classes such as pontine/ medullary neuroblast and neurons together in one group, radial glia and gliogenic progenitors (Progenitors, Glioblasts and PreO) into another group. Notably, in terms of shared TF-GRN activity, differentiated oligodendrocytes (MOG) were closer to Immune cells than their own precursors (PreO).

Altogether, by integrating transcriptomic and chromatin accessibility landscapes, we demonstrated that the tripartite interaction between TF expression, motif enrichment and accessibility of specific regulatory elements, translates into context-specific regulatory activity and transcriptomic states.

### Divergent regulatory programs determine regional identity of rhombic lip lineages

As described above, DV patterning is a major driver of neuronal heterogeneity, whereas the influence of AP patterning is comparatively subtle. However, this AP influence is particularly evident when comparing cerebellar neurons with those of the pons and medulla (Fig. 1C). A striking feature distinguishing these axial hindbrain domains is *HOX* gene expression, suggesting a role for HOX genes in establishing regional identity^28,29,35^. We therefore investigated the molecular and regulatory differences between trajectories derived from the ATOH1^+^ domain, specifically granule cells originating from the upper rhombic lip and pontine nuclei originating from the lower rhombic lip^52,53^ (Fig. 6A-C; Fig S13A,B). Although these two lineages arise from shared ATOH1^**+**^ precursors and exhibit overlapping meta-gene (gMPs) and meta-regulatory programs (rMPs) (Fig. S7A; Fig. S10A), they diverge in their transcriptomic identity, migratory routes, and function^52-54^ (Fig. S13A). We hypothesized that differential activity of TF-GRNs underlies this divergence. Consistent with this, we identified TF-GRNs associated with TGF-β signaling, WNT signaling, and HOX genes that were preferentially enriched in pontine nuclei precursors compared with granule cell precursors (Fig. 6D-F).

**Figure 6:**
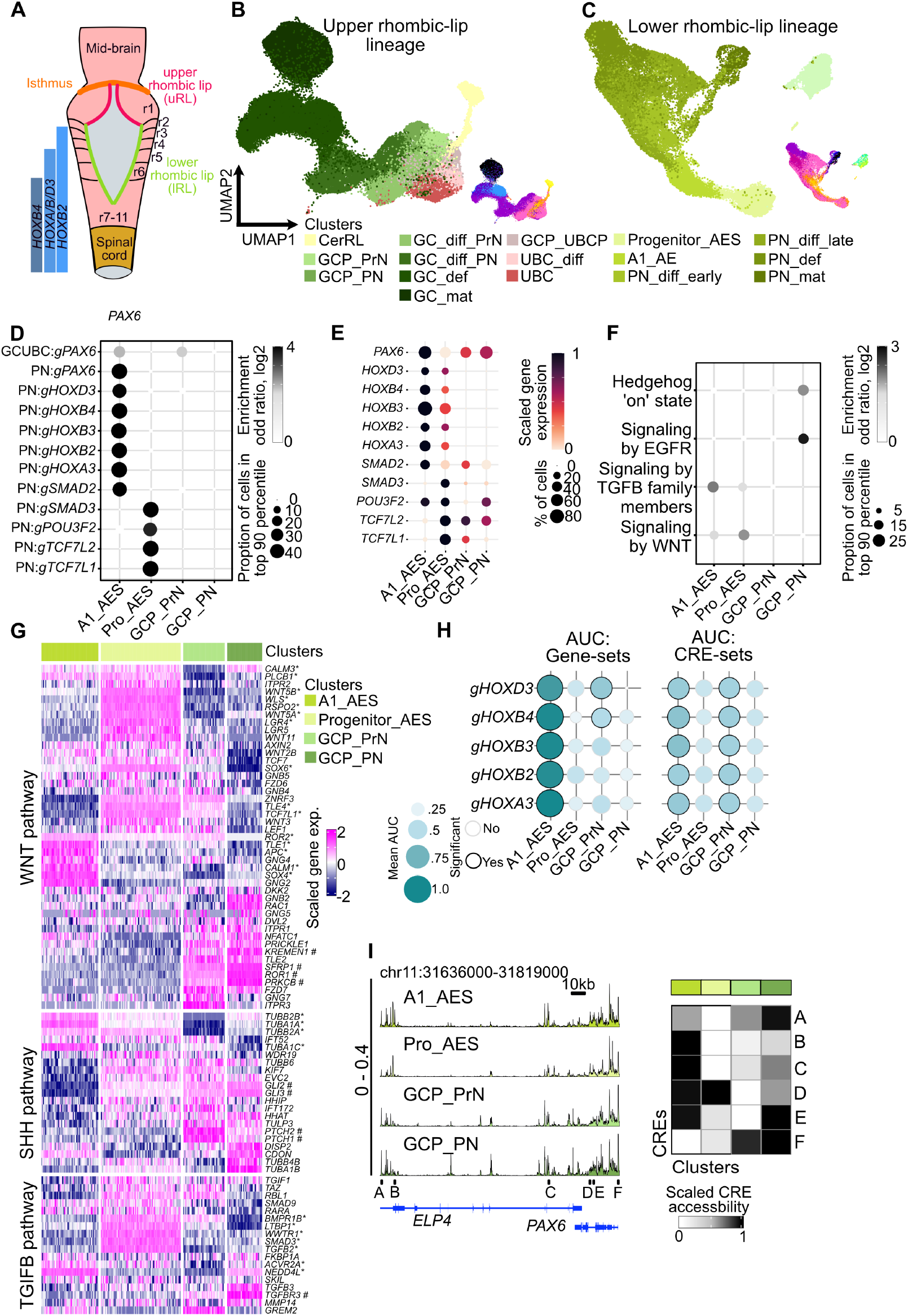
Divergence of ATOH1 lineage in upper and lower rhombic lip. **A**) Schematic of the upper and lower rhombic lip domains in the embryonic hindbrain. **B**) UMAP of the granule cell lineage colored by cluster identity. Inset, cells colored by stage. **C**) UMAP of the pontine nuclei lineage colored by cluster identity. Inset, cells colored by stage. **D**) Differentially enriched TF-GRNs in granule cells versus pontine nuclei precursors. Bubble size, proportion of cells in the top 90% ranked by gene set activity score. Bubble color, log2 enrichment odds ratio. **E**) TF expression in granule cell versus pontine nuclei precursors. Bubble size, proportion of cells expressing. Color, scaled gene expression. **F**) Differentially enriched signaling cascades in granule cell versus pontine nuclei precursors. Bubble size, proportion of cells in the top 90% ranked by gene set activity score. Bubble color, log2 enrichment odds ratio. **G**) Heatmap of SHH, WNT, and TGF-beta pathway genes expression in granule cell and pontine nuclei precursors. Columns, pseudo-bulked clusters. Rows, genes. *, genes differentially upregulated in pontine nuclei lineage. #, genes differentially upregulated in granule cell lineage. **H**) Pair-wise comparison of HOX TF-GRN gene sets activity (left) and CRE-set activity scores (right) associated in granule cell and nuclei pontine precursors. Bubble size and color represent mean AUC scores from pairwise Wilcoxon tests. Significant enrichment circled. **I**) Chromatin accessibility profile at the *PAX6* locus in granule cell and pontine nuclei precursors (left), with HOX binding sites (A–F) indicated; heatmap of scaled chromatin accessibility across HOX binding regions (right).

Notably, TCF and SMAD factors, while being enriched in lower rhombic lip precursors, were present in upper rhombic lip precursors. As the activity of these TFs factors is downstream of signaling cues^55,56^, differential accessibility to the inducing signal (RSPO2, WNT5A/B, LTBP1), expression of receptors (LGR4, BMPR1B, ACVR2A), or presence of antagonizing signals (SHH signal from Purkinje cell to GCPs^57^), could contribute to differential activity of these GRNs in distinct cell context (Fig. 6G).

AP-patterning along the body axis is primarily mediated by *HOX* genes that define axial domains in the hindbrain and spinal cord^58,59^. Therefore, we next investigated the regulatory dynamics underlying HOX-driven gene expression. Surprisingly, while HOX target gene sets were statistically enriched in the pontine nuclei neuroblasts, their associated CREs did not show corresponding differential enrichment (Fig. 6H). We also did not detect differential accessibility of CREs of HOX targets that are bound by non-HOX TFs, ruling out a dominant contribution of non-HOX factors in their differential expression (Fig. S13C). Further, promoter elements were not differentially enriched in HOX binding sites, compared to non-HOX binding sites (Fig. S13D). These data suggest that HOX-associated gain of gene expression is primarily driven by gain of *HOX* expression, translating into HOX activity, without a correlated reshaping of the permissive regulatory landscape, as shown by the representative example of HOX-dependent *PAX6* expression (Fig. 6I). We observed a similar gain of TF activity primarily driven by TF expression in the case of TCF and SMAD factors (Fig. S13E). In contrast, the most strongly differentially expressed genes between granule cell and pontine nuclei lineage precursors were linked to differentially accessible regulatory elements (Fig. S13F), which suggests that some significant transcriptomic changes do follow regulatory reshaping.

Overall, these comparisons suggest that spatial cues, including DV- and AP-signaling pathways, shape cellular identity in the rhombic lip lineages primarily by modulating TF expression (*e*.*g*., *HOX* genes) or activity (*e*.*g*., TCF7L1/2, SMAD2/3), while the regulatory landscape remains poised to respond to changes in TF activity.

### Developmental origins of pediatric glial tumors

We next used the hindbrain atlas to infer lineages of origin of hindbrain-localized pediatric brain tumors. We mapped the transcriptomic signatures obtained from bulk gene expression data for a set of pediatric brain tumors, including other non-hindbrain CNS tumors, to the transcriptomic hindbrain atlas. This all-vs-all comparison highlighted glial tumors enriched for glial-cell-type signatures. For example, high- and low-grade glioma were enriched for oligo and astrocytic cell clusters, while ependymoma best matched to ependymal cell clusters (Fig. S14). In contrast, embryonal tumors such as medulloblastoma matched to upper rhombic lip derived cell lineages, and ATRT matched to progenitors/ radial glia cell populations (Fig. S14).

To further dissect disease biology at single-cell resolution in the frame of reference of normal development, we next focused on pediatric glial tumors, particularly pilocytic astrocytoma (PA), diffuse midline glioma (DMG) and posterior fossa ependymoma (PFA), which are known to arise from distinct glial lineages^60^ (Fig. S14). We generated new single-nucleus transcriptomic data for PA (3 sample, 24,972 cells) and DMG (5 samples, 21,735 cells), and obtained published data for PFA^61^ (3 samples, 17,737 cells) (Fig. S15A; Table S8).

First, we determined similarities between the tumors and developmental lineages in hindbrain development. Using an SVM-based hindbrain classifier, we resolved the cellular composition across these tumors. We identified the tumor component in PA and DMGs as dominated by oligo and astrocyte-like cells, and in PFA by ependymal-like cells, with microglia comprising the predominant tumor microenvironment across these samples (Fig. 7A,B; Fig S15B), matching the inference gained from bulk transcriptomic analysis (Fig. S14). Meta-gene programs associated with tumor compartments also aligned with the meta-gene programs enriched in the equivalent normal cell populations (Fig. S15C). Between PA and DMG, DMG exhibited a greater cellular heterogeneity within the oligo- and astrocytic-tumor compartments, and a higher proportion of unclassified cells, suggesting a noisier transcriptomic signature in DMG (Fig. 7A,B; Fig. S15B). Notably, oligo-like progenitors in PA matched to postnatal OPCs (OPC_ PNAD) while DMG exhibited a mix of pre- and postnatal OPC types (OPC_PrN and OPC_PNAD), aligning with proliferative capacity of these tumors and the matching oligo-progenitors (Fig. S1G). Importantly, both PA and DMG lacked potency to differentiate oligo-progenitors into mature oligodendrocytes, potentially arising from a failure to downregulate TF-GRNs associated with normal oligo-progenitors (Fig. S15D). While oligo-like progenitors in tumors lacked terminally differentiated oligodendrocytes, they matured along the astrocytic lineage in both PA and DMG, which suggests potential bipotency of oligo-progenitors in the tumor context^62^.

**Figure 7:**
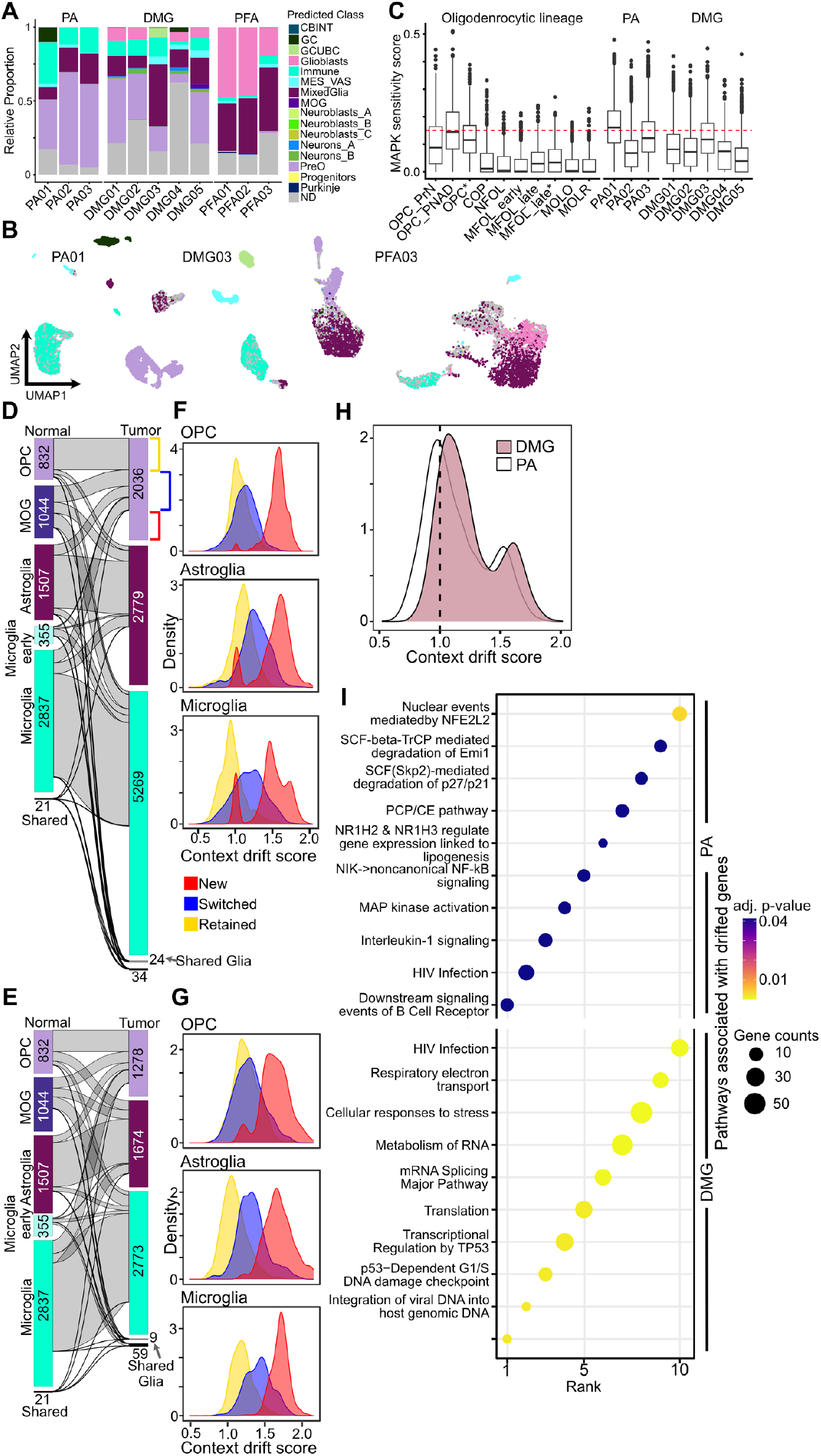
Gene context drift driving pediatric glioma tumor biology. **A**) Relative cellular composition of individual tumor samples belonging to pilocytic astrocytoma (PA), diffuse midline glioma (DMG) and posterior fossa ependymoma (PFA) in terms of cell classes in the hindbrain atlas obtained using a SVM classifier. Bars, predicted class identity. **B**) UMAP representation of annotated cells in representative samples of PA (PA01), DMG (DMG03) and PFA (PFA03). Cells are colored by class identity. **C**) MAPK pathway activity score^67^ for cell populations in normal hindbrain oligodendrocyte lineage, and oligo-compartments in PA and DMG tumors. Asterisk denotes cell clusters from MixedGlia class. **D**,**E**) Sankey plot showing gene community overlaps between shared glial populations in the reference atlas and a representative PA (**D**) and DMG (**E**) sample. **F**,**G**) Density plots of context-drift scores for retained, switched and gained (new) genes in each tumor gene community (colored traces) for PA (**F**) and DMG (**G**) sample. **H**) Density plot of average context drift score for same set of genes in obtain from analysis of 3 PA and 5 DMG samples. **I**) Top 10 Reactome terms among the top 500 drifted genes in PA and DMG samples.

Next, we focused on dissecting tumor-specific biology, comparing the cellular compartments in tumors to their equivalent normal cell types in the developing hindbrain. Comparison of tumor-associated microglia gene-signature with normal microglia revealed that, across tumor types, microglia exhibited more tumor-specific than shared gene ontology (GO:BP) terms among both upregulated and downregulated gene sets, highlighting a highly tumor-dependent microglial response (Fig. S16A). Notably, while microglia in all three tumor groups exhibited a shared pro-inflammatory cytokine response (*CCL3/4, TNF*), DMG-associated microglia additionally activated an anti-inflammatory response (*TREM1, PTGS2, PTGER4*) (Fig. S16B). In contrast, PFA-associated microglia also upregulated an extracellular matrix gene signature (Fig. S16C), hinting their PFA-specific immunosuppressive^63^ and pro-metastatic role^64,65^.

We next compared differentially expressed genes in the proliferating tumor compartment, focusing on oligo-compartments in PA and DMG. Surprisingly, we did not detect significant enrichment of any known oncogenic signaling pathways in either of the tumors, in contrast to the known oncogenic role of MAPK signaling in PA^66^. We next examined MAPK pathway activity during hindbrain development, which showed that the oligo-progenitor clusters (OPC_PrN, OPC_PNAD) do not exhibit significant enrichment for expression of MAPK pathway gene sets when compared to other cell clusters (Fig. S16D). However, within the oligodendrocytic lineage, MAPK pathway activity, as evaluated form enrichment of 10 pathway genes^67^, increases in oligo-progenitors and then downregulates during further differentiation (Fig. 7C). Additionally, MAPK activity within normal oligo-progenitors was comparable to MAPK activity in the proliferating oligo-compartment in PA. Among the distinguishing features of oligo-compartments of PA and DMG, were upregulation of an extracellular matrix gene-signature in PA (Fig. S16E), with potential immunosuppressive and pro-tumorigenic role^64,65^, and upregulation of synapse assembly pathways in DMG (Fig. S16F), previously shown to promote tumor aggressiveness through tumor-neuron crosstalk^68-70^.

Thus, through this comparative analysis, we revealed a striking preservation of basal cellular biology in tumors, including within proliferative compartments, coupled with an absence of overt oncogenic signaling at the transcriptomic level. Instead, oncogenic rewiring in these developmentally stalled tumors manifests through the emergence of tumor-promoting cellular states, including extracellular matrix-associated programs and anti-inflammatory signatures within both tumor cells and the tumor microenvironment.

*Context drift driving pediatric glial tumor biology*

Similar to cellular annotation, where differential expression represents important cell-specific biology, but fails to resolve coordinated gene expression, tumor versus normal comparison differential gene expression also fails to capture subtle, yet coordinated changes in biology. To capture such coordinated changes that do not pass the significance cut-off, we analyzed changes in geometry or context of gene expression, defined through gene community structure. Using gene-gene correlations to identify gene communities and a natural language processing-based approach to identify genes’ context in these communities^71^, we identified the rewired context of genes between equivalent cell types in the normal and tumor datasets. A change in a gene’s nearest neighbors, and the changes in neighborhood of those genes, is calculated in terms of context drift score per gene to describe the gene’s context change in a cellular compartment^71^ (see *Methods* for details). A comparative analysis identified that equivalent cell classes in tumor and normal maintain a basal gene community, switch in other gene communities and gain new gene communities (Fig. 7D,E). The newly ‘gained’ genes have the highest context-drift scores, as these have undergone the highest change in their gene neighborhood geometry, followed by switched genes, in comparison to genes maintaining their community membership across all cellular compartments in a tumor (Fig. 7F,G). Notably, comparing average drift score across a shared set of genes between PA and DMG exhibited a higher drift in DMG compared to that in PA, suggesting that highly aggressive DMG have undergone further shifts in their biology compared to less aggressive PA (Fig. 7H). Notably, context drift did not correlate with the magnitude of gain in expression (Fig. S16G), suggesting genes could switch their context without drastic changes in their own expression, highlighting the importance of even subtle changes in gene expression towards driving disease biology.

Druggable genes with high context drift were shown to be promising therapeutic targets in primary and relapsed tumors in a recent study^71^. Thus, to explore novel targetable pathways in PA and DMG, we performed reactome pathway analysis for the top 500 drifted genes (Fig. 7I; Table S9,S10). Notably, in PA, the enriched pathways such as NFE2L2 mediated events or PCP/PE pathway, and even MAPK pathway genes including *MAPKAP2* and *DUSP4*, were associated with tumor microglia compartment and not the tumor oligo- or astrocytic-compartments. This result suggests that microglia in low-grade glioma undergoes a significant shift in transcriptomic identity in response to the tumor, perhaps to sustain these slow-growing tumors. Importantly, genes associated with transcriptional and translation events including mRNA-splicing machinery were enriched among the top drifted genes, which matches with the increased splicing burden in high-grade gliomas (such as DMG) compared to low-grade gliomas (such as PA)^72^, along with a suggested oncogenic role of splicing machinery in high-grade gliomas^73-75^. This oncogenic driver pathway, which was not enriched in the differential overexpressed gene set, highlights the importance of subtle, but coordinated, changes in gene expression that has substantially larger impacts on cellular context, driving tumor biology and representing novel therapeutic targets.

Combined with the preservation of basal cellular biology, these results showed that oncogenic transformation arises from extensive rewiring of gene context, achieved through epigenomic reprogramming as in DMG, within conserved cellular states, and reveal therapeutic vulnerabilities missed by differential expression analyses.

### HindbrainExplorer provides access to single-cell multiomics hindbrain atlas

To serve the interest of the broader scientific community and propel further discoveries, we have provided unrestricted access to our analyzed multi-omics hindbrain atlas in a userfriendly way through a dedicated web-based application at https://apps.kaessmannlab.org/HindbrainExplorer/. Through the application, users can evaluate the cluster annotations and investigate the expression of gene(s) of interest in clusters/subclusters and cellular lineages. Users can also evaluate accessibility of 446,510 peaks across the clusters in the snATAC-seq atlas and identify potential cell type specific activity of a genomic region of interest (of 500 bp size) through our DeepHB model. For example, using DeepHB, we predicted that the enhancer class of regions harboring 69 out 83 SNPs that are statistically associated with inheritance of Alzheimer’s disease^76^, with the majority of them potentially acting within microglia enhancers (Fig. S17A). Often, SNPs associated with classified enhancers, suggesting a potential impact of the SNP harboring region on the regulatory grammar (Fig. S17B). Further, the application provides information regarding TF-GRN activity across clusters/sub-clusters per class, and cell-type-specific or shared activity of TFs across classes. Finally, the application provides interactive access to the Hindbrain classifier, which predicts best matching cell-type identities for a user-provided single-cell/nuclei RNA-seq dataset among 151 cell clusters or 16 classes in the Hindbrain reference atlas.

## DISCUSSION

Despite the functionally important role of the hindbrain, its constituent cellular diversity and underlying regulatory framework remain incompletely resolved. Here, we address this gap by generating a comprehensive single-cell multiomics atlas spanning embryonic to adult stages, integrating newly generated transcriptomic and chromatin-accessibility profiles from the developing pons and medulla with published multi-omics datasets from the human cerebellum^14,23^. This unified resource provides a multi-omics, single-cell resolution view of hindbrain development.

We constructed an extensively annotated cellular landscape to provide a high-resolution map of progenitor, neuronal and glial cell-populations across hindbrain development. To deepen our understanding of hindbrain development, beyond insights from differentially expressed features, we identify regulatory programs capturing coordinated gene expression and chromatin-accessibility modules. These programs exhibit varying degrees of cell type specificity and reveal transitional regulatory states that underpin lineage commitment. In doing so, we expand the repertoire of cell type defining molecular signatures and provide a richer resource for future investigations into hindbrain regulatory logic. Notably, we identify a more specific cell type definition in the cis-regulatory landscape compared to the transcriptomic landscape. This could potentially arise from the differences in total number of features considered, 446,510 peaks versus 37,226 genes, but also from involvement of multiple cisregulatory features in precise regulation of a transcriptomic feature^77-79^. These findings suggest that cis-regulatory based definitions should supplement current approaches that exclusively utilize marker gene expression. This intuitive shift in annotation first requires establishment of a CRE-cell-type dictionary through integration of already generated multi-omics data^14,17,19,23^.

Through deep learning^45^ and integrative multi-omic^47^ analysis, we resolve a multi-layered regulatory code in which CRE accessibility, TF-motif enrichment and TF expression jointly define hindbrain lineages through combinations of shared and unique features. However, two significant complexities are also present: degeneracy and redundancy of the regulatory grammar. A TF, rather than binding to a perfect motif, can bind to a family of similar motifs, and second, multiple TFs can bind to the same family of code. While redundancy within this regulatory grammar is modulated by the availability of the cognate TFs, as illustrated by the NEUROD2/OLIG2/BHLHE22 regulatory networks, how specificity is achieved with a degenerate code needs to be further resolved. Moreover, for a given TF-motif pair, variation in accessibility of motif-bearing sites determines context-dependent TF activity, revealing how cellular diversity emerges from a “mix-and-match” usage of regulatory components. These findings also highlight a fundamental question in lineage specification, does TF expression precede and shape CRE accessibility, or does accessible chromatin first create the permissive environment for TF deployment?

A particularly striking insight emerges from the upper versus lower rhombic lip lineages. Although these progenitors share nearly indistinguishable transcriptomic and chromatin-accessibility states, suggesting a common intrinsic regulatory baseline, they diverge into distinct neuronal lineages due to differences in their extrinsic signaling environment. This demonstrates how context-dependent signaling can redirect equivalently primed progenitors toward different developmental trajectories, providing a mechanistic explanation for how the hindbrain generates lineage diversity from uniform starting states.

Beyond understanding normal development, this resource is equally instructive in dissecting disease biology. Through representative examples, we resolve disease-specific and developmentally-shared biology in pediatric brain tumors. The hindbrain atlas provides an excellent reference to resolve tumor cellular composition and a baseline to unravel tumor specific oncogenic transformations. Significantly, in the case of pediatric brain tumors, we show that a tumor inherits the basal cellular identity of its lineage of origin, and builds up the oncogenic identity on top of that, while influencing the normal cells in the vicinity, creating a subtype-specific tumor cellular ecosystem. Beyond uncontrolled proliferation and halted differentiation, the oncogenic transformation includes subtype-specific transformation such as acquisition of extracellular matrix structure in PA or heightened synapse formation in DMG, with each transformation playing a supportive tumorigenic role^63-65,68-70^. Further, we show that in addition to significant changes, subtle yet coordinated shifts in transcriptomic identity also play a critical role in driving oncogenic identity. As in normal development, the relative contributions of cis-regulatory disruption versus primed-state responses to altered inductive cues, including changes in the tumor microenvironment, remain to be resolved and will require further investigation, for which the present multi-omics atlas provides an essential framework.

Lastly, we have compiled this resource in an accessible portal, *HindbrainExplorer*, to provide a user-friendly platform to allow broader scientific community in extracting further novel insights in cell type diversity, molecular markers and pathogenesis. In novel datasets obtained from hindbrain in context of normal or diseased tissues, the obtained transcriptomic and cis-regulatory signatures will be instructive in identifying cellular composition. The robust transcriptomic and cis-regulatory programs will also improve gene set or CRE-set enrichment analysis in molecular investigations associated with hindbrain biology.

### Limitations of the study

Our dataset has certain limitations arising from biological and technical factors.

Biologically, cellular annotations will benefit from inclusion of high resolution spatial transcriptomic approaches as in current data we failed to resolve progenitor/neuronal populations along the AP axis. Similarly, glial cell annotation would also benefit from spatial information regarding cellular neighborhoods, as these functionally distinct cell states are difficult to separate from transcriptional data alone. Due to technical challenges in obtaining neuronal cells from post-natal and adult tissues, which have a higher proportion of glial cells, our atlas also has reduced representation of mature neuron types particularly those of pons of medulla. Further, our annotation of cell states identified in chromatin accessibility atlas suffers from limitation of label transfer approaches. A true multi-omics data from the same cell would help improve annotation of ATAC-seq data in future efforts. Notwithstanding these limitations, our atlas provides a foundational resource for future molecular investigation focusing on hindbrain, and will provide crucial insights regarding regulatory framework in context of development and pathogenesis for this critical CNS region.

## Author Contributions

Conceptualization: PJ, HK, SMP, LMK

Data acquisition: PJ, MS, CS, JS, AW, MBJ

Data analysis: PJ, MS, IS, KO, TY

Methodology: PJ, MS, IS, KO, TY, ST

Resources: MS, IS, AAP, MBJ, JPM, KS, MBJ, BK, CMvT, OW, SL, MP, DTWJ

Website: PJ, NT

Funding acquisition: HK, SMP, LMK

Project administration: HK, SMP, LMK

Supervision: MS, ST, HK, SMP, LMK

Writing - original draft: PJ, MS, IS, KO, TY, HK, SMP, LMK

Writing - review & editing: All authors

## Data and code availability

Raw data will be provided after data transfer agreement. Scripts use in data processing are available on github: github.com/piyushjo15/HumHBAtlas

## Acknowledgments

We would like to express our gratitude to Human Developmental Biology Resource (HDBR; supported by grant MRC/Wellcome (MR/X008304/1 and 226202/Z/22/Z)), Maryland Brain Collection at the Maryland Psychiatric Research Center (NIH NeuroBioBank), Chinese Brain Bank Center, and Human Brain Tissue Bank at Semmelweis University for providing human samples. The study was funded in part by the European Research Council (ERC Consolidator grant 819894 (BRAIN-MATCH) to S.M.P and the ERC Advanced Grant 101019268 (VerteBrain) to H.K.), the European Horizon 2020 Programme - ‘iPC - individualizedPaediatricCure’(Grant 826121 to S.M.P). M.S. was supported by a Simons Foundation Autism Research Initiative Bridge to Independence Award (SFI-AN-AR-Ind ependencePostdoctoral-00007139) and the Mobilitas 3.0 project (MOB3ERC112) co-financed by the European Union and the Estonian Research Council. D.T.W.J. is supported by the EVEREST Centre for Low-grade Paediatric Brain Tumours (The Brain Tumour Charity, UK; grant number GN-000707). The study was also funded in part by the DFG Emmy Noether Program (Grant 551030459 to L.M.K.). The INFORM program is financially supported by the German Cancer Research Center (DKFZ), several German health insurance companies, the German Cancer Consortium (DKTK), the German Federal Ministry of Education and Research (BMBF), the German Federal Ministry of Health (BMG), the Ministry of Science, Research and the Arts of the State of Baden-Württemberg (MWK BW), the German Cancer Aid (DKH), the German Childhood Cancer Foundation (DKS), RTL television, the aid organization BILD hilft e.V. (Ein Herz für Kinder) and the generous private donation of the Scheu family. We also would like to express our sincere thanks to Carsten Maus, Erjia Wang (Genomics and Proteomics Core Facility, DKFZ) and Lena Weiser, Gregor Warsow (Omics IT and Data Management Core Facility, DKFZ) for their highly dedicated support in data management and processing.

