## Supplemental Figures + Methods for "A multi-omics atlas of human hindbrain development"

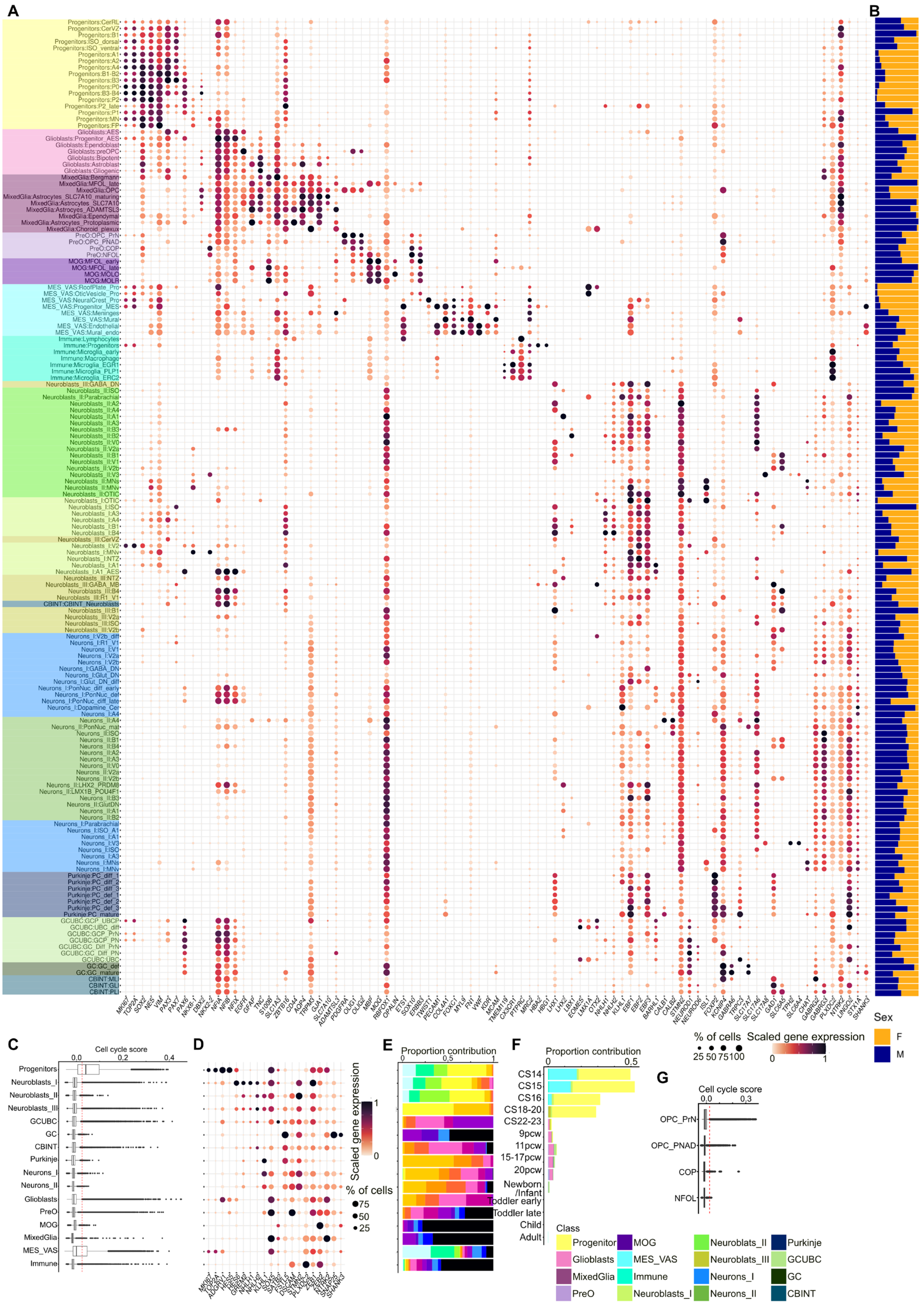




**Supplemental Figure S1** *legend next page* ▶

◀*cont. from previous page*

**Fig. S1: Marker based annotation of clusters in transcriptomic atlas.**

**A**) Expression of selected marker of neuronal or non-neuronal identity identified in the transcriptomic atlas in cell populations grouped by cluster identity. Clusters are named as 'Class:Cluster' format. Bubble size, proportion of cell population with expression. Color, scaled gene expression. **B**) Proportional contribution of cells by sex of donor cells per cluster. **C**) Boxplot depicting cell-score distribution by class. Red dashed line represents cell cycle = 0.025. **D**) Expression of selected markers associated with neuronal maturation per class. Bubble size, proportion of cell population with expression. Color, scaled gene expression. **E**) Proportion contribution of cells by stage per class. **F**) Proportion contribution of proliferating cells (cell cycle score >0.025) per stage colored by class. **G**) Cell cycle score distribution in PreO cell population by cluster. Red dashed line represents cell cycle = 0.025.

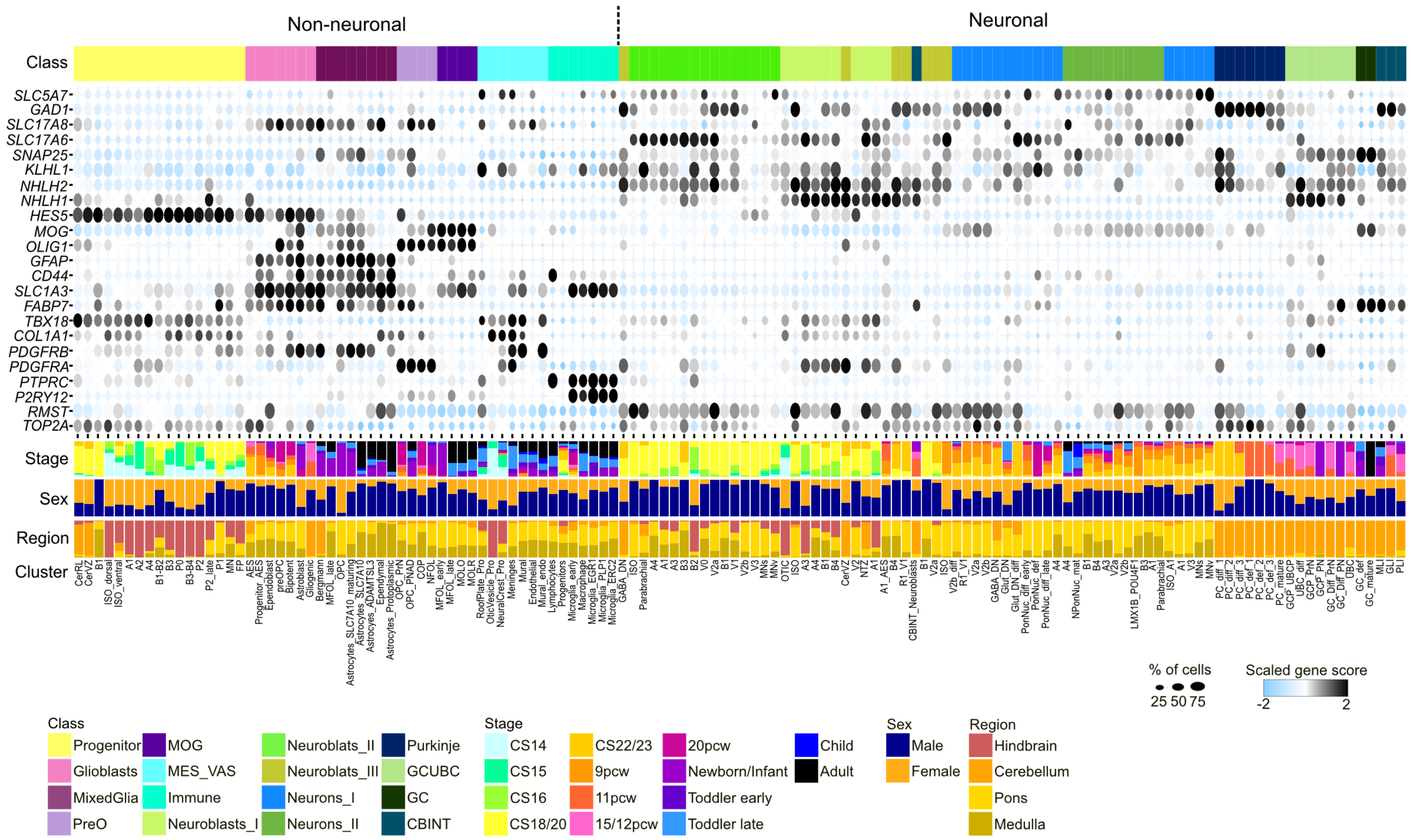


**Fig. S2: Annotated cell clusters in the snATAC-seq atlas.** Marker gene scores for clusters in the chromatin accessibility atlas. Bubble size, proportion of cells. Color, scaled gene score. Top bar shows columns colored by class identity per cluster. Bottom bars show proportional contribution of cells by stage (top), sex (middle) and region (bottom) per cluster.

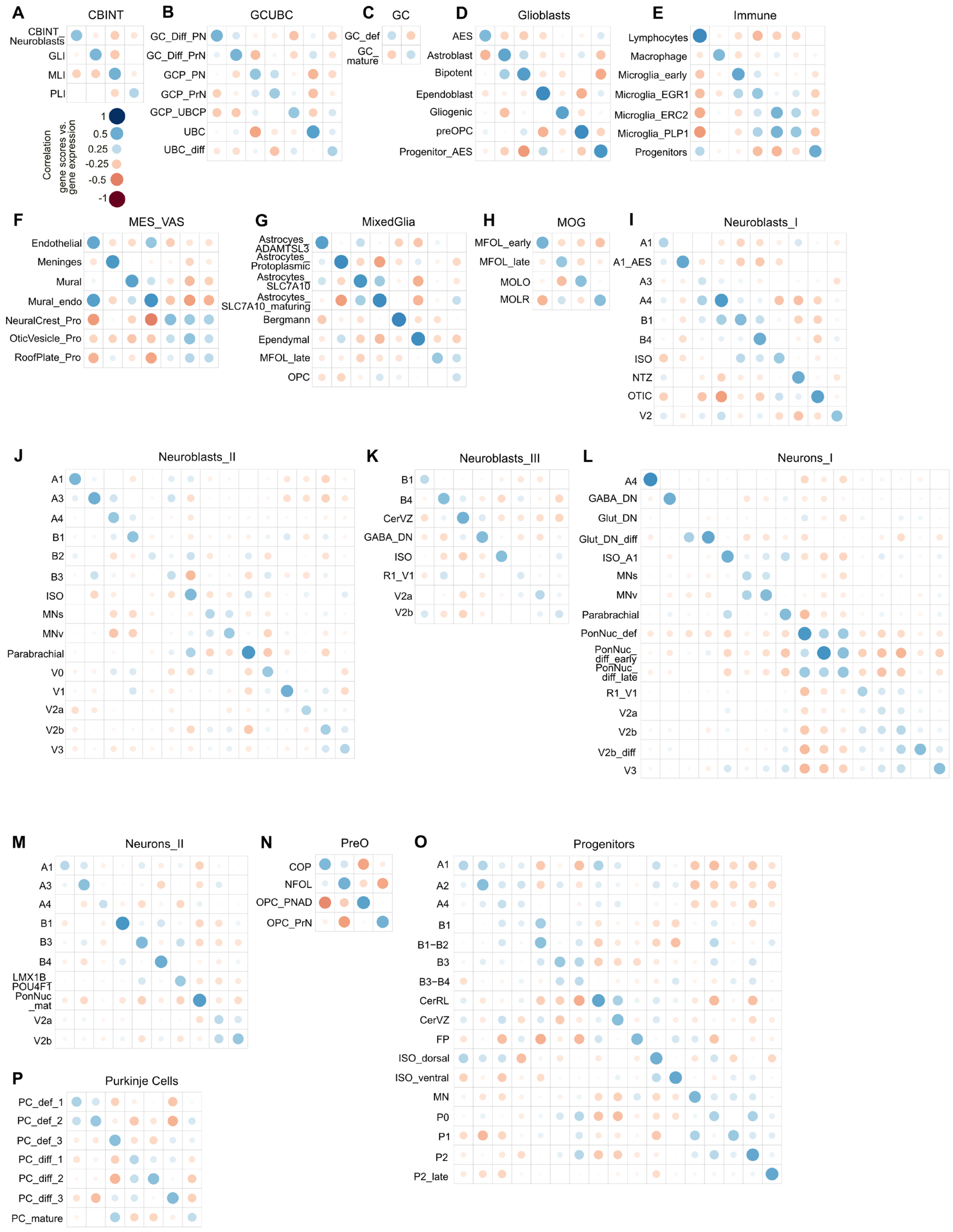


**Supplemental Figure S3** *legend next page* ►

◀*cont. from previous page*

**Fig. S3: Correlation between reference cell clusters in the transcriptomic atlas and equivalent clusters identified in the chromatin accessibility atlas.**

**A-P**) Correlation heatmap representing Pearson correlation between scaled pseudobulked gene expression (snRNA-seq) and gene score (snATAC-seq) profiles per cluster in the reference snRNA-seq atlas (x-axis) and annotated clusters in the snATAC-seq atlas (y-axis) obtained from label transfer per class. Cluster order on x-axis columns is same as the labels on y-axis.

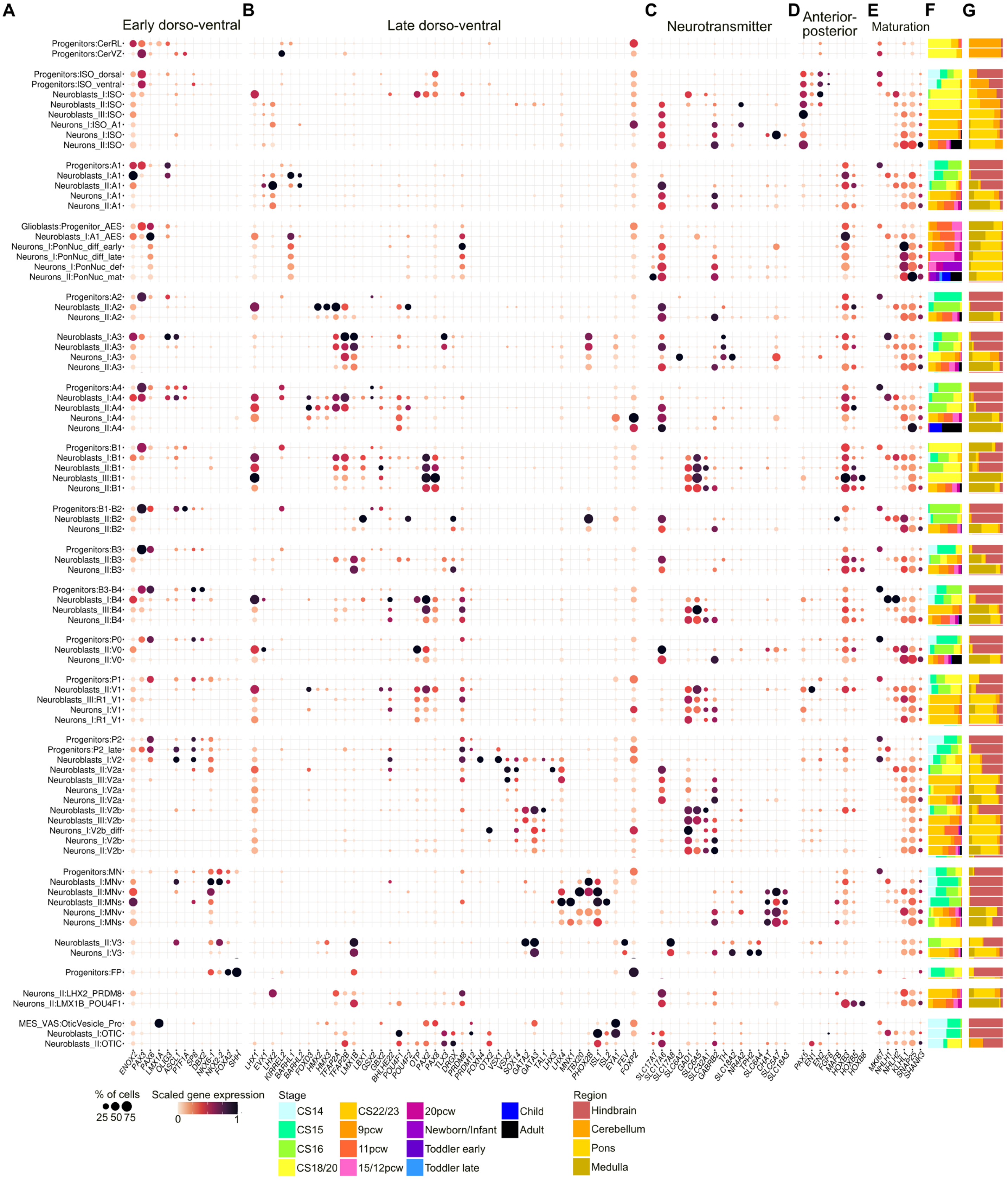


**Supplemental Figure S4** *legend next page* ►

◄*cont. from previous page*

**Fig. S4: Marker based annotation of progenitor and neuronal populations in the pons and medulla.**
**A-E**) Expression of selected markers for early dorso-ventral (**A**), later dorso-ventral (**B**), neurotransmitter (**C**), anterior posterior (**D**) and maturation status (**E**) identity of cell clusters. Bubble size: proportion of cell population with expression. Color, scaled gene expression. **F-G**) Proportional contribution of cells per cluster by stage (**F**) and region (**G**).

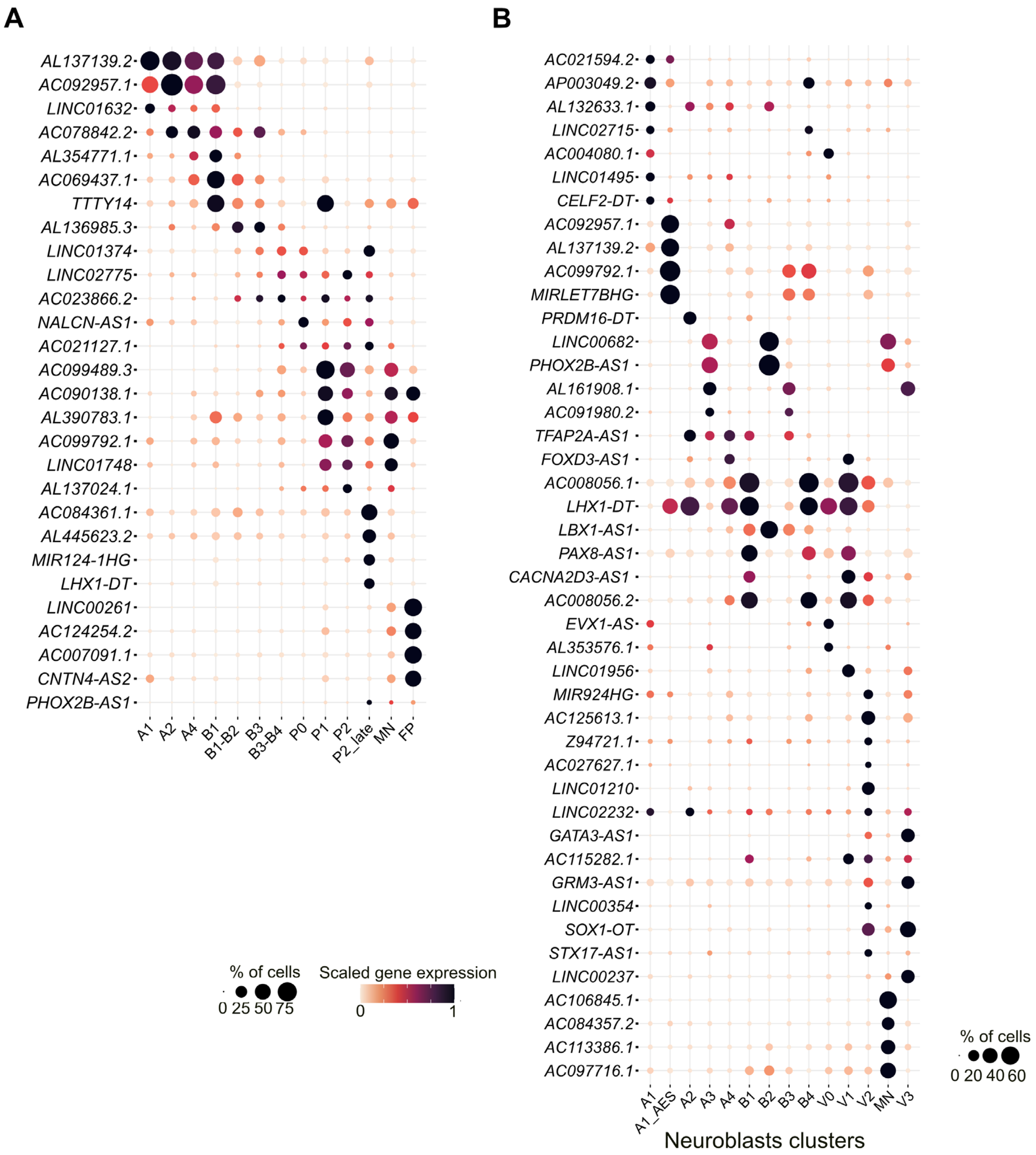


**Fig. S5: Novel non-coding markers of dorso-ventral progenitor and neuroblast domains in the developing pons and medulla.**

**A-B**) Expression of selected marker long-non coding RNAs for progenitor (**A**) and neuroblasts (**B**) domains identified in the developing hindbrain transcriptomic atlas. Neuroblasts clusters of the same domain are grouped together. Bubble size, proportion of cell population with expression. Color, scaled gene expression.

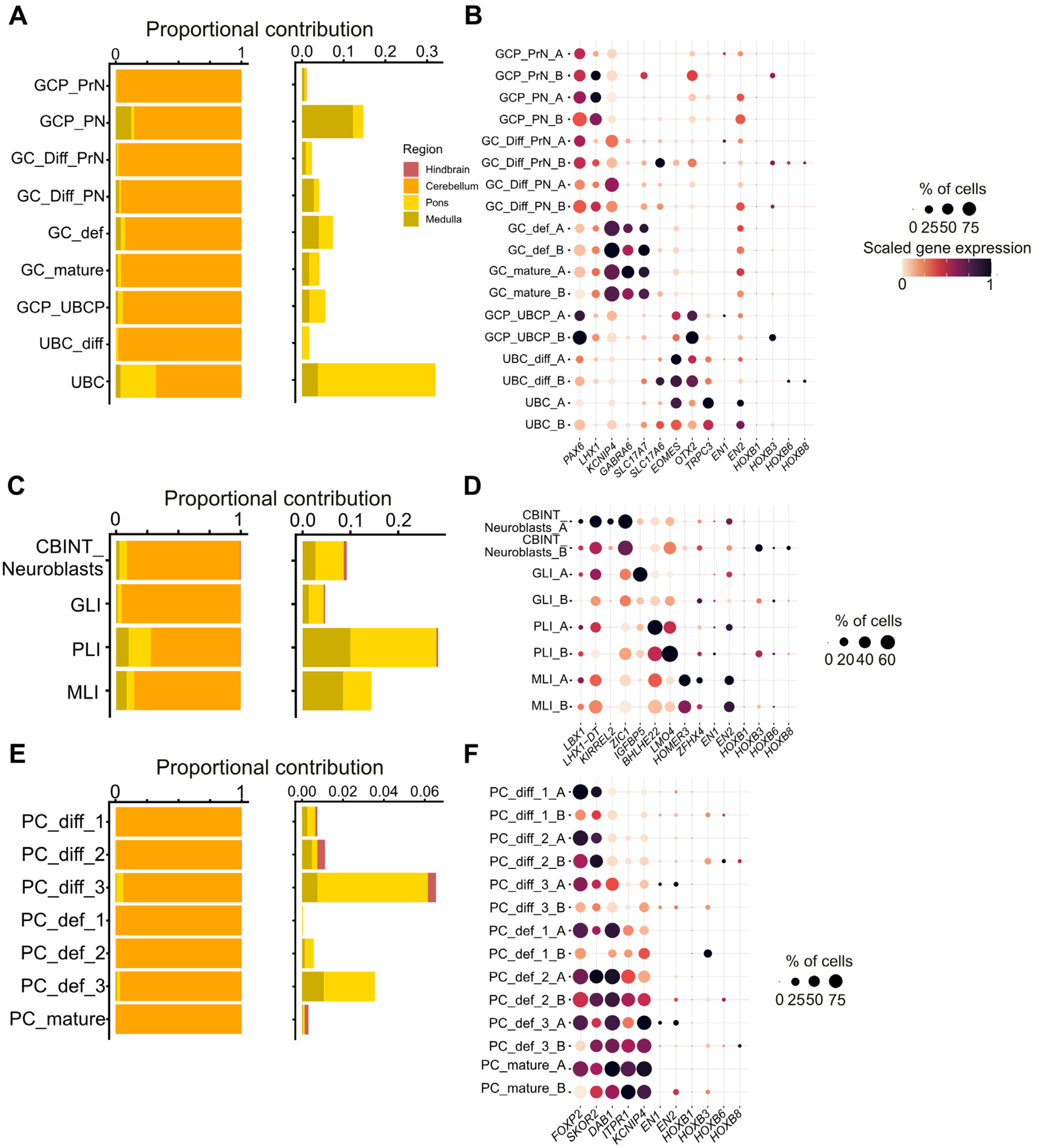


**Fig. S6: Distribution of cerebellar neuronal population in the pons and medulla.**

**A**) Proportional contribution from hindbrain, cerebellum, pons or medulla (left) or non-cerebellar contribution (left) to cell clusters of the granule cell and UBC lineage. **B**) Expression of selected marker for granule cell and UBC identity and axial identity (*EN1/2, HOX*) in cell types comprising the granule cell/UBC lineage. Clusters are grouped into cerebellar (_A) or non-cerebellar (_B) derived cells. **C**) Proportional contribution from hindbrain, cerebellum, pons or medulla (left) or non-cerebellar contribution (left) to cerebellar interneurons cell clusters. **D**) Expression of selected marker for cerebellar interneuron identity and axial identity in cerebellar interneuron clusters. Clusters are grouped into cerebellar (_A) or non-cerebellar (_B) derived cells. **E**) Proportional contribution from hindbrain, cerebellum, pons or medulla (left) or non-cerebellar contribution (left) to Purkinje cells. **F**) Expression of selected marker for Purkinje cell identity and axial identity in Purkinje cell clusters. Clusters are grouped into cerebellar (_A) or non-cerebellar (_B) derived cells.

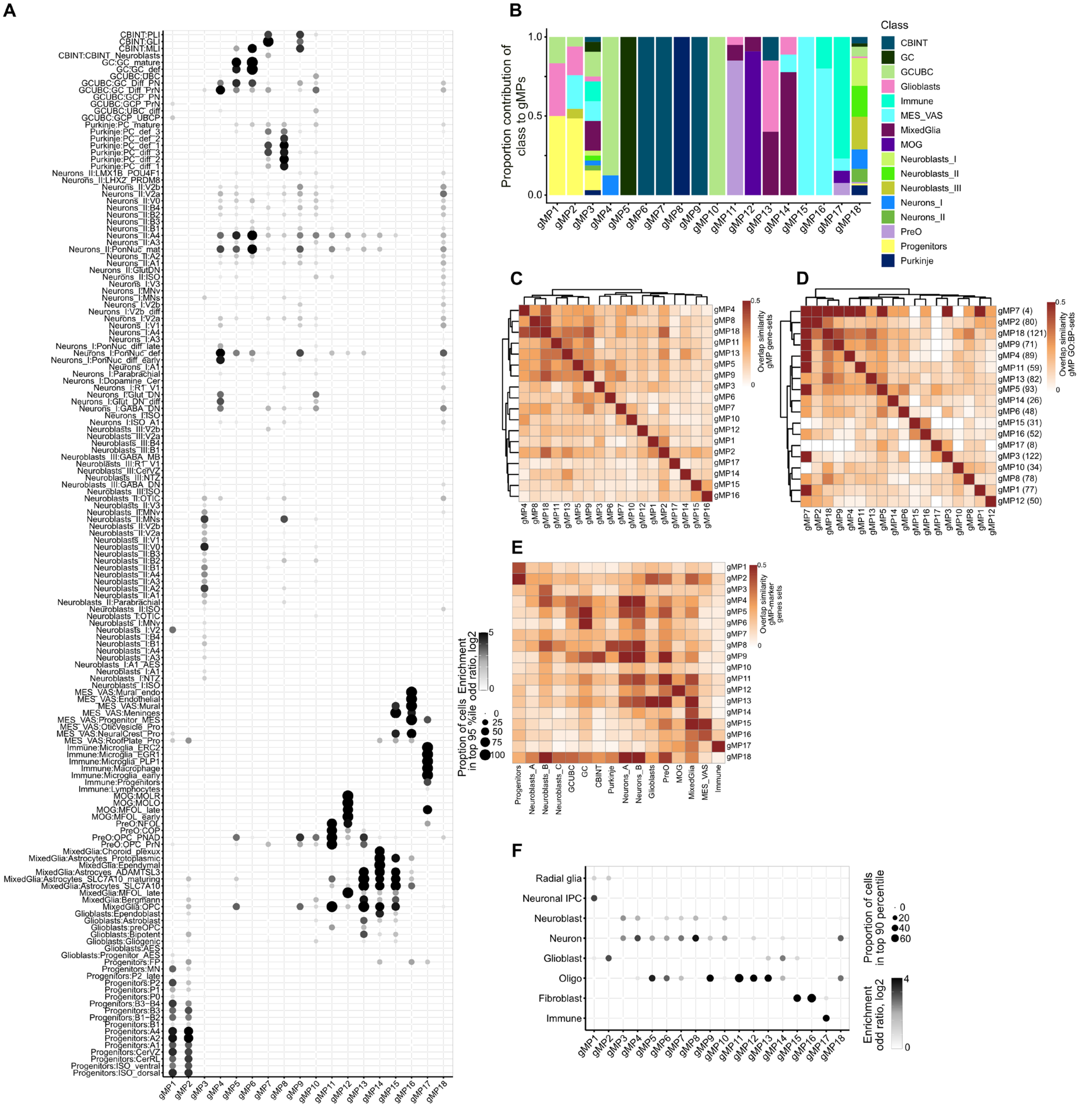


**Supplemental Figure S7** *legend next page* ►

◀*cont. from previous page*

**Fig. S7: Transcriptomic meta-gene program underlying developing human hindbrain cellular diversity.**
**A**) Enrichment of meta-gene programs gMP1-18 in individual cell populations grouped by cluster identity. Bubble size, proportion of cells in top 90th percentile of cells ranked by meta-gene program activity score. Color, $log_2$ enrichment odds-ratio. **B**) Proportional class contribution of constituting gene sets per meta-gene program. **C-D**) Overlap similarity between meta-gene programs in terms of representing gene sets (**C**) and associated GO:BP terms (**D**). Number in parentheses in D represent number of unique GO:BP terms per meta-program. **E**) Overlap similarity between meta-gene program gene sets (Table S6) and marker genes per class (Ext. Data 2, Zenodo) identified with pair-wise Wilcoxon test (fdr<1e-5 and AUC >0.65). **F**) Enrichment of meta-gene programs gMP1-18 in individual cell populations grouped by class identity from *Braun. et al*[18]. Bubble size, proportion of cells in top 90th percentile of cells ranked by meta-gene program activity score. Color, $log_2$ enrichment odds-ratio.

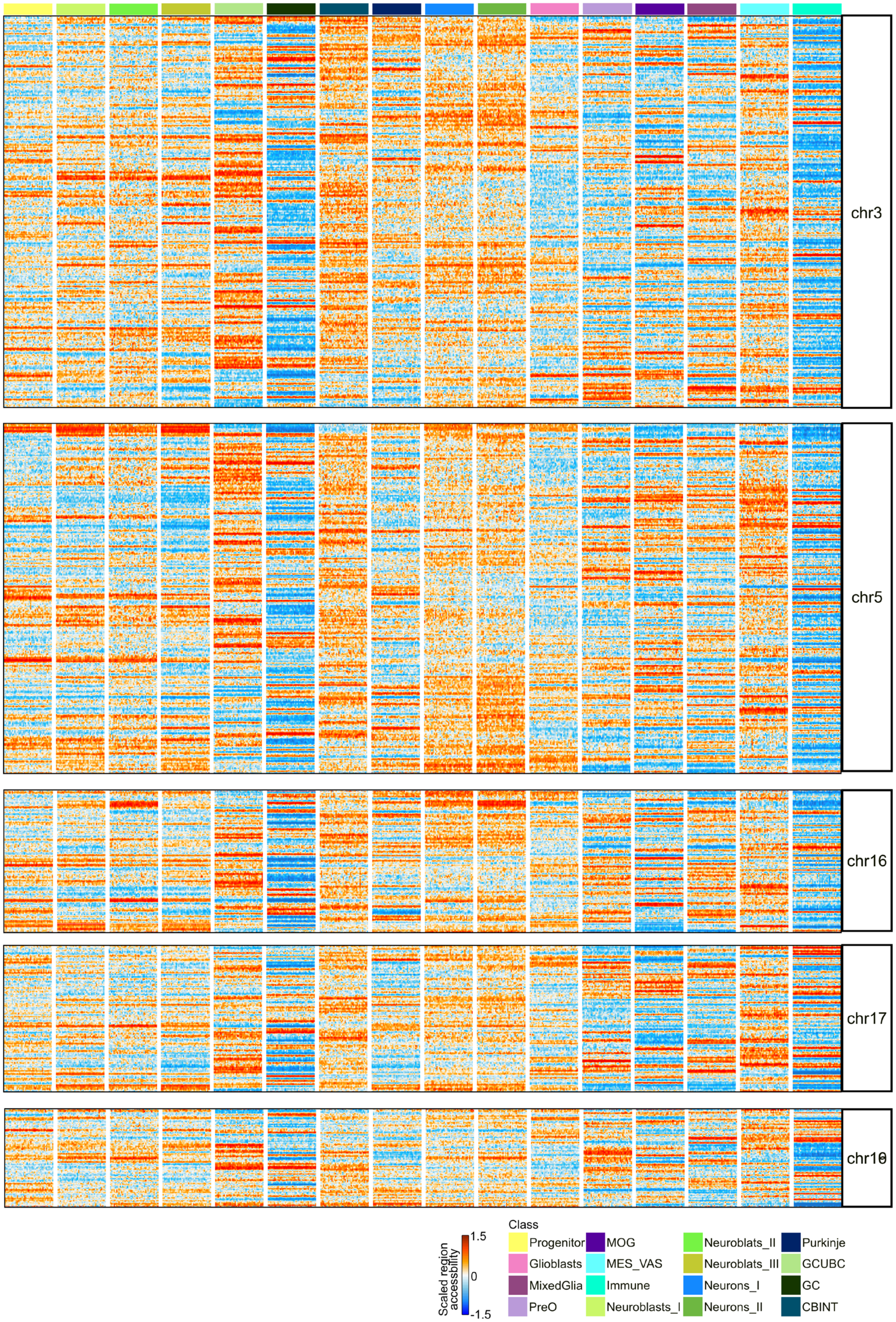


**Supplemental Figure S8** *legend next page* ►

◀*cont. from previous page*

**Fig. S8: Relative enrichment of chromosome accessibility per class.**
Heatmap depicting relative peak counts per mega-base-pair windows pseudobulked by 100 cells per class (subsetted to 4,000 cells) for a chromosome scaled across classes for representative chromosomes 3, 5, 16, 17 and 19.

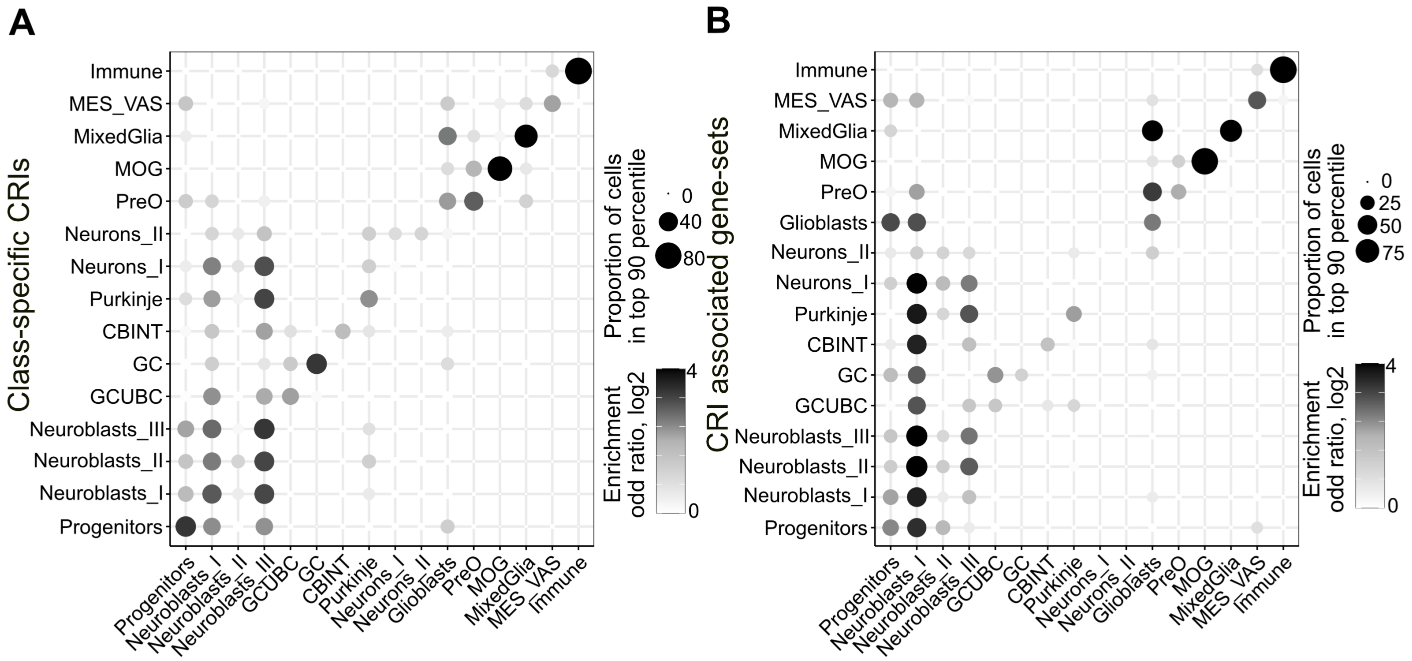


C

| Class | Observed Mean | Permutated Mean | p.value (permuta-tion) | p.value (Wilcox) | Gene count |
|---|---|---|---|---|---|
| MixedGlia | 0.281 | 0.174 | 1.00E-04 | 1.10E-06 | 149 |
| Progenitors | 0.232 | 0.179 | 8.00E-04 | 0.000019 | 142 |
| MOG | 0.285 | 0.186 | 1.00E-04 | 3.40E-04 | 157 |
| Immune | 0.23 | 0.157 | 0.0001 | 0.00055 | 231 |
| PreO | 0.22 | 0.182 | 0.029 | 0.0055 | 166 |
| Neurons_II | 0.209 | 0.176 | 0.043 | 0.033 | 139 |
| CBINT | 0.203 | 0.16 | 0.0051 | 0.061 | 181 |
| Purkinje | 0.252 | 0.185 | 2.10E-03 | 0.063 | 128 |
| GCUBC | 0.173 | 0.157 | 0.16 | 0.067 | 146 |
| Neuroblasts_III | 0.206 | 0.166 | 0.0071 | 0.088 | 160 |
| MES_VAS | 0.208 | 0.176 | 0.067 | 0.092 | 61 |
| Neuroblasts_II | 0.226 | 0.199 | 0.076 | 1.10E-01 | 147 |
| Glioblasts | 0.187 | 0.166 | 0.093 | 0.2 | 150 |
| Neuroblasts_I | 0.215 | 0.193 | 0.11 | 0.43 | 161 |
| GC | 0.155 | 0.163 | 0.64 | 0.5 | 113 |
| Neurons_I | 0.17 | 0.172 | 0.51 | 0.8 | 125 |

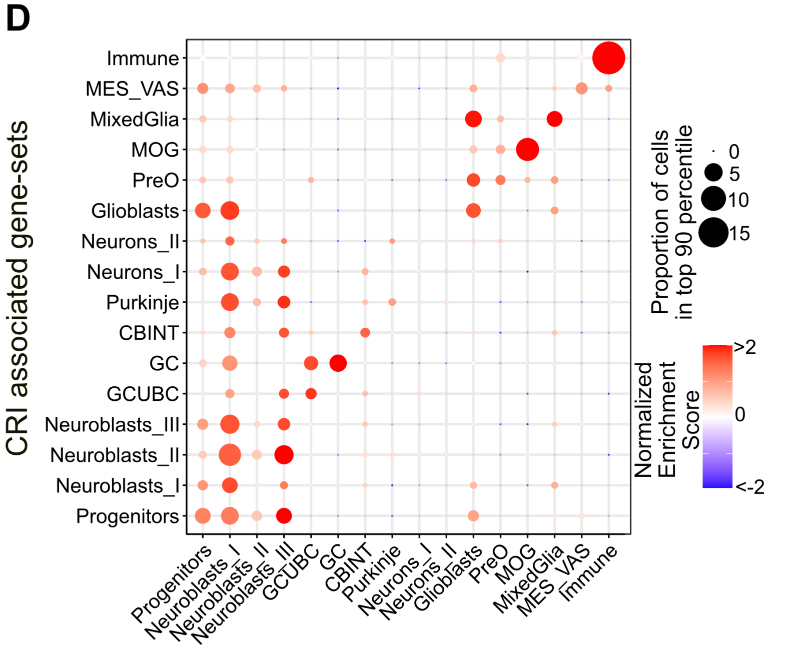


**Fig. S9: Cis-regulatory islands represent cell-type-specific regulatory activity driving cell-type-specific gene expression.**

**A**) Activity of class-specific cis-regulatory islands (CRIs) in cell populations of each of the cell classes. Bubble size, proportion of cells in top 90th percentile of cells ranked by CRE-set activity score. Color, $\log_2$ enrichment odds-ratio. **B**) Enrichment of CRI associated gene-sets in cell populations per class. Bubble size, proportion of cells in top 90th percentile of cells ranked by gene-set activity score. Color, $\log_2$ enrichment odds-ratio. **C**) Observed mean of CRI associated highly expressed (top 500) and highly variable genes (top 2000), compared to mean expression of 10,000 permutated gene sets of the same length among the class-specific highly expressed and highly variable genes. **D**) GSEA of CRI associated gene-sets in marker genes per class. Bubble size, adjusted p-value. Color, normalized enrichment score.

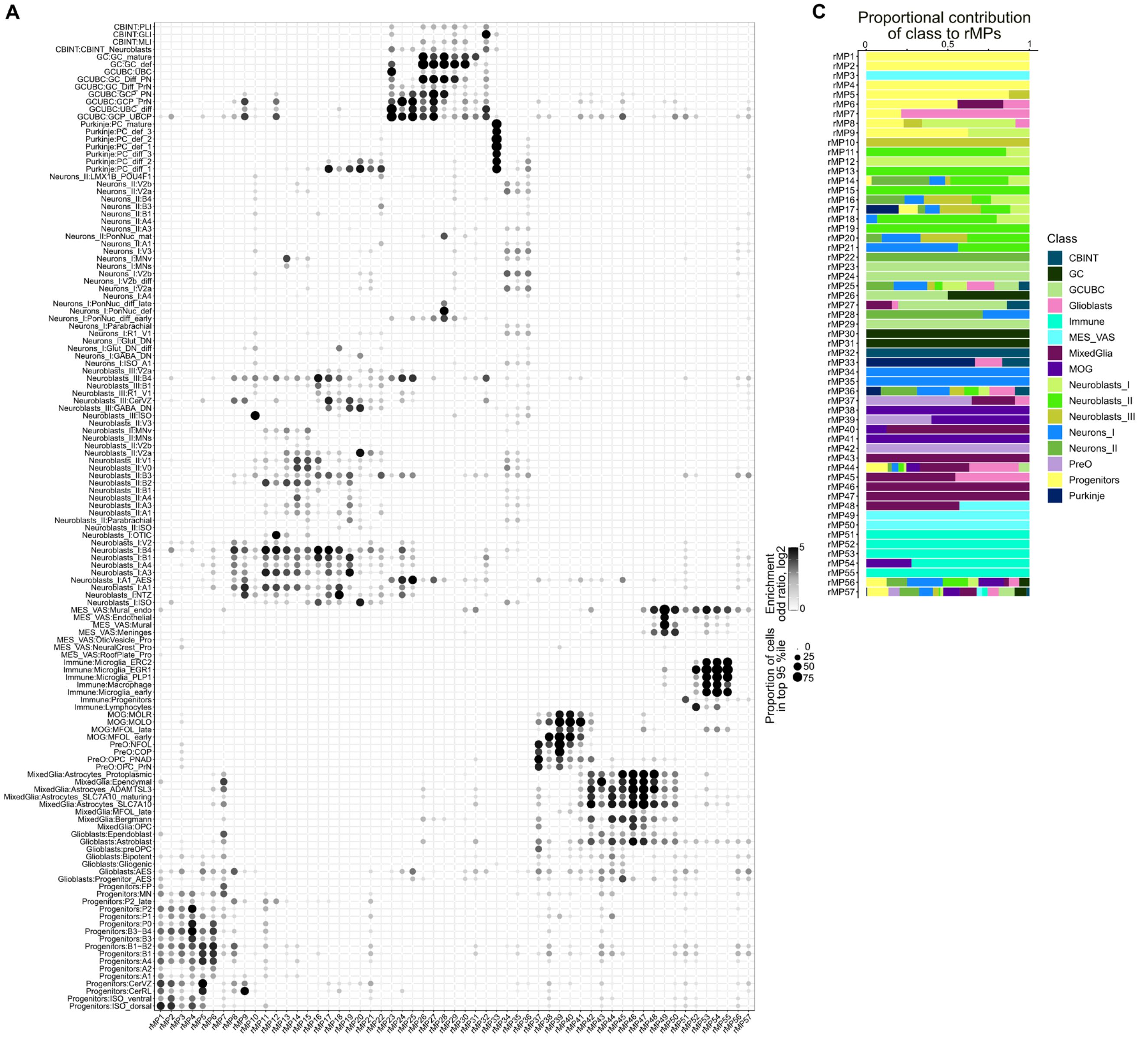


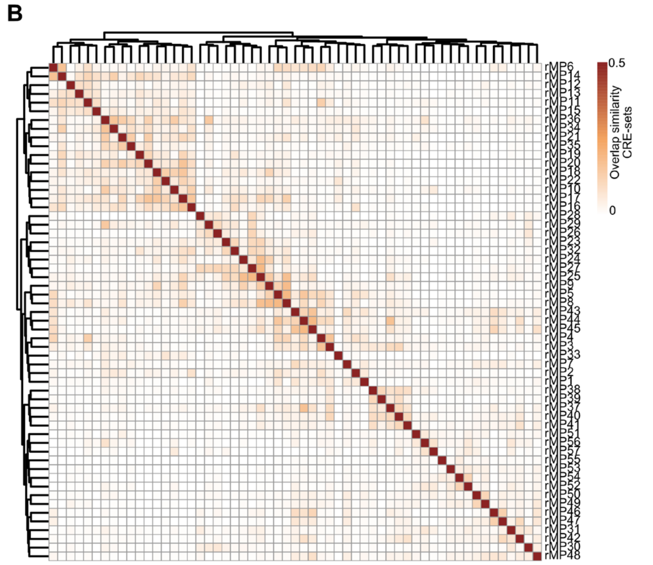


**Fig. S10: Cis-regulatory programs exhibit cell-type-specific enrichment.**
**A**) Enrichment of meta-regulatory programs rMP1-57 in individual cell populations grouped by their cluster identity. Bubble size, proportion of cells in top 95th percentile of cells ranked by meta-regulatory program activity score. Color, $\log_2$ enrichment odds-ratio. **B**) Overlap similarity between meta-regulatory programs in terms of representing CRE-sets. **C**) Proportion contribution of class derived CRE-sets per programs.

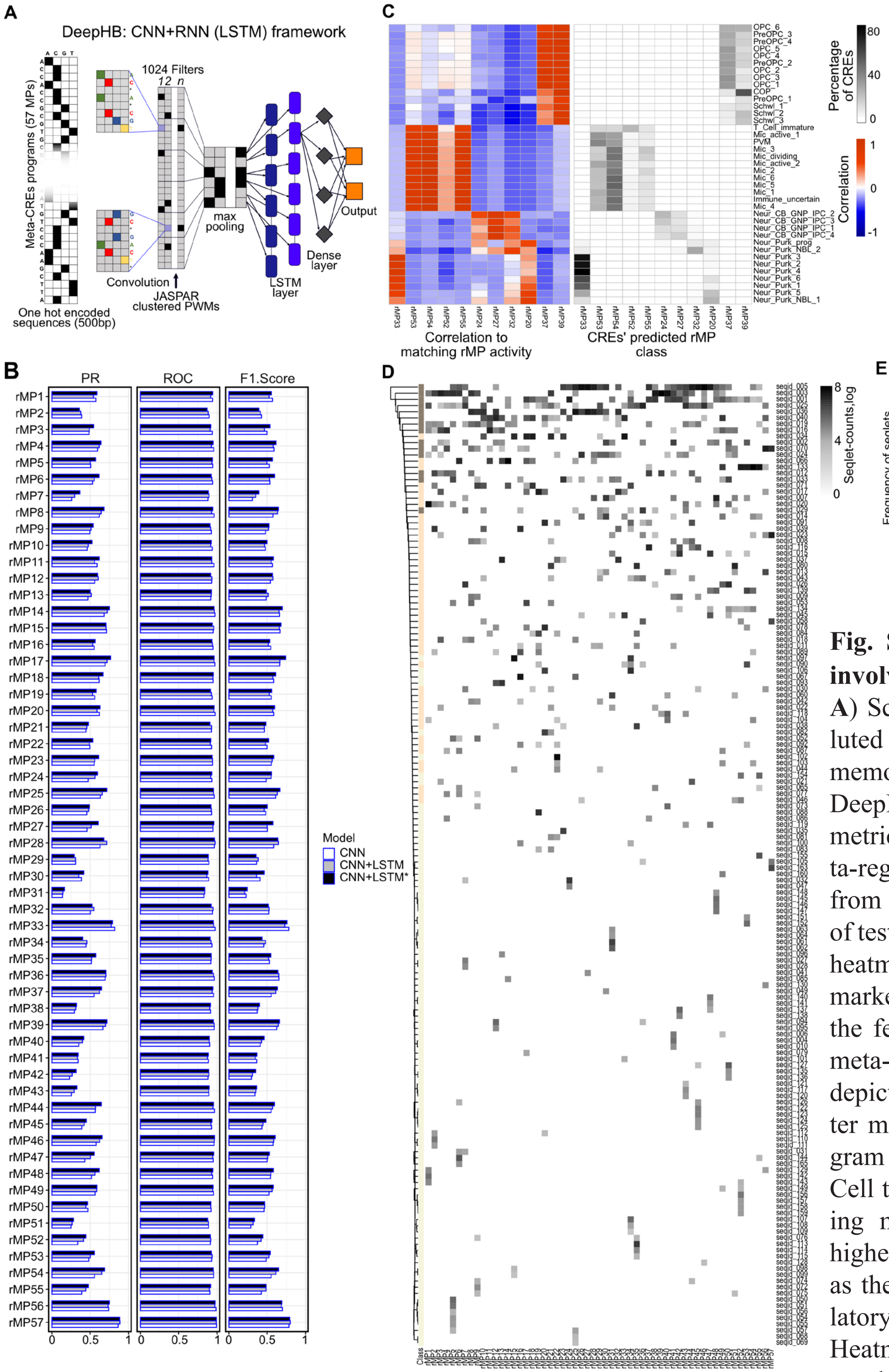


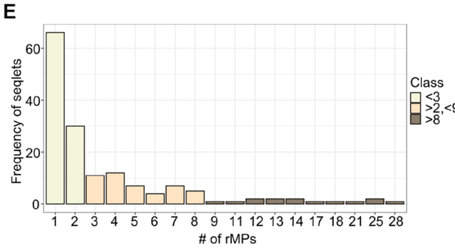


**Fig. S11: DeepHB, a classifier of CREs involved in hindbrain development.**
**A**) Schematic representing a hybrid convoluted neural network and long short-term memory (CNN+LSTM) model base for DeepHB. **B**) Barplot depicting evaluation metrics: auROC, auPR and F-score per meta-regulatory program (rMP1-57) obtained from the DeepHB model-based prediction of test-cohort CRE-sets. **C**) Left: Correlation heatmap between CRE-set activity score of marker CREs per cell cluster identified in the fetal brain atlas[19] (y axis) and selected meta-regulatory programs. Right: Heatmap depicting proportion of classified cell cluster marker CREs into meta-regulatory program classes (x-axis) using the DeepHB. Cell types enriched in both the atlas including microglia, oligo, Purkinje cells have higher number of markers CREs classified as the equivalent class enriched meta-regulatory program enriched in the hindbrain. **D**) Heatmap depicting seqlet counts per meta-regulatory program for all the observed 156 seqlets across 57 rMPs. **E**) Barplot depicting frequency of motifs (x-axis) shared across meta-regulatory programs (y-axis).

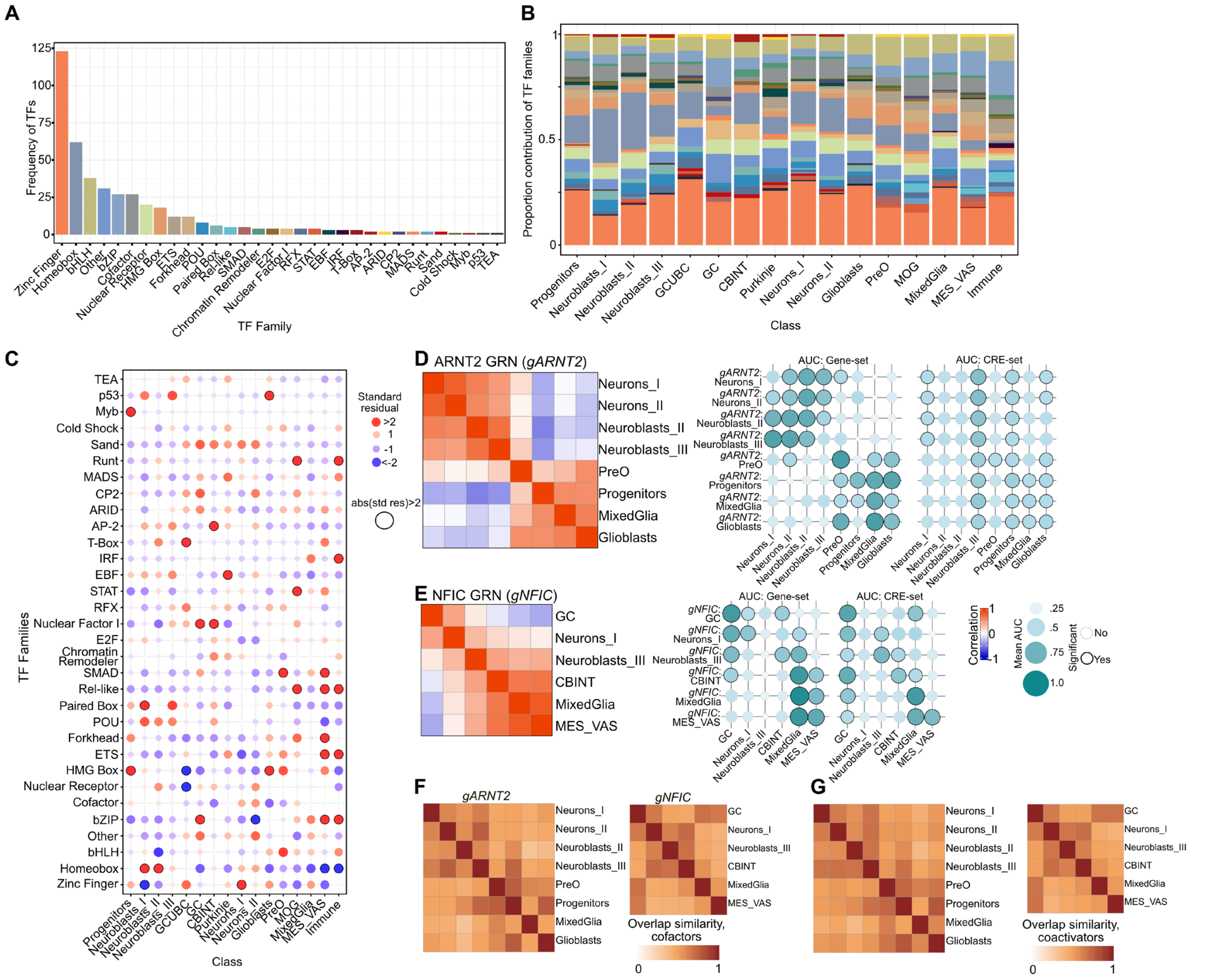


**Fig. S12: Transcription factor gene regulatory network (TF-GRN) activity driving hindbrain development.** **A**) Frequency of TFs per family detected as active across hindbrain cell classes. **B**) Proportional contribution of TF-families per class. **C**) Chi-square based enrichment of TF-families distribution per class. Size and color of bubble, residual obtained per class. Comparison with above absolute residual value of 2 are circled. **D**) Correlation heatmap of NFIC-regulated GRNs (*gNFIC*, top) in granule cells (GC), Neurons_I, Neuroblasts_III, cerebellar interneurons (CBINT), MixedGlia and mesenchymal/vascular (MES_VAS). Bubble plots showing pair-wise comparisons of *gNFIC* gene-set activity (bottom left) and CRE-sets activity scores (bottom right) in the selected classes. Bubble size and color represent mean AUC scores from pair-wise Wilcoxon tests. Significant enrichment circled. **E**) Correlation heatmap of ARNT2-regulated GRNs (*gARNT2*, left) in Neurons_I, Neurons_II, Neuroblasts_II, Neuroblasts_III, oligodendrocytes precursors (PreO), Progenitors, MixedGlia and Glioblasts. Bubble plots showing pair-wise comparisons of *gARNT2* gene-set activity (bottom left) and CRE-sets activity scores (bottom right) in the selected classes. Bubble size and color shows mean AUC scores from pairwise Wilcoxon tests. Significant enrichment circled. **F-G**) Heatmap depicting overlap similarity of co-factors (TFs binding to shared CREs, **F**) or co-activators (TF regulating same targets, **G**) for ARNT2 and NFIC GRNs across classes.

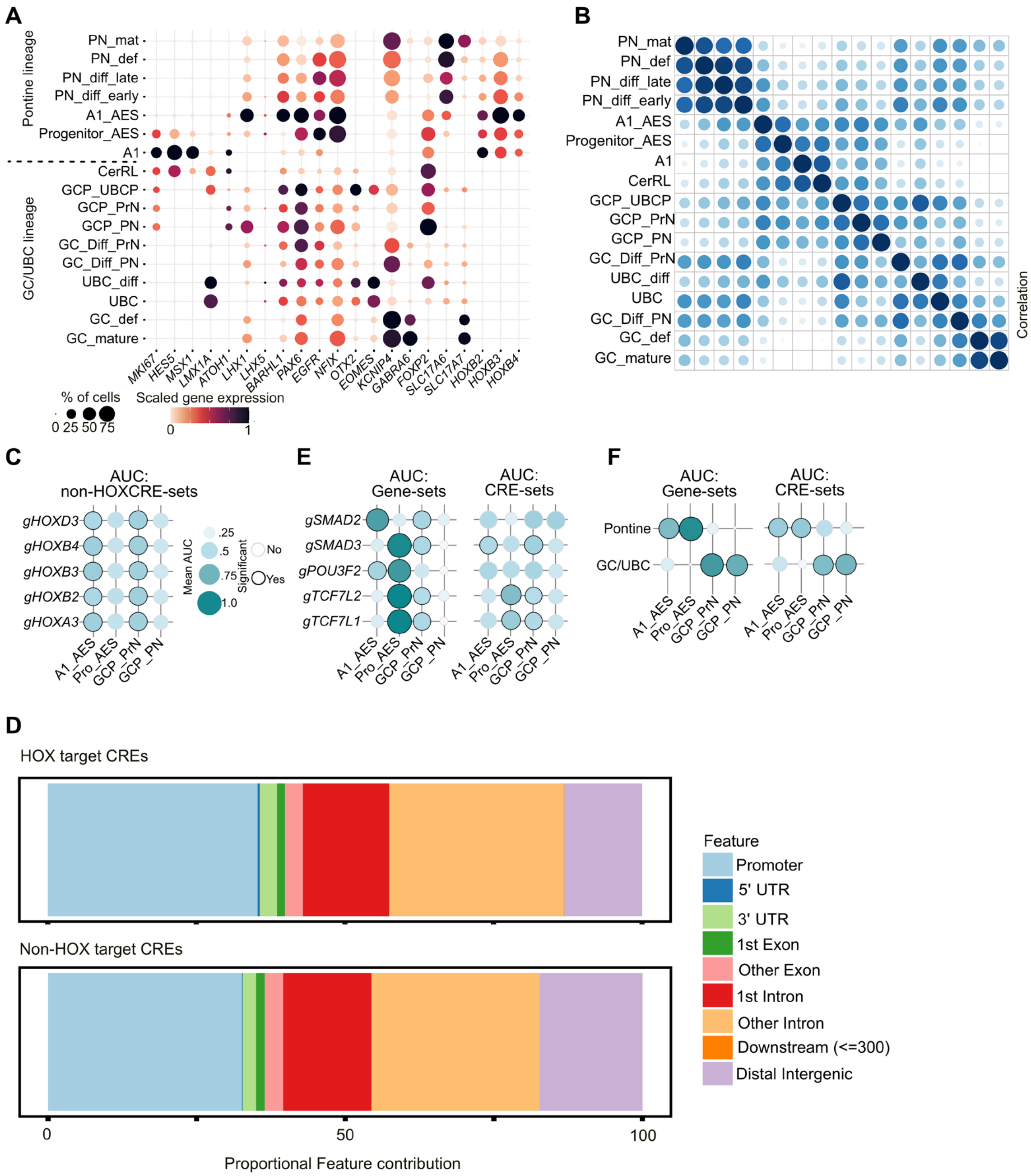


**Fig. S13: Upper and lower rhombic lip lineages in human hindbrain development.**
**A**) Expression of marker genes in cell clusters across upper (GC/UBC) and lower (pontine nuclei) lineages. Bubble size, proportion of cell population with expression. Color, scaled gene expression. **B**) Pearson correlations between cell cluster in upper and lower rhombic lip lineages. Correlation was calculated between log-normalized pseudo-bulked profiles of highly variable genes. **C**) Bubble plot depicting pair-wise comparisons of CRE-sets scores for non-HOX binding site for HOX targets in rhombic lip precursor clusters. **E**) Proportional distribution of genomic annotation for CREs associated with HOX TF-GRNs and non-HOX TF-GRNs identified in pontine nuclei lineage. **F**) Bubble plots showing pair-wise comparisons of gene-set activity (left) and CRE-set activity scores (right) for selected non-HOX TF-GRNS in rhombic lip precursor clusters. Bubble size and color shows mean AUC scores from pairwise Wilcoxon tests. Significant enrichment circled. **G**) Bubble plots showing pair-wise comparisons of gene-set activity (left) and CRE-set activity scores (right) for top 100 differentially upregulated genes in pontine or granule cell precursors; and their associated CREs as identified from TF-GRNs identified in pontine nuclei lineage. Bubble size and color shows mean AUC scores from pairwise Wilcoxon tests. Significant enrichment circled.

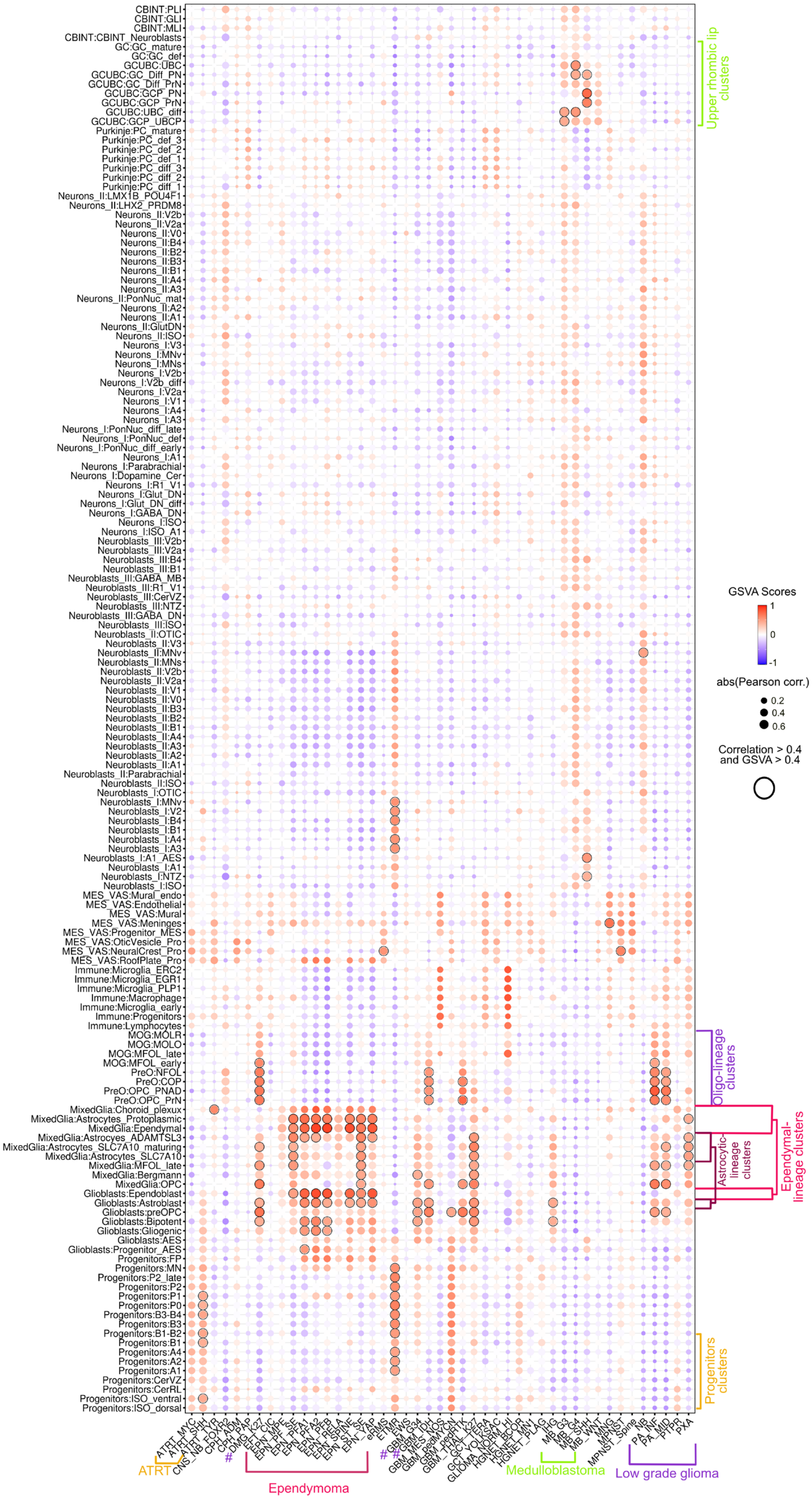


**Supplemental Figure S14** *legend next page* ►

◀*cont. from previous page*

**Fig. S14: Hindbrain reference identifies potential lineage of origin of pediatric brain tumors.**
Bubble plot showing GSVA-correlation analysis of transcriptomic signature between the hindbrain cell clusters (y-axis) and tumors grouped by subgroup identity (x-axis). Bubble size, Pearson correlation between expression of highly variable genes in reference and target. Color, GSVA enrichment values. Comparisons with correlation > 0.4 and GSVA-score > 0.4 are circled highlighting significant matches. Selected hindbrain cell clusters and associated tumors are highlighted in the same colors. Selected high-grade glioma are marked by #.

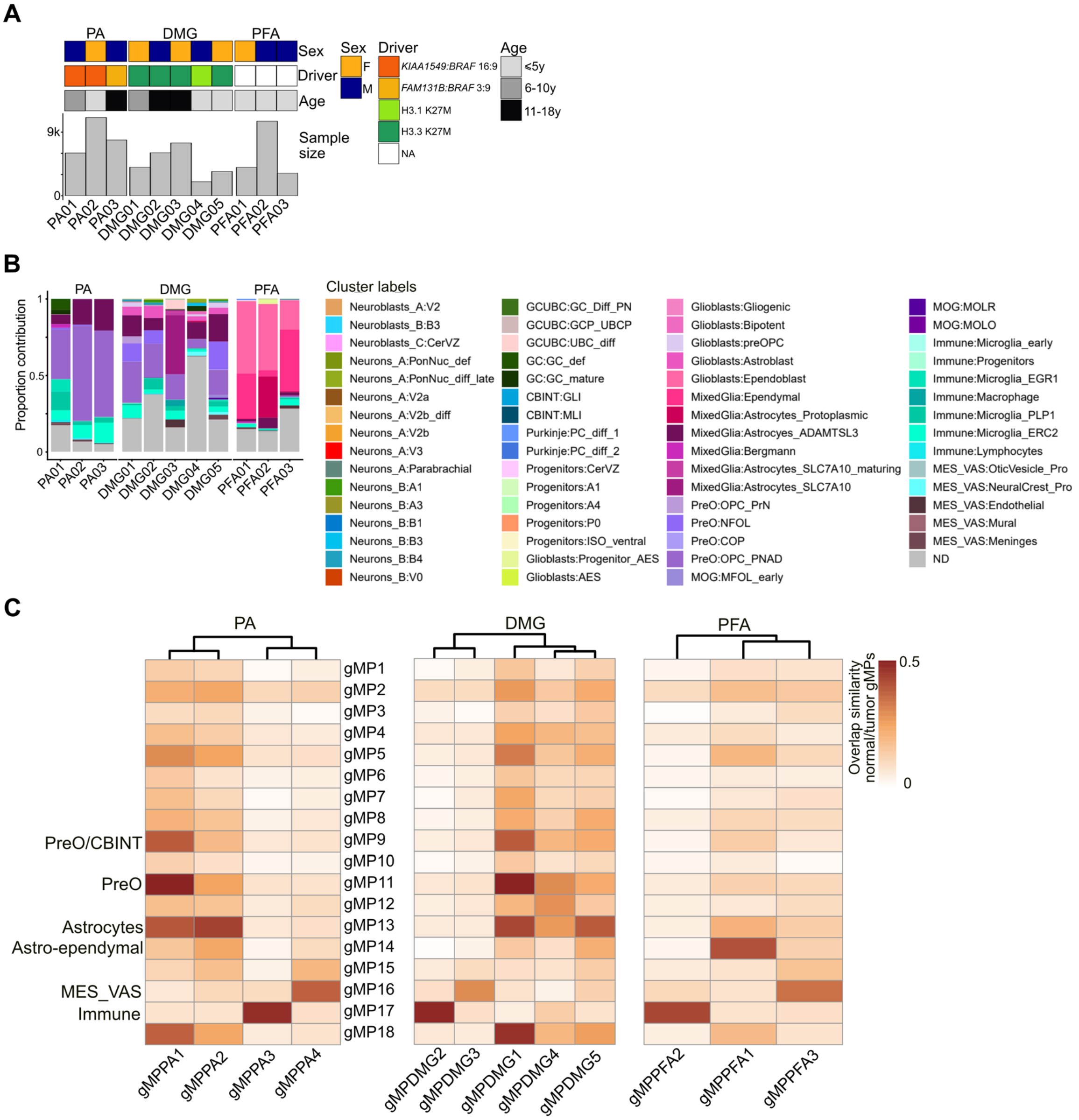


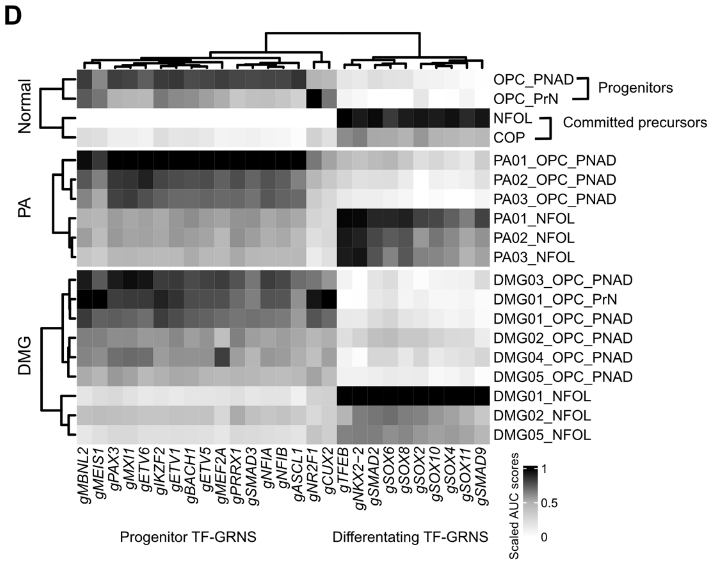


**Fig. S15: Conserved identity of lineage of origins in tumors.**
**A**) Pediatric glioma single-cell RNA-seq data cohort description. **B**) Proportional contribution of predicted cluster labels per cell per sample. Cells with less than 0.4 prediction scores are assigned “ND” identity (grey). **C**) Heatmap depicting Jaccard similarity between representative meta-gene programs identified in tumors (x-axis), including the tumor microenvironment compartment, and the best matching meta-gene program identified in the hindbrain atlas (y-axis) in pilocytic astrocytoma (PA), diffuse midline glioma (DMG), and posterior fossa ependymoma (PFA). **D**) TF-GRN enrichment heatmap for progenitors and committed precursors in normal oligodendroglia lineage, and equivalently assigned tumor cells PA and DMG. Selected TF-GRNs enriched in progenitors and differentiating normal oligodendroglia cells are shown.

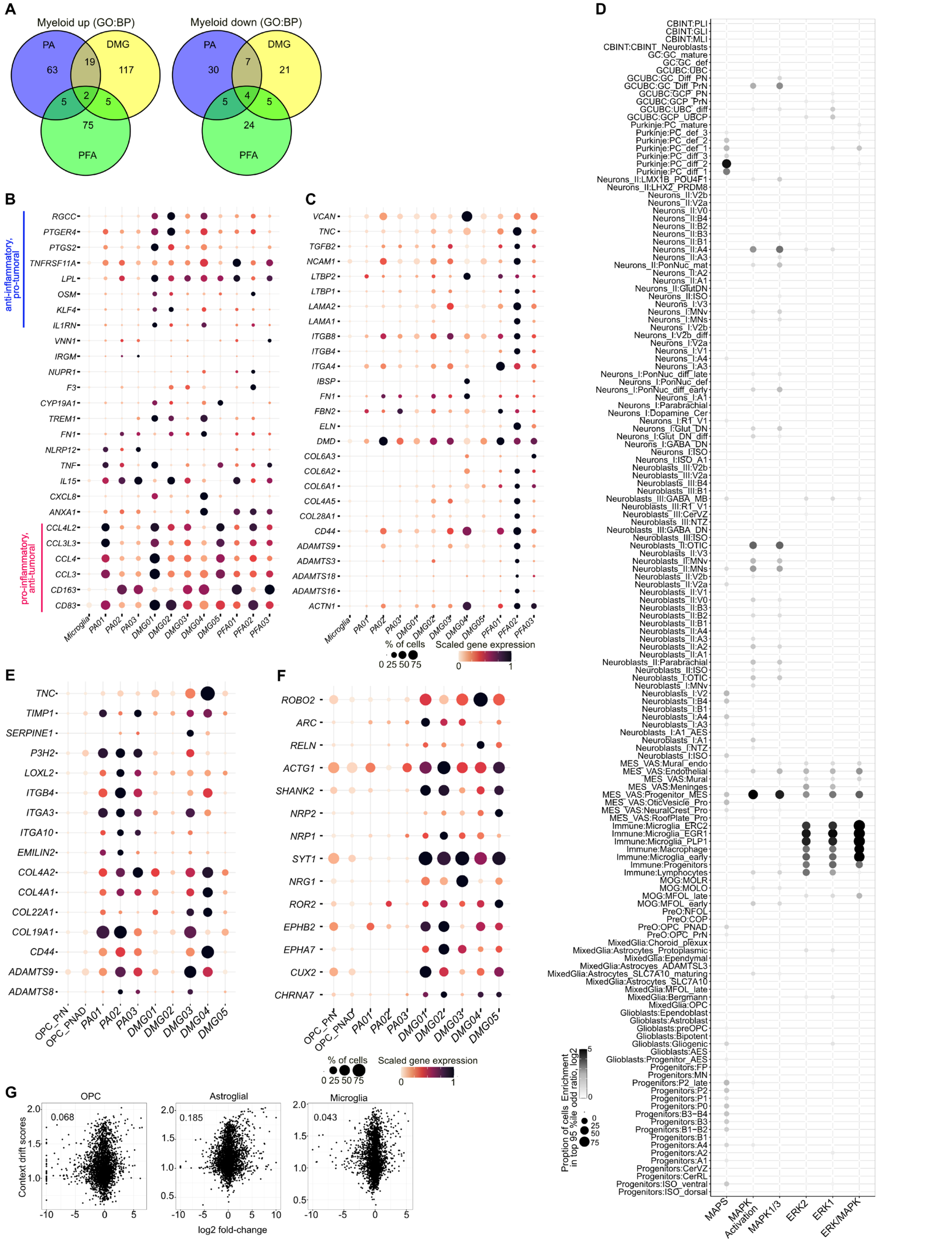


**Supplemental Figure S16** *legend next page* ►

◀*cont. from previous page*

**Fig. S16: Differentially upregulated gene sets in tumor cellular compartments.**

**A**) Venn-diagram depicting overlap of differentially up- (left) or down- (right) regulated genes in PA, DMG and PFA myeloid compartment compared to normal microglia. **B,C**) Expression plot depicting expression of differentially upregulated genes in tumor myeloid compared to normal microglia. Selected pro- and anti-inflammatory genes (**B**) and gene associated with extracellular matrix pathway (**C**) are shown. **D**) Bubble plots depicting enrichment of MAPKi sensitivity (MAPS) and MAPK kinase activation (R-HSA-450294), MAPK1/MAPK3 signaling (MAPK1/3, R-HSA-5684996) , MAPK1 (ERK2) activation (R-HSA-112411), MAPK3 (ERK1) activation (R-HSA-110056) and ERK/MAPK targets (R-HSA-198753) gene sets in clusters in human hindbrain. Bubble size, proportion of cells in top 95th percentile of cells ranked by gene set activity score. Color, $\log_2$ enrichment odds-ratio. **E,F**) Expression plot depicting expression of differentially upregulated genes in tumor oligo-compartment compared to normal OPC. Selected genes associated with extracellular matrix pathway (**E**) and synapse assembly pathway (**F**) are shown. **G**) Scatter plot for context drift scores (y-axis) and $\log_2$ fold-change (x-axis) of genes in oligo, astrocytic and microglial compartment in PA01. Pearson correlation between context drift score and $\log_2$ fold-change is mentioned for each comparison.

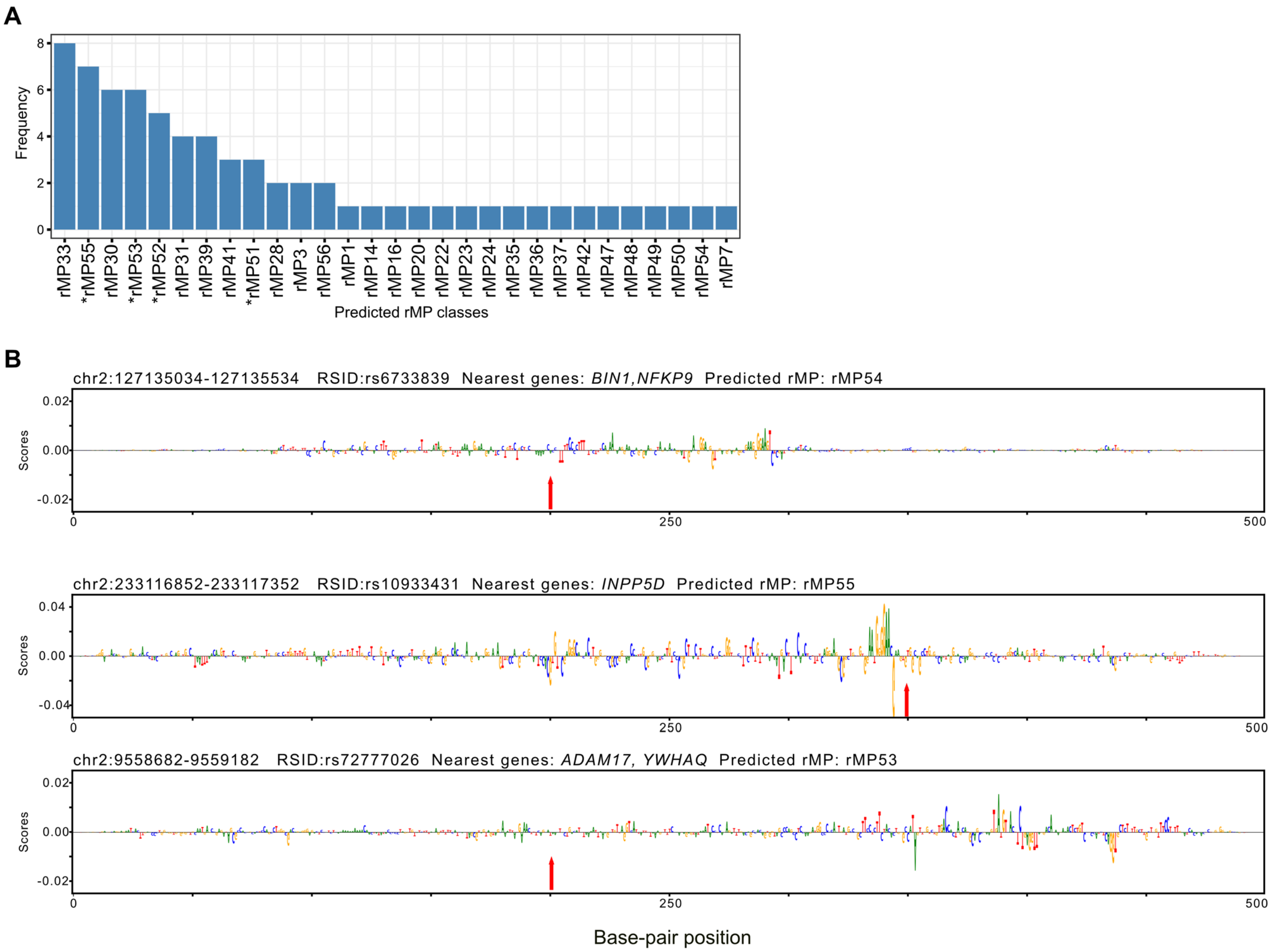


**Fig. S17: DeepHB based classification of putative enhancer elements.**
**A**) Barplot distribution of assigned meta-regulatory program class for 69 SNPs that were assigned to one of the 57 meta-regulatory programs using DeepHB classifier. Meta-regulatory programs enriched in Immune cell populations are marked with asterisk. **B**) Representative example of CRE segments (500 bp) predicted to be Immune enhancers. Alphabets represent contribution scores in terms of the best matching meta-regulatory program. Arrow marks the location of SNP.

### Supplemental Table Legends

**Table S1.** Cohort description for snRNA-seq and snATAC-seq data.
**Table S2.** Per library QC metrics for the snRNA-seq atlas.
**Table S3.** Per library QC metrics for the snATAC-seq atlas**.**
**Table S4.** List of class, cluster and subcluster in the annotated transcriptomic atlas.
**Table S5.** List of selected early and late DV, AP and neurotransmitter markers.
**Table S6.** Gene-sets per meta-gene program.
**Table S7.** TF-families of TFs associated with GRNs.
**Table S8.** Pediatric glioma snRNA-seq data cohort description.
**Table S9.** Context drift scores of genes in PA.
**Table S10.** Context drift scores of genes in DMG.

### Materials and Methods

#### Sample selection and ethics statement

No statistical methods were used to predetermine the sample size for either the normal hindbrain atlas or the pediatric glioma cohort. Sample size per stage was dictated by the availability of scarce tissue. Wherever possible, biological replicate data was generated from at least 2 donor tissues from either sex. The sample cohort for normal hindbrain is described in Table S1 and for pediatric glioma is described in Table S8. Experiments were not randomized and investigators had access to full metadata during data analysis.

For the normal hindbrain atlas, human prenatal samples were donated voluntarily to the MRC Wellcome Trust Human Developmental Biology Resource (HDBR; UK). The human postnatal samples were from the University of Maryland Brain and Tissue Bank of National Institutes of Health NeuroBioBank (USA), and Lenhossék Human Brain Program, Human Brain Tissue Bank at Semmelweis University (Hungary). All tissues were determined to be healthy through screening for typical pathogenic signatures at respective biobanks. Informed consent for the use of tissues for research was obtained in writing from donors or their family.

For the pediatric glioma cohort, frozen tumor tissues for pilocytic astrocytoma (PA) and diffuse midline glioma (DMG) were used. For the tumor tissue, informed consent was obtained from patients and/or parents or legal guardians according to International Cancer Genome Consortium (ICGC; for PA) or INdividualized Therapy FOr Relapsed Malignancies in Childhood (INFORM; for DMG) guidelines.

The use of human samples was approved by an ERC Ethics Screening panel (associated with H.K.'s ERC Consolidator Grant 615253, OntoTransEvol and S.M.P's ERC Advanced grant) and ethics committees in Heidelberg (authorization S-220/2017 and S-257/2017), North East-Newcastle & North Tyneside (REC reference 95 18/NE/0290), London-Fulham (REC reference 18/LO/0822), Ministry of Health of Hungary 96 (No.6008/8/2002/ETT) and Semmelweis University (No.32/1992/TUKEB).

#### Single-nuclei extraction, single-nuclei RNA-sequencing and single-nuclei ATAC-sequencing

Flash frozen samples were processed to extract nuclei as described in Sepp *et al.*[14]. Normal brain samples from prenatal and postnatal donors were first dissected into pieces using a surgical blade on dry ice. Dissected tissue was homogenized in the homogenization buffer (for details of reagents[14]*)* by trituration (prenatal) or douncing (postnatal) with a micro-pestle. For postnatal samples, lysate was filtered using a 70µm filter to remove bulk of the cellular debris. Homogenized tissue was centrifuged at 100g for 1 min to remove cellular debris, followed by two rounds of nuclei washing at 500g for 5 min. Nuclei obtained from postnatal donors, except from donor UMBN_6172, were then depleted for oligodendrocytes using Myelin Removal Beads II and LS MACS columns (Miltenyi Biotech). Eluted or washed nuclei were pelleted and re-suspended in 1x Nuclei buffer (10x Genomics) and strained through a 40µm filter to remove leftover debris and nuclei clumps. Nuclei were diluted 1:10 and stained with Hoechst DNA dye and propidium iodide to measure concentration on Countess II FL Automated Cell Counter (Thermo Fisher Scientific). Extracted nuclei were split into two vials and further processed for single-nucleus RNA-sequencing (snRNA-seq) and single-nucleus ATAC-sequencing (snATAC-seq). For snRNA-seq, nuclei were processed with Chromium Single Cell 3' v2.1/Next GEM Single Cell 3' v3.1 kit for and with ATAC v1.1 kits for snATAC-seq, as per manufacturer's recommendation. On average, 15,000-20,000 nuclei were loaded per channel along with the 3'/ATAC gel bead. DNA and cDNA libraries were prepared as described in respective kit protocols. Libraries were quantified using Qubit Fluorometer (Thermo Fisher Scientific) and profiled using Fragment Analyzer. GEX and ATAC libraries were sequenced using NextSeq2000 to recommended lengths and depth.

Tumor samples for PA or DMG were processed for nuclei extraction as described in Ernst *et al.*[80]. Post-nuclei extraction, PA samples were processed using Chromium Next GEM Single Cell 3' v3.1 kit for snRNA-seq and DMG samples were processed using Chromium Single Cell Multiome ATAC + Gene expression kit. Obtained cDNA libraries were sequenced as above. For ATAC-seq data generated from DMG, only 1 out of 5 samples passed QC parameters and thus these data were not included in the analysis.

#### snRNA-seq data processing and data integration

Normal brain and brain tumor snRNA-seq data was processed using the same pipeline. Demultiplexed sequencing reads were aligned to human genome assembly GRCh38 (v. p13, release 37, gencodegenes.org). Corresponding Gencode comprehensive gene annotation (PRI) was customized by filtering to transcripts with the following biotypes: protein coding, lncRNA, Immunoglobulin (IG) and T cell Receptor (TR) gene and pseudogene as recommended by *cellranger mkgtf* wrapper. Reads were aligned using

STARsolo[81] with parameters: --soloType *CB_UMI_Simple* --soloFeatures *Gene GeneFull* --soloUMIfiltering *MultiGeneUMI* --soloCBmatchWLtype *1MM_multi_pseudocounts* --soloCellFilter *None* --outSAMmultNmax *1* --limitSjdbInsertNsj *1500000*. In intronic alignment, reads assigned to overlapping genes are not counted leading to underestimation of certain genes' expression. For these overlapping genes, exonic alignment counts were used. Cells were separated from debris using *diem*[82] pipeline (*R*). Low-quality cells were filtered using a combination of following criterion: mitochondria fraction > 1 median absolute deviation (MAD) above the mean or above 2% (whichever was greater), number of detected reads (nUMIs) above 97.5%, and an intronic fraction (number of reads aligned to intron/total number of reads aligned to exon+intron) below 25%. Filtered cells were then corrected for background signal using *SoupX*[83] (*R*) and *decontXcounts()* (*celda*[84], *R*) pipelines. In the last step, putative doublets identified by *scrublet*[85] (*Python*) were removed. Filtered gene expression matrices were normalized using the *scran*[86] (*R*) approach.

For normal hindbrain atlas, log-normalized gene expression data from 107 libraries, representing 612,249 cells, were merged together using *mulitbatchnorm()* (*batchelor*[86], *R*). Highly variable genes (HVG) were obtained for the integrated data after removing mitochondrial (prefix: MT-), ribosomal genes (prefixes: RPS, RPL, MRPS, MRPL) and sex-chromosome-specific genes (chr X and Y). The integrated log-normalized data was sub-setted to top 4,000 HVG and scaled using *cosinenorm()* (*batchelor, R*) and factorized using non-negative matrix factorization (NMF) for a rank of 100 (*sklearn.decomposition.NMF(n_components=100, init='nndsvda', n_max_iter=10000*) to obtain a joint embedding. No batch correction algorithms were used to integrate the transcriptomic hindbrain atlas.

Tumor samples for PA/DMG/PFA were integrated by tumor type. Log-normalized gene expression data were merged together (*multibatchnorm(), batchelor, R*). HVG were first identified per sample after removing mitochondrial and ribosomal genes. Then a union of the top 1,500 HVGs per sample were obtained and sex-chromosome specific genes were removed from this obtained list to obtain a final HVG set per tumor type.

### Annotating the developing hindbrain atlas

Cells were algorithmically grouped in two-steps using the joint NMF embedding of the integrated data as input. Initial grouping was obtained from n-nearest neighbors (n=50) graph using Jensen-Shannon divergence (*NNDescent(),PyNNDescent, Python*) and leiden partitioning (*la.find_partition(la.ModularityVertexPartition, n_iterations=-1,max_comm_size=4000), leidenalg, Python*). Obtained cell clusters were then grouped (level 1 grouping) based on hierarchical clustering of pseudo-bulked gene expression profiles. This level 1 grouping represented an approximation of class level grouping. Cells assigned to same group at level 1 were further partitioned (level 2) using radial nearest neighbor (NearestNeighbors(n_neighbors=6, radius=0.5) graph with Jensen-Shannon divergence (*cdist(), scipy.spatial.distance, Python*) and leiden partitioning (*la.find_partition(la.ModularityVertexPartition, n_iterations=-1,max_comm_size=1000))*. Singleton and small sets (< 25 cells) obtained at level 2 grouping were excluded from annotation. Altogether this two-step clustering resulted in a total of 594,817 cells and 1482 cell clusters.

Partitioned cells grouped by shared transcriptomic signature were then manually annotated to assign a 'sub-cluster' level identity based on expression of cell-type-specific marker genes, stages and anatomical regions of contributing cells, and correlation to other cell partitions of the same group (level 1). 'Sub-clusters' were grouped into a 'Cluster' level hierarchy based on a high correlation with each other and a lack of distinguishing marker genes excluding sex-specific chromosomal genes. Finally, 'Clusters' were grouped into a 'Class' based on hierarchical clustering of 'Clusters' and if it belonged to the initial grouping (level 1). 'Clusters' belonging to neuroblasts/neurons were analyzed together to improve resolution of Neuroblasts_I/II/III or Neurons_I/II classes. Table S4 provides details of marker genes and annotation remarks.

### snATAC-seq data processing and data integration

Demultiplexed ATAC-seq reads were aligned to GRCh38 using Cellranger's *cellranger-atac* wrapper and processed downstream using *ArchR*[87] (*R*). Briefly, per sample, fragment files obtained post alignment were converted into arrow files (*createArrowFiles()*) using custom gene annotation (same transcriptomic annotation as used for snRNA-seq analysis) with a cut-off Transcription Start Site (TSS) enrichment of 3 and minimum 3000 fragments per cell. Putative doublets were identified by ArchR's *addDoubletScores()* and removed with *filterDoublets(filterRatio=1)*. Cells with high fragment counts, above 99 percentiles, were also filtered. Post-filtering, cells were embedded in latent-semantic indexing (LSI) space using *addIterativeLSI(),* grouped using *addClusters()* and visualized using UMAP embedding obtained through *addUMAP()*. Finally, a gene score matrix was obtained using *getMatrixFromProject()*. Individual clusters obtained post-processing were checked for aberrant gene score profile of marker genes, low number of expressing cells but high mean value or vice-versa, and were manually filtered out.

After sample wise pre-processing of snATAC-seq data, arrow files from all 84 libraries were integrated by creating a combined *ArchR* project resulting in 424,533 cells. A joint LSI embedding was obtained using *addIterativeLSI()* and which was then used for joint UMAP embedding using *addUMAP()* and cell partitioning using *addClusters()*. A combined gene score matrix for integrated data was also obtained by combining gene score matrix from individual samples. Similar to snRNA-seq analysis, obtained cell clusters were grouped together based on similarity of marker gene score profile to obtain level 1 grouping. Cells grouped together at level 1 were then partitioned again using *addClusters()* to obtain level 2 clustering. Singleton, small sets (< 25 cells), and cell partitions with aberrant gene score profiles were removed from further analysis, resulting in a total number of 422,850 cells.

**Annotating the hindbrain chromatin accessibility atlas**

Cells in the chromatin accessibility atlas were assigned a 'Cluster' level identity in a two-step process using the transcriptomic atlas as a reference. First, labels were transferred from the integrated transcriptomic atlas to the integrated chromatin accessibility atlas using *ArchR's* implementation of Seurat canonical correlation analysis[27] (CCA) (*addGeneIntegrationMatrix*()) with the top 3000 HVG identified in the integrated RNA-seq atlas. This approach uses gene score profiles of snATAC-seq and gene expression profiles of the reference transcriptomic atlas to transfer labels from the reference to equivalent cell-states in the target using CCA. Then, each ATAC cell cluster (level 2, as obtained above) were assigned the identity of the most frequent 'Cluster' and the assignment was manually validated by comparing marker gene expression profiles obtained in the RNA-seq atlas to the gene score profile of the snATAC-seq atlas. This step yielded a 'Class' level separation of cells in the snATAC-seq atlas. To improve 'Cluster' identity of individual cells, label transfer was performed again between subsetted data using *addGeneIntegrationMatrix*() and a list of the top 2000 group-specific HVGs. Cells were mostly grouped by 'Class', but, as cells belonging to either Neuroblasts_I/II/III, Neurons_I/II, Glioblasts/MixedGlia or GC/GCUBC exhibited comparable gene expression profiles, cells assigned to these classes were grouped together for the second round of label transfer to improve annotation. The 'Cluster' level annotation obtained through this two-step process was verified by comparing correlations between pseudo-bulked gene expression and gene score profiles between 'Clusters' belonging to a 'Class' as shown in Fig. S3.

Of note, three other independent approaches were explored for label transfer including scJoint[88] (supervised and unsupervised), MaxFuse[89] and scGlue[90], both at integrated and 'Class' level label transfer. 'Cluster' labels obtained from these approaches were judged inferior or equivalent when compared to those obtained using CCA (as described above), based on correlation between pseudo-bulked gene expression and gene score profiles.

**Peak calling for the integrated snATAC-seq atlas**

Using the integrated data, first a minimum of 100 cells and a maximum of 5000 cells per group (level 1 clusters), with a sampling ratio of 0.8, were used to generate pseudobulk replicates via *addGroupCoverages().* Then peaks were identified using MACS2[91] caller via *addReproduciblePeakSet()* and peaks with reproducibility less than 4 were removed. Further, peaks that were detected in less than 3% of all 'Class' populations or peaks overlapping blacklisted regions were further filtered to obtain a set of 446,510 peaks.

**Transcriptomic meta-gene programs**

Meta-gene programs were obtained per class through a modified version of the approach used in Kinker *et al*[92]. Annotated cells in the transcriptomic atlas were first randomly divided into two approximately equal parts and processed similarly. Gene expression data per-class per-split was subsetted to the top 2000 HVG per class and max 3000 cells per cluster. The gene expression matrix was then factorized using Latent Dirichlet Allocation (*LatentDirichletAllocation(),sklearn, Python*) for ranks from 3 to 30 in increments of 3, in total obtaining 165 factors per class and split. Each latent factor was represented by the top 100 genes ranked by contribution per factor. In total 5,280 gene sets were used as input to obtain a representative set of meta-gene programs.

The Jaccard and an overlap similarity (size of intersection/ size of the smallest set) score was obtained for each factor when compared to all other factors. Representative meta-gene program gene sets were obtained in a two-step process.

First, non-redundant robust representative gene sets

were obtained per class from all genesets obtained from both of the splits. Robustness was measured by overlap of a gene set with other gene sets obtained from independent runs of a different rank. Redundancy was measured by overlap with gene sets obtained from the same rank. For each rank, gene sets from both splits were ranked by their maximum overlap with all the factors obtained for other ranks. Potentially noisy gene sets with overlap of less than 20% were discarded. Then, the gene sets were ranked by their maximum overlap with each other, and sets with 50% or more overlap to a higher ranked gene set were discarded to remove redundancy. This was performed for all ranks for a class. Then, redundancy was removed across ranks from the selected gene sets by ranking them by their maximum overlap among each other and removing sets with 50% or more overlap with a higher ranked gene set. A union of such robust non-redundant gene set was obtained from runs for all classes resulting in 307 gene sets.

In the next step, the obtained gene sets were grouped into meta-gene programs using hierarchical clustering and an initial set of 40 programs using *cutree()* (*hclust,R*) was identified. Each meta-program was represented by a set of the top 100 genes ranked by a combination of the number of times a gene appeared in constituting gene sets and an average rank of the gene (by contribution) across these sets. The initial 40 metaprograms were recursively merged with another meta-program based on overlap similarity, until all meta-programs had less than 33% overlap with each other, resulting in a final set of 18 meta-gene programs represented by gMP1 to gMP18.

**Gene set activity score enrichment analysis**

For individual gene sets, a module score, representing the gene set activity score, per cell was calculated using *addmodulescore()* (*Seurat*, *R*). Enrichment of the gene set activity score in a cell population was then evaluated through mean $\log_2$ odds-ratio when compared to the rest of the population, and the proportion of cells in top 90th (class level comparison for whole atlas or cluster level comparison involving only selected classes) or 95th (cluster level comparison of whole atlas) percentiles. Cells were down-sampled to a maximum of 9000 cells per class or 200 cells per cluster in comparisons involving the entire atlas. In tests involving only selected classes, cells were subsetted to a maximum of 1000 cells per cluster.

**Gene ontology and pathway term overrepresentation enrichment analysis**

Overrepresented biological function (GO:BP) and reactome pathway terms per meta-gene program were obtained using *gproflier2*[93] (*R*). Obtained terms were ranked by their p-value, and all ancestors and descendants of a highly ranked term were discarded if present to obtain a final set of biologically unique terms.

**Marker cis-regulatory features**

Marker peaks per cell groupings were obtained using *getMarkerFeatures()* (*ArchR*). For marker peaks per class, a cut-off FDR < 0.01 and Log2FC>=2 or AUC >0.52 was used. Filtered peaks were ranked by FDR and a maximum of top 10,000 peaks per class were used. For marker peaks per cluster, a cut-off FDR < 0.05 and Log2FC>=2 or AUC >=0.55 was used and a maximum of top 500 peaks ranked by FDR were used per cluster.

Marker peak distribution across non-sex-specific chromosomes was obtained per class. Enrichment of a particular chromosome across classes was evaluated via chi-square test (*chisq.test(), stats, R*).

**Cis-regulatory islands**

Spatially clustered CREs constituting cis-regulatory islands were identified using a modified ROSE (rank ordering of super-enhancers)-based super-enhancer analysis approach[44]. Briefly, cells in the snATAC-seq atlas were split into two approximately equal parts. Cells per-class and per-split were subsetted to a maximum of 3000 cells per cluster. BAM files per sample were de-duplicated, sorted and indexed with *samtools.* BAM reads per class and split were extracted from each sample and merged together to obtain a BAM file per class and split. Peaks were called per BAM file using *MACS3* command *macs3 callpeak -f BAMPE -g hs -B -q 0.001.* Peaks obtained from the integrated snATAC-seq atlas that overlapped with *MACS3* identified peaks were used as input for ROSE-based super-enhancer analysis.

ROSE analysis was performed per class and split BAM. BAM files with reads for randomly selected 10,000 cells from the snATAC-seq atlas was used as the control input. An intersect of super-enhancers identified for both splits of a class were used as super-enhancers or cis-regulatory islands for the class. Cis-regulatory islands were ranked by their signal intensity and a maximum of 100 cis-regulatory islands was used per class.

Peaks identified from the integrated analysis overlapping each of the obtained cis-regulatory island windows comprised the cis-regulatory island peak-set.

Topologically associating domains (TADs) in

developing human cortex were obtained from *Won et al.*[94]. For classes with proliferative cells (Progenitors, Glioblasts, Neuroblasts_I/III, PreO, MES_VAS and GCUBC), TADs associated with germinal zones were used, and for rest, TADs associated with the cortical and subcortical plate was used. Genes located in the same TAD as the candidate cis-regulatory island comprised the cis-regulatory island gene set.

For cis-regulatory island gene sets, gene set activity score enrichment analysis was performed as described above. Gene set enrichment analysis (GSEA) of cis-regulatory island gene sets in marker gene per class was performed using *fgseaMultilevel()* (*fgsea*[95]*,R*). A permutation test was used to evaluate the significance of expression of class-specific cis-regulatory island gene sets compared to other highly expressed genes in that class. A union of the top 500 highly expressed and top 2,000 HVGs was obtained per class and cis-regulatory island gene set was subsetted to overlapping genes. 10,000 permutated control gene sets of the same length as of the subsetted cis-regulatory gene set were obtained from the union of highly expressed and highly variable genes, not including cis-regulatory island associated genes. A one-sided Wilcoxon-test was used to obtain the p-value comparing mean expression value of the permuted gene sets to that of the cis-regulatory gene set and p-values obtained for all classes were adjusted using *p.adjust()* (*stats,R*) to obtain corrected p-values.

**Chromatin accessibility meta-regulatory programs**

Meta-regulatory programs per class were obtained through an equivalent approach as described for the meta-gene programs. Annotated cells in the snATAC-seq atlas were divided into two approximately equal parts, subsetted to a maximum of 3000 cells per cluster for each class and split and processed similarly. Peak matrix per class and per split was used to create a cisTopic object (*pycisTopic*[47]*, Python*) and processed further for topic analysis for ranks ranging from 10 to 75 in increments of 5, resulting in 595 topics per class and split. Each topic was represented by top 2000 peaks ranked by contribution. In total 19,040 peak-sets were used as input to obtain a representative set of cis-regulatory meta-regulatory programs.

The Jaccard and an overlap similarity score was obtained for each topic peak-set when compared to all the others topics. Representative peak-sets were obtained in a two-step process.

First, non-redundant robust representative peak-sets were obtained per class from all peak-sets obtained for all ranks and both splits. Robustness was measured by overlap of a peak-set with other peak-sets obtained from independent runs of a different rank. Redundancy was measured by overlap with peak sets obtained from the same rank. For each rank, peak-sets from both splits were ranked by their maximum overlap with all the other ranks, and peak sets with overlap less than 20% were discarded. Then, selected peak-sets were ranked by their maximum overlap among each other, first removing peak sets with less than 33% overlap with any other peak set of the same rank, and then discarding sets with 33% or more overlap with a higher ranked ranked-set were to remove redundancy. This was performed for all ranks for each class. Then, redundancy was removed across ranks for a class, first ranking them by their maximum overlap among each other and removing sets with 66% or more overlap with a higher ranked peak set. A union of such robust non-redundant peak set was obtained from runs for all classes, resulting in 1194 peak-sets.

In the next step, the obtained peak-sets were grouped into meta-regulatory programs using hierarchical clustering and an initial set of 75 programs using *cutree()* (*hclust,R*) was used. Each meta-program was represented by a set of the top 2500 peaks ranked by a combination of the number of times a peak appeared in constituting peak-sets and an average rank of the peak (by contribution) across these sets. The initial 75 meta-regulatory programs were recursively merged with another meta-program based on overlap similarity, until all meta-programs had less than 15% overlap with each other, resulting in a final set of 57 meta-regulatory programs represented by rMP1 to rMP57.

**Peak set activity score**

Individual peak-sets activity scores were calculated using score *AUCell_run()*(*AUCell*[96], *R*). Enrichment of peak-set activity in a cell population was then evaluated through a mean log-odds ratio when compared to the rest of the population, and the proportion of cells in top 90th (class level comparison for whole atlas or cluster level comparison when done in selected classes) or 95th (cluster level comparison of whole atlas) percentiles. Cells were down-sampled to a maximum of 9000 cells per class or 300 cells per cluster in comparisons involving the entire atlas.

**DeepHB classifier**

We used the *CREsted*[45] (*Python*) framework to train classifiers for the 57 meta-regulatory programs (as obtained above), and benchmarked three approaches: a convoluted neural network (*deeptopic_cnn()*,CNN), a hybrid convoluted neural network-long short-term memory networks (*deeptopic_lstm()*,CNN+LSTM), or a hybrid CNN+LSTM (CNN+LSTM*) initialized with 159 motifs obtained from clustering non-redundant JASPAR CORE Vertebrates motif collection[97]. First, the peak-sets representing each of the 57 meta-regulatory programs were augmented from the initial 2500 non-overlapping peaks by extending each sequence by 100 bases on each side, using a stride length of 50 bp to generate partially overlapping sequences. The augmented set was then divided into train-validate-test sets in an 80:10:10 proportion. The CNN model used default parameters except for *seq_len=500* and *first_kernel_size=30*. Non-default parameters for CNN+LSTM and CNN+LSTM* models were *seq_len=500, max_pool_size=16, max_pool_stride=8, lstm_out=512, lstm_do=0.3, pre_dense_do=0.3* in addition to motif initialization for CNN+LSTM*. All models used default configuration for '*topic_classficiation*'. Models were trained on GPU (Nvidia RTX 2080 10.7G) using for a maximum of 100 epochs with early stopping. The best model with the least cross entropy loss in the validation set was selected for each approach. The obtained models' performance was evaluated on the test cohort, based on average auROC (*keras.metrics.AUC(curve='ROC'), tensorflow*, *Python*), auPR (*keras.metrics.AUC(curve='PR'), tensorflow)* and F1 score (*f1_score(), sklearn.metric*s) values. Model performance was further evaluated using cluster specific marker peaks obtained from Mannens *et al*.[19]. Prediction thresholds for each meta-regulatory program class per model were obtained by classifying the original 2500 peaks per meta-regulatory program and obtaining the prediction cut-off that maximized F1 scores per meta-regulatory programs. Marker peak sets were then assigned the identity of the meta-regulatory program with the highest prediction score above program-specific thresholds for a model. Peaks where the prediction score for each of the meta-regulatory programs fell below threshold were labelled as NA. Finally, for marker peaks per cluster, a proportion of assigned meta-regulatory program class was obtained as shown in Fig. S11C.

**TF-MoDIsco-derived regulatory grammar**

We used the CNN+LSTM* as trained above to obtain contribution scores for each of the 2500 peaks representing the original 57 meta-regulatory programs using *contribution_scores()* (*CREstd*) using detail parameters. We then used *CREsted's* implementation of TF-MoDIsco[46] (*modisco.tfmodisco()*) to obtain seqlets (short sequence instances important for model predictions) per meta-regulatory program. Obtained seqlets across meta-regulatory programs *modisco.process_patterns()* and *modisco.create_pattern_matrix()* resulted in 165 seqlets and a seqlet abundance matrix across all 57 meta-regulatory program peak-sets. Out of the 165 seqlets, one seqlet with less than 3 bases and eight seqlets with only negative seqlet counts (representing negative contribution scores) were discarded from further analysis, resulting in a final set of 156 seqlets or motifs. Obtained seqlets were converted into MEME format for HOMER based enrichment analysis. TOMTOM[98] from MEME Suite (https://meme-suite.org) was used to identify the best matching known TF-motifs per-seqlet in JASPAR2024_CORE_vertebrates_non-redundant_v2 database.

**Transcription factor gene-regulatory networks (TF-GRNs)**

TF-associated gene regulatory networks (TF-GRNs) were obtained per class using the *SCENIC+*[47] (*Python*) approach for non-multi-omic data. As described earlier, annotated cells in both transcript (*Transcriptomic meta-gene programs*) and chromatin accessibility atlas (*Cis-regulatory islands*) were split into two approximately equal parts. Each class and split was processed separately in the exact same manner, and the output was merged to obtain a consensus intersecting output, as described below. The analysis, described in detail below, involved obtaining TF-target adjacency lists, creating *cisTopic* objects for topic modelling and obtaining motif-enrichment dictionary, creating a *SCENIC+* object using transcriptomic and chromatin accessibility, and finally obtaining TF-GRN networks.

*TF-target adjacency list*

A union of the top 2000 HVGs, 500 highly expressed genes and TFs expressed in at least 50% of any cluster in a class population were obtained per class. The genes were subsetted to protein coding genes. For each class and split, a loom file was created using *build_loom()* (*SCopeLoomR,R*), which was used as input for *pyscenic (-m genei3)* (*Python*)[17] to obtain TF-target adjacency lists. Correlation values were added to each TF-target pair using *pyscenic add_cor* command. TF-adjacency lists from both splits of a class were merged, first by removing TF-target links with 'NA' correlation in any split, averaging importance and rho scores, and finally filtering out TF-target links below 0.03 'rho'.

*Topic modeling and motif-enrichment dictionary*

The *cisTopic* object created in '*Chromatin accessibility*

*meta-regulatory programs'* was used for topic modelling. A peak matrix was reduced to 50 topics (*run_cgs_models(), pycisTopic*), obtained topics were binarized into region sets by '*otsu'* method and selection of top 2000 regions per topic.

Candidate enhancer regions identified from topic analysis were then assessed for motif-enrichment, leading to the creation of cistromes, an object associating TFs to potential target regions. We used *run_pycisTarget()* wrapper from *SCENIC+*, along with motif-ranking, motif-score and motif-annotation provided by the Aerts Lab[47] for GRCh38 to obtain the TF-region cistromes per sample. Default settings were used for the function, with the exception of *run_without_promoters = True*. Further, only TFs that were present in the class specific selected gene sets were selected for further processing.

*Gene regulatory network identification*

We used the *SCENIC+* approach for the non-multi-omic data to identify TF-GRNs per class and split. For each class and split, we used snRNA-seq data (after converting it into *annData* object), snATAC-seq data (as *cisTopic* object) and motif-enrichment dictionary (obtained from pycisTarget) to create a *SCENIC+* object. The TF-adjacency lists described above with correlation values were also provided as input. *SCENIC+* first identified region-to-gene linkage for identified enhancers and their target genes, and then assigned TF-to-gene links by associating TFs that are enriched in the enhancers found linked to target genes. In the final step, *SCENIC+* uses region-to-gene and TF-to-gene links to identify regulons (TF-to-region-to-gene links) that are among the top ranked, based on importance scores and assigned positive or negative regulatory relationships based on the correlation between the TF and the assigned target gene. *SCENIC+* outputs a list of possible regulons with putative activation or repression relationships. For our analysis, we focused on positive TF-target interactions, represented as '+_+' in *SCENIC+*.

*Merging TF-GRNs*

TF-GRNs obtained per class and split were merged by keeping those TF-target links that appeared in both runs. TF-GRNs obtained across families were then merged together to obtain a comprehensive list of TF-GRNs obtained across classes resulting in 439 TFs associated with 1796 GRNs.

**Context-specific TF-GRNs activity**

439 TFs identified to be associated with GRNs were grouped into families. Association of a TF-family with a particular class was statistically evaluated using chi-square (*chisq.test()*) and negative binomial model (*glm.nb(), MASS, R*).

For each of the 1796 TF-GRNs, a module score was obtained across all the cells. To identify context-specific activity of a TF associated with GRNs across multiple classes, the data was first subsetted to those selected classes. A correlation matrix of TF-GRN sets of the selected TF was obtained from a Pearson correlation of module scores. Positive correlations represented similar activity, as the genes forming the individual gene sets showed similar expression trends, while zero or negative correlation represented un-correlated and thus context -specific activity.

Significance of a TF's context-specific activity was tested through comparing the module score of the comprising genes and the AUC score of associated CREs with one-sided pairwise Wilcoxon-test (*pairwiseWilcox(), scran*). The module score or AUC score associated with a particular GRN was compared to all other GRNs of the same TF in the selected classes where the TF was 'active'. An average AUC value for all the comparisons was obtained per TF-GRN per class and representative p-values were obtained using Fisher's method (*pchisq(), stats, R*). A TF-GRN's module score or AUC score was statistically enriched in a particular class if the average AUC was above 0.50 and if it passed a significance cut-off (AUC>0.51 and adjusted p value <1e-5) in more than half of the comparisons involving other GRNs of the same TF in that class.

**Upper and lower rhombic lip comparison**

Cells grouped in class GCUBC and GC constituted the upper rhombic lip. Cells belonging to cluster "Progenitors:A1", "Glioblasts:Progenitor_AES", "Neuroblasts_I:A1_AES", "Neurons_1:PonNuc_diff", "Neurons_I:PonNuc_def", and "Neurons_II:PonNuc_mat" constituted the lower rhombic lip derived pontine nuclei lineage. For TF-GRN analysis, gene set enrichment, gene expression heatmap and CRE accessibility analyses, cells grouped in class "GCUBC" constituted upper rhombic lip precursors and cells labelled as "Glioblasts:Progenitor_AES" and "Neuroblasts_I:A1_AES" constituted pontine nuclei precursors. TF-GRN were obtained for pontine precursors using *SCENIC+* as described above for per class-split. Gene set enrichment analysis was performed at the cluster level. For gene expression heatmap, pseudobulks were obtained by splitting each cluster into 50 cells max. CRE accessibility profiles for the *PAX6* locus was obtained using *getBW*().

**Tumor versus normal gene set variance analysis**

We compared the transcriptomic signatures of tumor samples to those of normal hindbrain cell clusters (as reference) based on a combination of Pearson correlation estimates and Gene Set Variance Analysis (GSVA22 package v1.28).

**DeepHB prediction of single-nucleotide polymorphism harboring CREs**

Genomic location of 83 single-nucleotide polymorphism (SNP) sites statistically associated with Alzheimer's heritability were obtained from *Bellenguez et al.*[76]. Each SNP was represented by 5 regions of 500 bases length with the SNP at the $150^{th}$, $200^{th}$, $250^{th}$, $300^{th}$ or $350^{th}$ base, totaling in 415 candidate regions that were classified as one of the meta-regulatory programs using the CNN+LSTM* version of the DeepHB classifier. Each peak was assigned the best matching meta-regulatory program identity as described above (*DeepHB classifier*). As each SNP was represented by 5 peaks, a SNP was assigned to either the most-frequently classified class or the class with the highest prediction score, and the peak region with highest prediction score for the assigned class was selected as the representative enhancer peak harboring the SNP. Contribution scores for the representative peak with respect to the assigned meta-regulatory program was obtained using *contribution_scores()* and visualized using *patterns.contribution_scores()* (*CREsted*)

**Tumor snRNA-seq data integration**

snRNA-seq data per sample was pre-processed as above (snRNA-seq data processing and data integration). A joint NMF embedding was obtained per tumor type using cosine normalized log-counts subsetted to a union of top 1500 HVGs obtained per sample, excluding sex-chromosome specific genes. The obtained NMF embedding was used for clustering using k-nearest neighbor graph (*kneighbors_graph(n_neighbors=26, metric="cosine"), sklearn*) and leiden clustering (*la.find_partition(la.ModularityVertexPartition), leidenalg*). Integrated clusters were assigned identity based on expression of marker genes. In PA/DMG/PFA, clusters annotated as microglia, lymphocytes, endothelial or neuronal were considered 'normal'. Additionally, in PFA, clusters assigned oligodendrocyte identity were also assigned as 'normal'. In PA/DMG, clusters identified as oligodendrocytes or astrocytes were assigned 'tumor' identity and in PFA, 'tumor' cells were predominantly assigned ependymal identity.

**Tumor transcriptomic meta-gene programs**

Meta-gene programs were obtained per tumor type using an approach similar as described in *Transcriptomic meta-gene programs*. Individual samples were first factorized using LDA using ranks ranging from 3 to 15 in an increment of 3. Each latent factor was represented by the top 100 genes ranked by contribution. The Jaccard and an overlap similarity matrix was obtained for latent factor gene sets obtained from samples belonging to a tumor type. Then, a set of non-redundant and robust representative gene sets was obtained. Robustness of a gene set was evaluated by overlap with gene sets obtained for other samples, and redundancy was evaluated by overlap of gene sets obtained for the same sample across ranks. First, for a particular sample, all the latent-factor gene sets were ranked by their maximum overlap with gene sets obtained for other samples. Then, gene sets with less than 33% overlap were removed as being non-robust. Then, gene sets with less than 70% overlap with any other gene sets obtained for the samples were removed. Finally, redundancy in a sample was removed by filtering out sets with at least 33% overlap with a higher ranked gene set. Non-redundant robust gene sets obtained across samples were then clustered using *hclust*(). A final set of four, five and three representative meta-gene programs was obtained for PA, DMG and PFA, respectively.

For tumor meta-gene programs, their enrichment in tumor compartments was evaluated using module scores across cells in a tumor sample. Meta-gene programs from tumor and normal hindbrain were compared using overlap similarity.

**Tumor versus normal differential expression gene analysis**

For cellular composition analysis of PA and DMG, three broad tumor cellular compartments were identified as OPC (composed predominantly of proliferating oligodendroglia), Astroglia (astrocyte-like cells) and microglia (microglia/macrophages). Cells belonging to other cell types were discarded. Differentially expressed genes were obtained using *glm_gp()* (*glmGamPoi*[99]*, R*). Tumor OPCs were compared to clusters of PreO class, Astroglia were compared to cells belonging to clusters Astroblast (Glioblasts), Astrocytes_SLC7A10_maturing, Astrocytes_SLC7A10 and Astrocytes_ADAMTSL3 (MixedGlia), and Microglia were compared to cells belonging to Macrophage, Microglia_EGR1, Microglia_ERC2 and Microglia_PLP1 (Immune). A cut-off of log2 fold-change of 2 and adjusted p-value $< 1e-5$ was used to obtain significantly differentially expressed genes per sample per cellular compartment. Genes differentially expressed in more than half of the samples (2/3 PA and 3/5 DMG) were kept for gene ontology (GO:BP) and reactome pathway analysis using *gprofiler2*.

**Gene communities in tumor and normal hindbrain**

A modified version of the approach in Jassim *et al.*[71] to identify gene communities was used. Correlation of Pearson residuals was used as input to obtain undirected weighted graphs for both the tumors (per sample) and normal reference.

Tumor cells were divided into three compartments (OPC, Astroglia, Microglia) as mentioned above. Cells were downsampled to a maximum of 3000 cells per compartment and an expression matrix was subsetted to protein coding genes expressed in at least 5% of cells belonging to any cell type. A Pearson residual matrix was obtained for the subsetted expression matrix using *SCTransform()* (*Seurat*).

Equivalent cellular compartments were obtained from the normal hindbrain atlas by sub-setting for the following clusters: OPC_PrN, OPC_PNAD, NFOL, COP (from PreO) and OPC (from MixedGlia) representing an OPC equivalent; Astrocytes_SLC7A10_maturing, Astrocytes_SLC7A10 and Astrocytes_ADAMTSL3 (MixedGlia) representing Astroglia; MFOL_early, MFOL_late, MOLR, MOLO (MOG) and MFOL_late (MixedGlia) representing mature oligodendrocytes; Progenitors and Microglia_early (Immune) comprising Microglia_early; and Macrophage, Microglia_EGR1, Microglia_ERC2 and Microglia_PLP1 (Immune) constituting Microglia. Cells were then downsampled to a maximum of randomly selected 3000 cells per cluster and an expression matrix was subsetted to protein coding genes expressed in at least 5% of any cluster. To improve robustness, three such normal Pearson residual matrices were obtained for the normal reference.

A weighted gene-to-gene graph was derived using the gene-gene Pearson correlation matrix obtained from the residual matrix with edges linking genes (nodes) with a minimum of 0.1 coefficient and edge-weight being coefficient^6. The obtained graph was partitioned using *best_partition()* (*community*, *Python*) to obtain gene sets, and a gene set was assigned to a cellular class based on enrichment of its module score in that cellular compartment. Gene sets enriched equally in multiple cellular compartments were assigned a ‘shared’ class. The same approach was used to identify gene communities in the three normal references and individual tumor samples. To obtain a combined gene community annotation for the normal reference from the three trials, genes were assigned to the most frequent cell class obtained in the three analyses.

**Gene expression context drift analysis**

Gene-to-gene graphs obtained above for tumor samples and each of the three normal references were processed using Node2Vec[100] followed by the skip-gram objective of the Word2Vec[101] architecture as detailed in Jassim *et al.* Briefly, Node2Vec carries out biased random walks starting from each node (gene) in its neighborhood that defines its context. Hyperparameters $p$ and $q$ were set to 1, and 200 walks of length 80 were carried out for each node. These walks were used as an input for the Word2Vec model, which converts the context into semantic representations of each node in terms of vectors and identifying relationships between nodes in terms of cosine similarity of those vectors. The parameters for the Word2Vec model included *vector_size=64, window=10, min_counts_5, sg=1, negative=5 and epochs=10*.

The change in a gene’s context between normal and tumor was analyzed by aligning Word2Vec models obtained for tumors to the normal reference using Procrustes alignment between the embedding matrices of models in a pairwise manner. From the aligned tumor Word2Vec models, cosine similarity (1- cosine distance) was obtained by comparing the vectors associated with the same gene in the tumor and the normal reference. Genes that were present only in normal graphs were assigned a value of -1.1 and genes present only in tumor graphs were assigned a value of 1.2. Finally, a context-drift score was calculated for each gene in the tumor graph as a sum of four scores: the gene’s cosine-drift score (1-cosine similarity, or 1 if gene was present only in tumor), the mean of context drift scores of immediate neighbors of the gene that were also its neighbors in the normal graph, number of new neighbors of the gene normalized to total number of genes in the tumor graph and total number of unique genes in the immediate and second-hop neighborhood of the gene normalized to total genes in the tumor graph. Context drift scores obtained from comparisons of tumor graphs to three normal reference graphs were averaged to obtain a final context drift score list per sample.
